# Antiquity and fundamental processes of the antler cycle in Cervidae (Mammalia)

**DOI:** 10.1101/2020.07.17.208587

**Authors:** Gertrud E. Rössner, Loïc Costeur, Torsten M. Scheyer

## Abstract

The origins of the regenerative nature of antlers, being branched and deciduous apophyseal appendages of frontal bones of cervid artiodactyls, have long been associated with permanent evolutionary precursors. In this study, we provide novel insight into growth modes of evolutionary early antlers. We analysed a total of 34 early antlers affiliated to ten species, including the oldest known, dating from the early and middle Miocene (approx. 19 to 12 million years old) of Europe. Our findings provide empirical data from the fossil record to demonstrate that growth patterns and a regular cycle of necrosis, abscission and regeneration are consistent with data from modern antlers. The diverse histological analyses indicate that primary processes and mechanisms of the modern antler cycle were not gradually acquired during evolution, but were fundamental from the earliest record of antler evolution and, hence, explanations why deer shed antlers have to be rooted in basic histogenetic mechanisms.

## Introduction

Antlers, paired osseous outgrowths of the deer skull, were described as ‘improbable appendages’ (Goss 1983, Kierdorf et al. 2009) due to their unique, periodically repeated, cycle of growth, death and epimorphic regeneration (*de novo* formation of a lost appendage distal to the level of amputation, Li et al. 2005) *in toto.* The strongly programmed, genetic and physiological complex antler cycle outpaces any body part renewal known (Goss 1983, Bubenik 1990, Price et al. 2005, Davis et al. 2011, Kierdorf et al. 2009, Kierdorf & Kierdorf 2011, Li 2013, Li & Suttie 2012, Wang et al. 2019, Landete-Castillejos et al. 2019). In addition, antlers are so deeply integrated into socio-reproductive behaviour of cervids (deer, moose, elk, and relatives; Artiodactyla, Mammalia), the only animals developing this headgear, that the existing cervid diversity is largely a product resulting from sexual (antler) selection interacting with intrinsic as well as environmental constraints (e.g., Darwin 1871, Whitehead 1972, Clutton-Brock et al. 1980, Clutton-Brock 1982, Goss 1983, Geist 1989, Janis 1990, Samejima and Matsuoka 2020). Each species is characterized by a specific antler morphology, and in many species sexual selection has even forced up the regrowth by larger and more complex successors with every antler generation. On the other hand, physical condition and morphology of antlers is extremely sensitive to nutrition, health and social status (Landete-Castillejos, Estevez et al. 2007; Landete-Castillejos, Currey et al. 2007, Caecero et al. 2019, Cappelli et al. 2020) and, hence, serve as a mirror of life factors.

Tissue regeneration itself is a known biological phenomenon across all groups of vertebrates, mostly from wound healing abilities. Modifications in the context of self-amputation is recorded by even 300 million years old fossils, existing long before the appearance of mammals (see Fröbisch et al. 2014, Fröbisch et al. 2015, LeBlanc et al. 2018), and includes a number of very bizarre cases (e.g., Maginnis 2006, Scherz et al. 2017). These examples, however, are never comparable with the complexity, completeness, and escalation in antlers. In mammals, appendage regrowth is commonly limited to digit tips (Gardiner 2005, Han 2005), yet the exceptional case of antlers demonstrates existence of fundamental conditions developing epimorphic regeneration *in toto* in the clade.

Although antlers are bony structures (derivates of “modified endochondral ossification” sensu Banks and Newbrey 1982; as described also in Li 2013), they do not share major functions of bones of the skeletal system. Neither do they form a substrate for muscles, nor they protect internal organs, articulate with other bones or support the body. When coming into function in intraspecific combats, they are already lifeless (Currey 1979). Antlers grow from perennial, cylindrical protuberances (pedicles) of male frontal bones (on exceptions see below) (Figure 1i-j). They grow in form of longitudinal, but not straight, branched structures. The beam is the principal cylindrical element, from which side branches dichotomously split, often in regular intervals. Re-growth of antlers always happens at the full diameter of pedicles (so-called antler pushing) without noteworthy circumferential growth (subperiosteal bone apposition in appendicular long bones), and, hence, the cross-section outline of pedicles defines that of beams (Li et al. 2005, Price et al. 2005: Fig. 2). Antler generations not only increase in size and branching complexity with progressive age of the individual, but also reach maximum daily growth rates up to 27.5 mm (in *Cervus canadensis*, Goss 1970) (not considering the putatively higher maximum growth rate of the giant deer *Megaloceros giganteus* from Pleistocene times, Lister 1994); though maximum size and branching pattern is species specific (e.g., Geist 1998, Krauss et al. 2011, Caecero 2015, Heckeberg 2017a). The onset of antlerogenesis comes with sexual maturity. Antler size and complexity peak before senescence, while during the latter aberrant forms are frequent. The burr, a ring-shaped protuberance around the base (Waldo & Wislocki 1951: plate 1, plate 5 figs 38a-c; Heckeberg 2017b: fig. 1), is an indicative character of second and subsequent generation antlers. Its position, directly above the area of bone resorption (what equates to the distal end of the pedicle) prior to antler shedding has prompted conclusions assessing its presence necessarily related to antler shedding (e.g., Lartet 1839, Dawkins 1881, Rütimeyer 1881, Filhol 1890, Lydekker 1898, Matthew 1908, Macewen 1920, Hilzheimer 1922; Pocock 1923, Schlosser 1924, Zdansky 1925, Stehlin 1928, 1937, 1939, Kraglievich 1932, Teilhard de Chardin 1939, Teilhard de Chardin and Trassaert 1937, Colbert 1936, Pilgrim 1941, Simpson 1945, Thenius 1948a, Waldo & Wislocki 1951, Crusafont 1952, Young 1964, Barrette 1977, Leinders 1983; Bubenik 1990, Ginsburg & Azanza 1991, Dong 1993, Gentry 1994, Azanza and Ginsburg 1997, Azanza et al. 2011).

**Figure 1:**
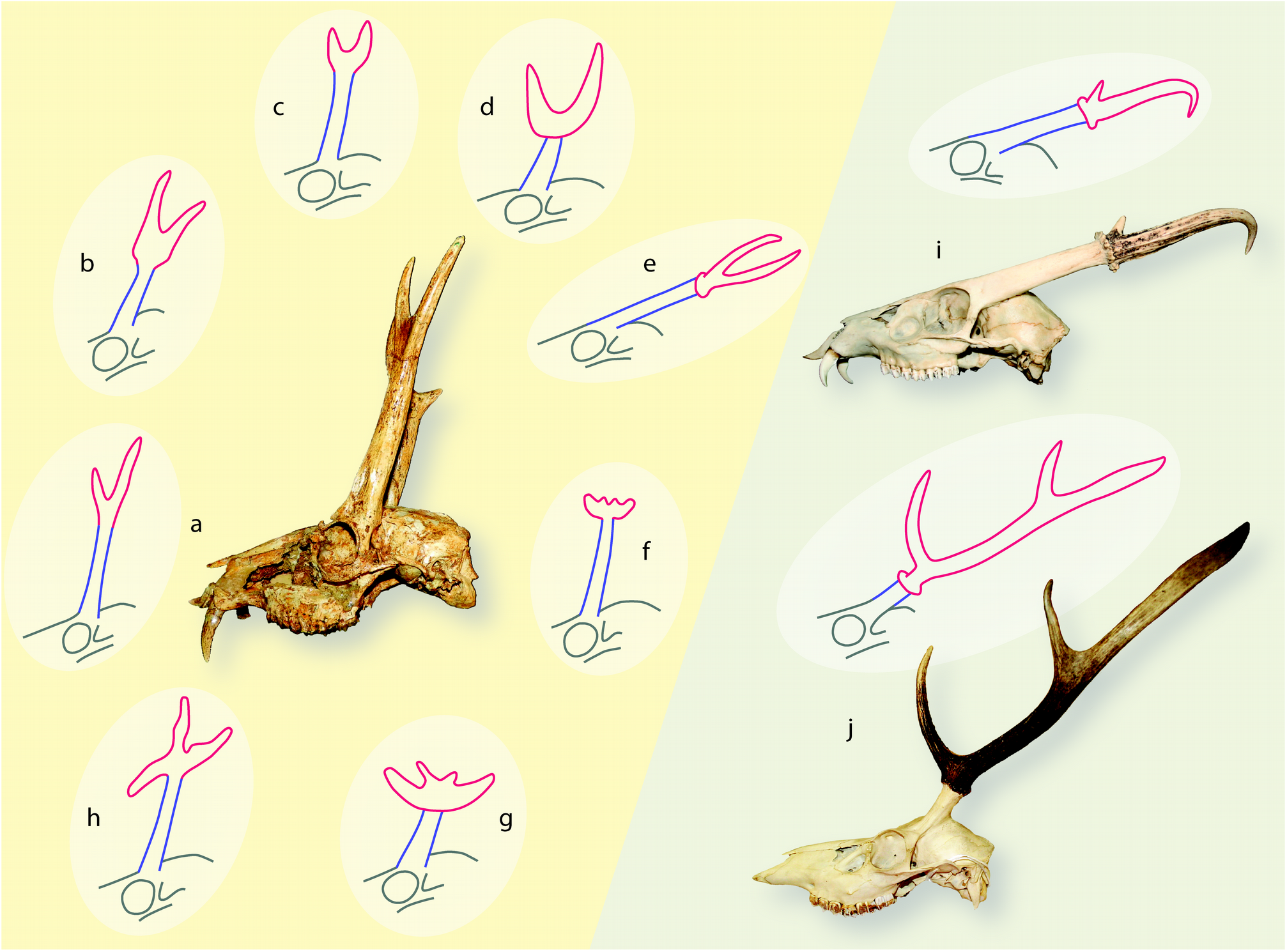
Schematic sketchs of antler morphotypes and original skulls from the Early and Middle Miocene (**a-h**) as well as extant (**i-j**) with relative antler-pedicle-proportion as well as positioning and inclination of pedicle on the skull roof. Antlers are indicated in red, pedicles in blue. Size is not to scale. **a** dichotomous geometry, *Procervulus*; left sketch, right BSPG-SNSB 1979 XV 555; **b** dichotomous geometry with basal thickening, *Heteroprox*; **c** dichotomous geometry with basal thickening, *Acteocemas*; **d** dichotomous geometry with transversal basal extension*, Dicrocerus*; **e** dichotomous geometry with burr and shaft, *Euprox*; **f** palmate geometry with transversal basal extension, *Lagomeryx*; **g** palmate geometry with transversal basal extension, *Paradicrocerus*; **h** trichotomous geometry, *Ligeromeryx*; **i** dichotomous geometry with burr, *Muntiacus muntjak*, top sketch, bottom NMB C.2023; **j** beam geometry of principal longitudinal cylindrical element from which prongs branch-off, with proximal burr and shaft (region between first basal-most split and burr), *Cervus elaphus*, top sketch, bottom NMB n.N.372.

**Figure 2:**
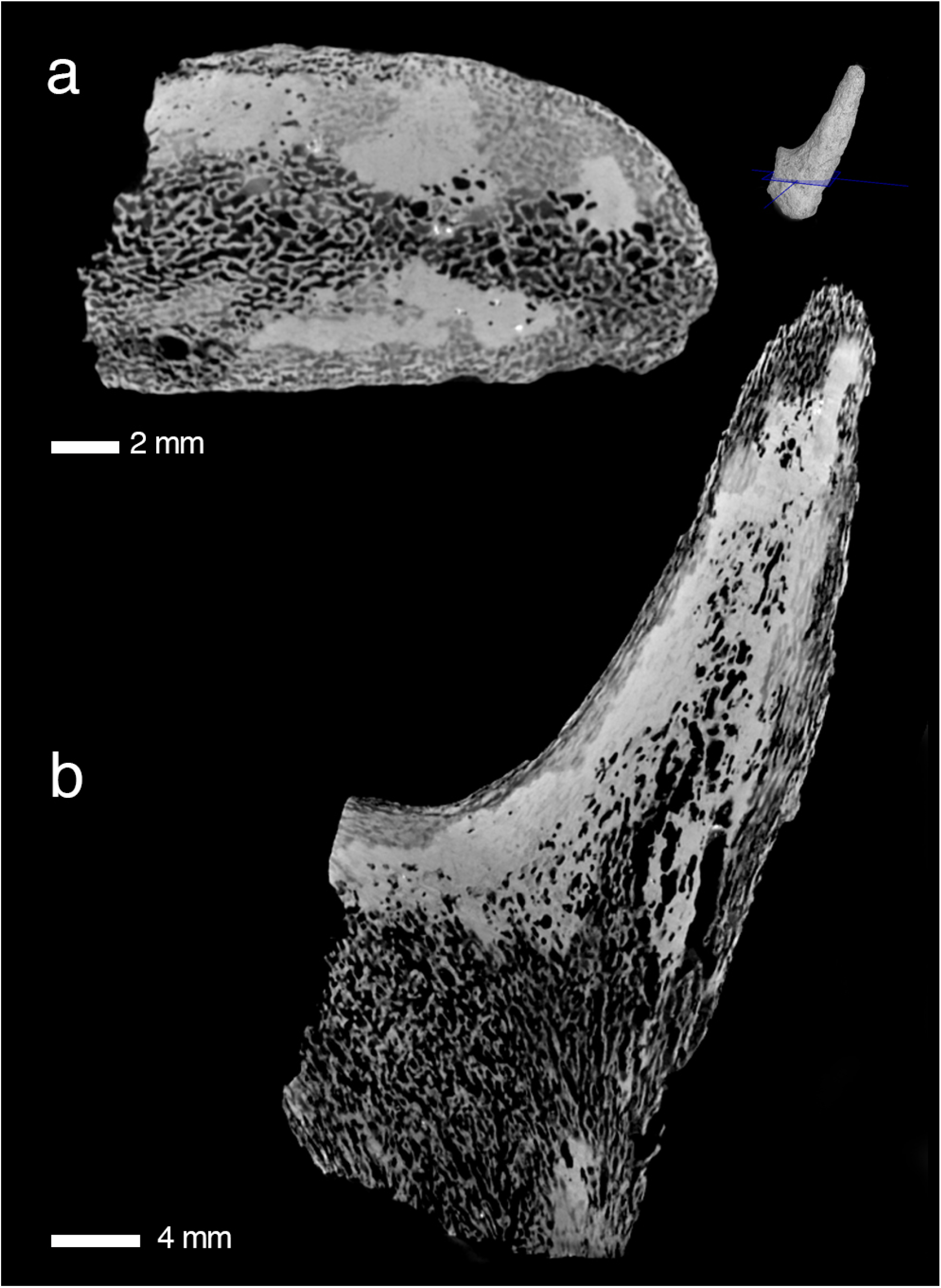
Primary bone scaffold in early antlers. An unshed antler of *Procervulus praelucidus*, SNSB-BSPG 1937 II 16810, Wintershof-West (Germany), Early Miocene (MN3), matches the initial stage of recent cortical antler bone development in Kraus et al. (2009: Fig 3d, c) and extant cancellous bone zone of proximal antler in Kierdorf et al. (2013: Fig. 8e).

The complexity of antler cycle physiology, though, is not yet fully understood (Price et al. 2005, Kierdorf et al. 2007, Kierdorf et al. 2009, Davis et al. 2011, Li and Suttie 2012, Li 2013, Wang et al. 2019). It is under intrinsic hormonal control – predominated by testosterone levels – which in turn is synchronised with extrinsic seasonality or day Iight supply: the more pronounced seasonality, the more regular antler cycle. Whereas timing of the antler cycle in the subtropical, temperate or cold zone follows a regular rhythm, tropical deer are reported to only irregularly replace their antlers (Mohr 1932, Morris 1935, Van Bemmel 1952, Ashdell 1964, Ables 1977, Loudon and Curlewis 1988, van Mourik and Stelmasiak 1990, Bubenik et al. 1991, Samsudewa and Capitan 2011, Kavčić et al. 2019 and others) up to a supposedly missing antler cycle in *Elaphodus cephalophus* (Mattioli 2011), but see Nowak (1999) and Pohle (1989), the latter described regular antler cycle in *Elaphodus cephalophus* in a German zoo under temperate climatic conditions. Other extremes are the holarctic *Rangifer tarandus* (reindeer) with antlers in both sexes (Holand et al. 2004 and references therein) and the Asian temperate *Hydropotes inermis* (water deer) which does not have antlers at all (Schilling and Rössner 2017 and references therein). The simple antler morphology in combination with an extraordinarily long pedicle rooting above the orbit in *Muntiacus* spp. and *Elaphodus cephalophus* (Figure 1i) is a striking disparity among living cervids; along with enlarged upper canines in those species, they resemble phenotypes of early times in cervid evolution (e.g. Chow and Shih 1978, Rössner 1995, Aiglstorfer et al. 2014). Small-sized antlers with simple morphology of South-American *Mazama* spp. and *Pudu* spp. are considered results from dwarfing (Eisenberg 1987). Exceptional antler-bearing females were reported from several species (Wislocki 1954; Donaldson & Doutt 1965).

The unique biology of antlers has been considered an unparalleled opportunity in order to explore processes and mechanisms of full mammalian organ regeneration (Li and Suttie 2012; Dong et al. 2019; Wang, Berg et al. 2019; Wang, Zhang et al. 2019). However, how this complex physiology has evolved over time, has received comparably little attention so far. Although there is substantial palaeontological record of antlers what allows for insights into their evolutionary history (e.g., Lartet 1839; Fraas 1862; Dawkins 1881; Rütimeyer 1881; Zittel 1893; Lydekker 1898; Zdansky 1925; Stehlin 1928, 1937, 1939; Colbert 1936; Bohlin 1937; Young 1937; Pilgrim 1941; Dehm 1944; Thenius 1948b; Young 1964; Fahlbusch 1977; Lister 1987; Azanza and Menéndez 1990; Bubenik 1990; Vislobokova 1990; Ginsburg and Azanza 1991; Azanza 1993; Dong 1993, 2008; Wang et al. 2009; Gentry 1994; Rössner 1995; Gentry et al. 1999; Azanza and Ginsburg 1997; Azanza Asensio 2000; Merino and Rossi 2010; Rössner 2010; Azanza et al. 2011, 2013; Böhme et al. 2012; Vislobokova 2013; Aiglstorfer et al. 2014; Croitor 2014; DeMiguel et al. 2014; Suraprasit et al. 2014; Hou 2015). Of particular interest are 19 to 12 million years old (early and middle Miocene) branched frontal appendages from Eurasia. They are small, have no beam structure and most of them no burr, but represent a variety of morphotypes comprising simple dichotomous, trichotomous, palmated and irregularly ramified structures (Figure 1a-h). Some of them show a more or less pronounced basal thickening proximal of the ramified distal part, some extend their bases far beyond the pedicle outline and form proximal transversal extensions with arising tines. They grew from long pedicles — instead of short ones like in most living cervids — similar in proportion with the antler length alike in living *Muntiacus* spp. and *Elaphodus cephalophus.* Unlike in modern cervids, their pedicles grew from the orbital roof upwards, causing pedicle positions directly above the eyes. This outstanding morphological disparity between cranial appendages of stem and crown cervids caught the attention of many authors and stimulated efforts in order to conclude on gradually achieved modern antler traits with consequences to Cervidae systematics. It was attempted to find homologues for morphological elements of modern antlers in early fossil antlers (burr, brow tine, shaft, beam, sculpturing). Whereas some morphotypes were recognised always as fossil homologues of their modern antler successors, others (*Procervulus*, *Lagomeryx*-related) went through odysseys of interpretations. According to the lack of a burr and a smooth surface, the latter also spurred interpretations of permanent fossil precursors of antlers (Lartet 1839, 1851; Fraas 1862; Gaudry 1878; Dawkins 1881; Rütimeyer 1881; Filhol 1890; Zittel 1893: 393; Matthew 1904; Abel 1919; Hilzheimer 1922; Colbert 1936; Bohlin 1937; Frick 1937; Stehlin 1937, 1939; Teilhard de Chardin 1939; Pilgrim 1941; Dehm 1944; Simpson 1945; Thenius 1948; Crusafont 1952; Bubenik 1966; Young 1964; Goss 1983, McFarland et al. 1985; Vislobokova et al. 1989; Bubenik 1990; Azanza 1993; Gentry 1994; Azanza and Ginsburg 1997) and as such an evolutionary pre-stage to deciduous antlers (e.g., Dong 2008, Wang et al. 2009). However, evidence for deciduousness in even these earliest antler-like organs without burrs was occasionally described during the latest decades (Ginsburg 1985; Rössner 1995; Azanza and Ginsburg 1997; Rössner 2010, Azanza et al. 2011), and only recently, a study especially dedicated to external morphological resorption traits on abscission scars in early antlers provided unequivocal evidence (Heckeberg 2017b). Thus, below we refer to ‘antlers’ when describing the respective fossils rather than to ‘antler-like organs’ (see e.g., Bubenik 1990, Azanza and Ginsburg 1997, Gentry 1994, Azanza et al. 2011, De Miguel et al. 2014). Radiographs and histological studies of some of these species and specimens enabled more differentiated insights and stimulated hypotheses on antler evolution with gradually achieved modern antler traits (Bubenik 1990, Vislobokova and Godina 1993, Rössner 1995, Azanza and Ginsburg 1997, Azanza et al. 2011). Histological features were interpreted to reflect different modes in the cycle of regeneration as compared to antlers in living cervids, including irregular shedding of still alive antlers and long-term persistence. The latter was underpinned by terms “protoantler” and “protoburr” as well as “true antlers” for modern antlers.

However, although the available histological results opened a significant new window into antler evolution, they are too sparse to provide fundamental information. A major obstacle in this respect is the destructive nature of histological methodologies as well as their limitations in the size of study objects. Since antlers constitute important (if not to say the most important) diagnostic remains of extinct deer, i.e. are frequently holotypes or included in the type series, especially those from the Miocene, invasive techniques are often not an option.

In this respect, the meanwhile established standard technique of 3D micro-computed tomography plays a critical role in overcoming the addressed problems and provides a promising new approach, not only to complement histological studies with data on internal gross microanatomy of early antlers, but also to much more easily generate data of a larger specimen sample. Accordingly, in order to substantiate the previous findings on the evolution of antlerogenesis, we here present novel research on internal antler structure using 3D micro-computed tomography. We extended specimen sampling to a much more comprehensive taxonomical coverage and were able to include type materials and rare well-preserved specimens. Most importantly, the CT scans enable us to study any section of interest (transversal, longitudinal and any in between) and therefore provide a three-dimensional understanding of the internal structuring. In addition, we used newly prepared thin-sections of a selection of specimens and species to overcome resolution deficits of the CT scans for many histological details. The new data set (1) gives insight into growth patterns in evolutionary early antlers, (2) enables comparison with modern antlers, and (3) allows for inference on evolution of underlying physiological processes. We hypothesize that all studied specimens exhibit the histological peculiarities as recognized earlier (Bubenik 1990, Azanza and Ginsburg 1997, Vislobokova and Godina 1993), and, hence, support the interpretation of a gradually acquired modern regular cycle of necrosis, abscission, and regeneration during time.

## Results

In the present study, both classical histology and non-invasive CT scanning yielded complementary data sets enabling for integrative analysis of microstructures of 34 antler and / or pedicle specimens (see Online Resource 1). Generally, histology of the fossilised antlers revealed that they are made up and shaped in their external morphology by a primary longitudinal bone scaffold of ramifying trabeculae that got filled by lamellar bone (osteons) (Figures 2-4, Online Resources 4-8,10-37). In subsequent phases of osteon formation, remaining intertrabecular, non-bone compartments got impregnated with partial replacement of the original bony framework (development of Haversian bone). In some specimens internal structure and histology show concentric differentiation with decreasing (outer cortical) lamellar bone, and reciprocally increasing woven / trabecular bone from periphery towards the centre, always restricted to regions where branching happened (Figure 4, Online Resources 21, 29, 34). Antlers with basal transversal extension and tines arising from, consist of Haversian bone only (Figures 6c-d, Online Resources 2D-F; 3E-F, K; 10-13; 23-33). With the exception of tine tips and ornamentation protuberances (Online Resources 10A-B, 12 C-D), the cortical periphery is composed of thin primary bone consisting of osteonal parallel-fibered bone (Figure 4d, f; Online Resources 4I-L, 7C-H; 12I). Unfilled erosion cavities are found in places in still attached antlers (Figures 4e). In several places of the cortical periphery, we found remains of fibres extending perpendicularly into cortical bone tissue (Figures 4d, f; Online Resources 7E, G; 13G). These fibres, which are not as prominent and coarse as in pedicles (see below) are consistent with connecting Sharpey’s fibres from the periosteum into the circumferential and interstitial lamellae of the cortical bone tissue.

In all specimens, longitudinal osteons run unidirectional from the antler’s base along morphogenetic axes. An area of wider Haversian canals are recorded from base and tine centres (Figure 3; Online Resources 14, 21, 24-26, 29, 34, 36) in still attached as well as shed antlers, but not in trichotomous antlers and palmated antlers with basal transversal extension (Figure 8; Online Resources 16, 18, 23, 30, 31, 33). Short radial Volkmann’s canals connect longitudinal vascular spaces and also meet the external surface in still attached, but also in shed specimens (Figure 3; Online Resources 10A-B, 11A-C, 12A-B, 15, 19, 21, 23, 30).

**Figure 3:**
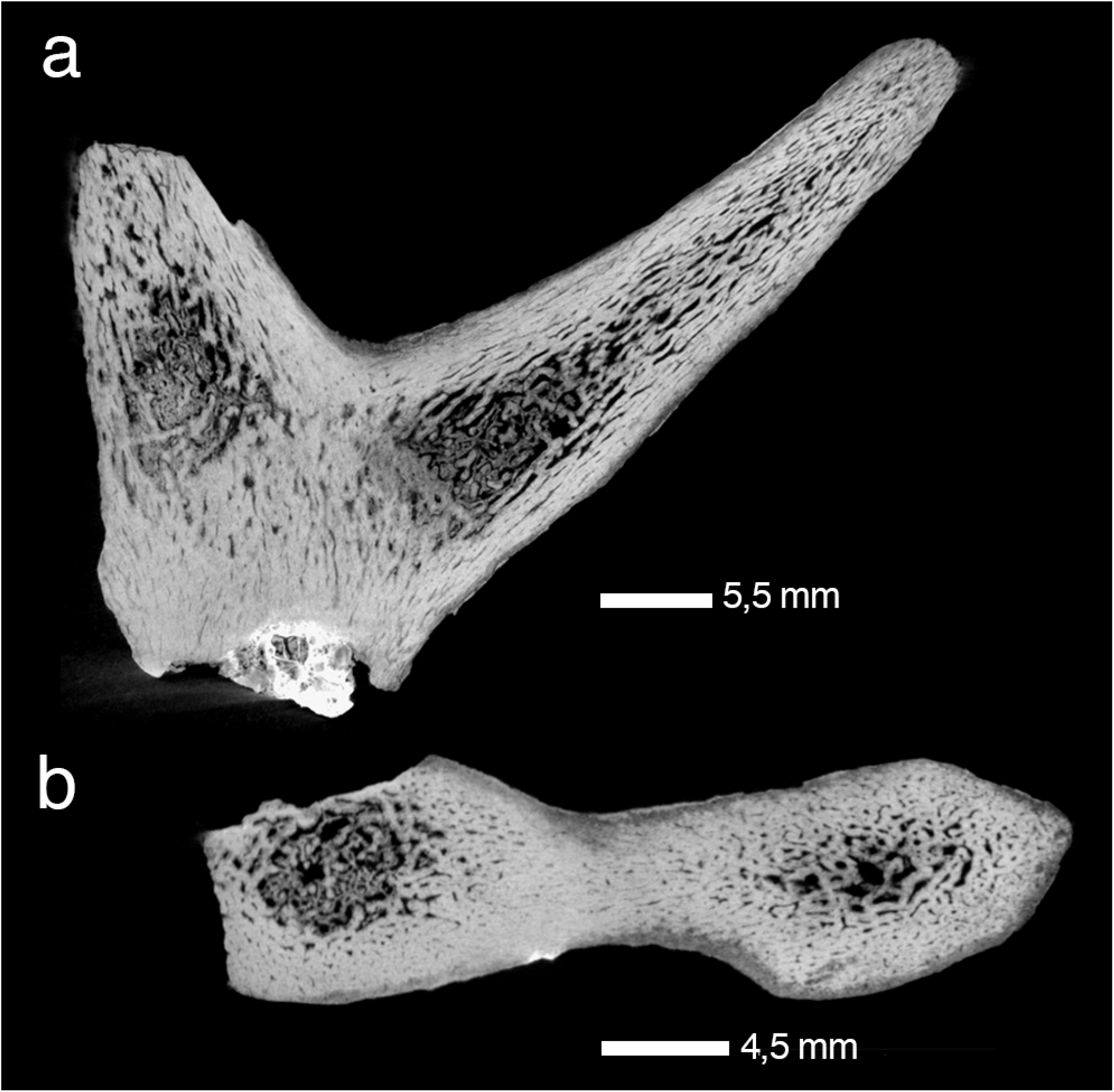
Principal tissue components in early antlers. Radiographic sections of shed antler of *Procervulus praelucidus*, SNSB-BSPG 1937 II 16842, Wintershof-West (Germany), Early Miocene (MN3), reveal predominant dense cortical bone and some trabecular bone in the tine centres as opposed to the general structure in extant antlers in Rolf and Enderle (1999: Fig. 1A, B), and trabecular bone in extant antlers (Kierdorf et al. 2013: Fig. 8e) in specific. **a** parasagittal section, **b** transversal section.

Besides remnants of primary tissue, secondary osteons are widely distributed, more extensively in the proximal base of antlers (Figure 4; Online Resources 4A-B, 12A-B, 13C-I). There are clearly less secondary osteons in tine tips of still attached antlers (Online Resources 4E-F, I-J) and ornamentation protuberances (Online Resources 12C-D). The latter exhibit even less mature bone tissue formed by woven / trabecular bone only, incompletely filled with primary osteons. Globular cell lacunae without canaliculi in the very tips of tines (Online Resources 5C-D, 11A-C) are interpreted here as chondrocyte lacunae, indicating the presence of hypertrophied remnants of cartilage.

**Figure 4:**
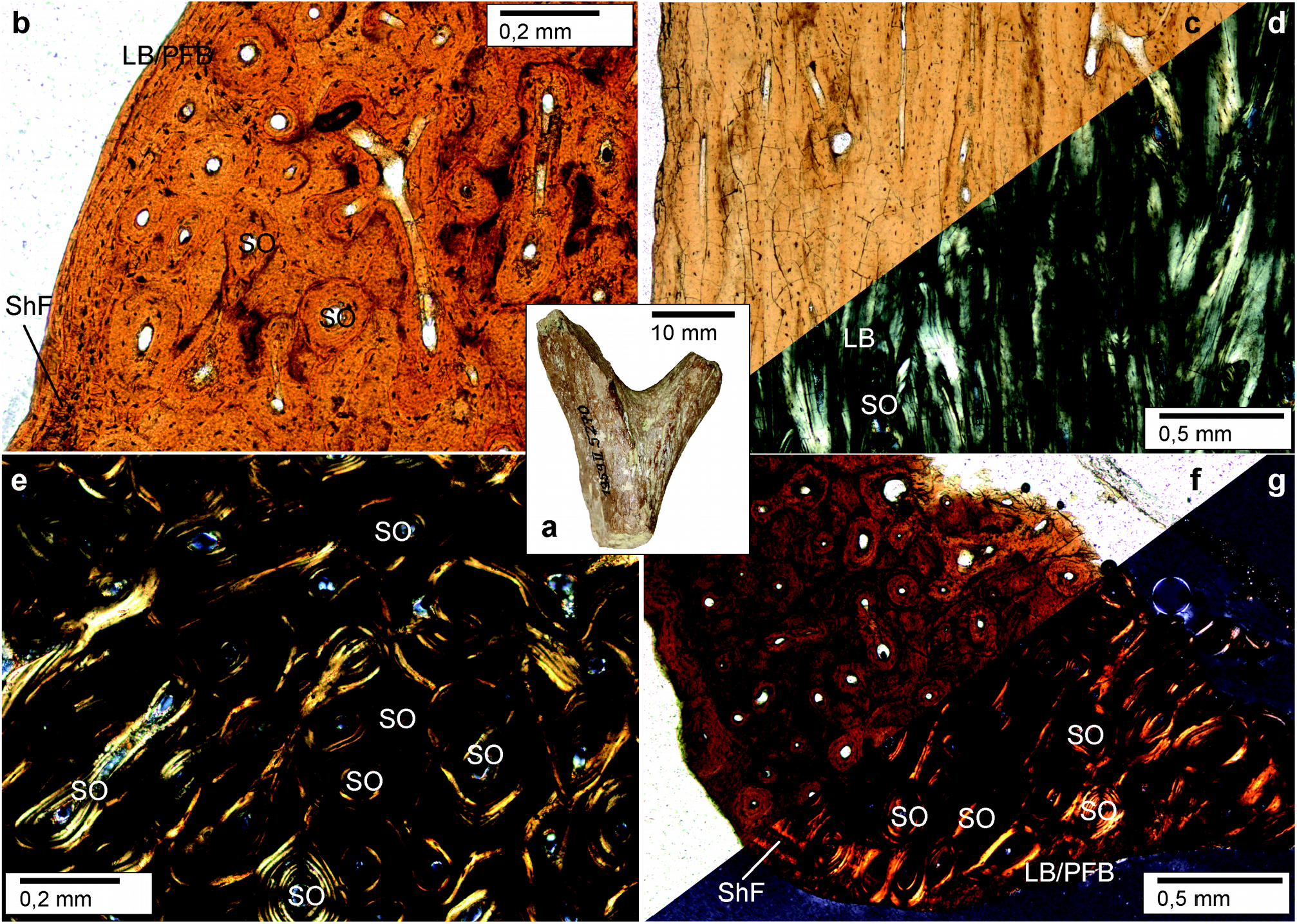
Haversion bone in early antlers. Histology of shed dichotomous antler of *Heteroprox eggeri*, SNSB-BSPG 1959 II 5270, Sandelzhausen (Germany), Middle Miocene (MN5), resembles extant proximal antler (Kierdorf et al. 2013: Fig. 7b). **a** Lateral view of specimen before sectioning, black arrowheads indicate position of thin sections taken in addition to longitudinal sections. **b, c, d, e** Close-up of the compact bone of the proximal antler. Note thin primary bone in the cortical periphery consisting of lamellar/parallel-fibred bone (well visible in **d**; note also presence of Sharpey’s fibres) and the strongly remodelled interior bone largely consisting of dense Haversian bone; **b** longitudinal section normal transmitted light, **c** longitudinal section cross-polarised light, **d** cross section normal transmitted light, **e** cross section cross-polarised light. **f**, **g** Focus on the bone tissue of the distal part of the tine. Here, most of the bone is also remodelled into dense Haversian tissue, and the external-most layer still consists of primary lamellar / parallel-fibred bone tissue, crossed by thin Sharpey’s fibres; **f** cross section normal transmitted light, **g** cross section cross-polarised light. LB, lamellar bone; PFB, parallel-fibred bone; ShF, Sharpey’s fibres; SO, secondary osteon.

Branching occurs exclusively via growth centre splitting at the distal aspect of the growing antler (Figures 2, 3, 8; Online Resources 3, 7, 10, 15-37), either dichotomous, trichotomous or palmated. Antler base thickening, ornamentation and morphotypes with bases widely extending the pedicle diameter are exclusively formed of proliferated osteonal bone (Figure 8; Online Resources 3F, 13A, 15, 31-34).

Abscission scars of shed antler specimens show enlarged spaces across osteons (Figure 5; Online Resources 2F, 3F, 13A-B, 21, 26-27) resulting from Howship’s lacunae. The latter are resorption bays caused by osteoclast activity, as can be observed in modern antlers (Kölliker 1873, Li et al. 2005, Landete-Castillejos et al. 2019).

**Figure 5:**
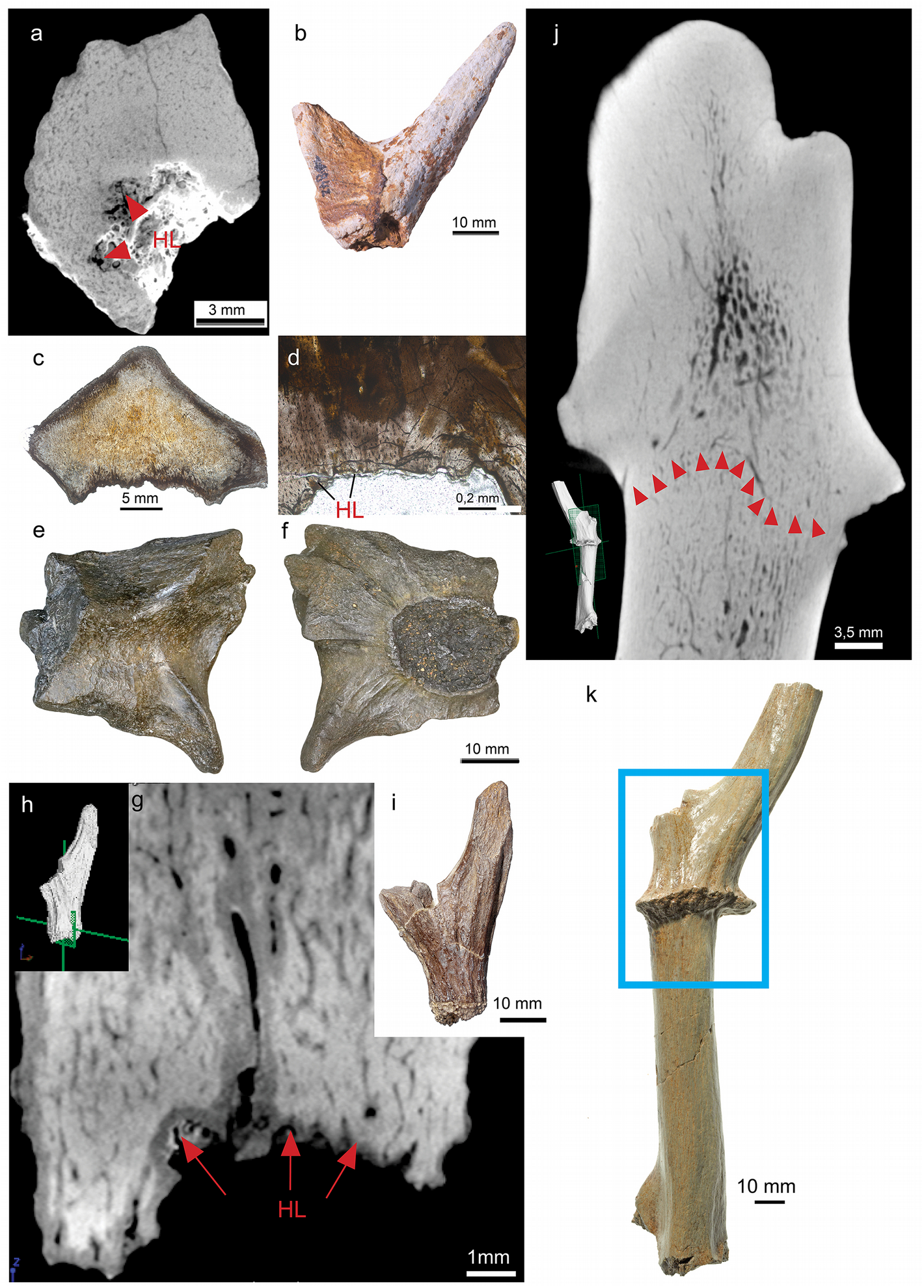
Abscission scars with Howship’s lacunae in early antlers. Shed dichotomous antler of *Procervulus praelucidus*, SNSB-BSPG 1937 II 16842, Wintershof-West (Germany), Early Miocene (MN3); **a** radiographic section through abscission scar; **b** specimen, side view. Shed palmate antler of *Paradicrocerus elegantulus*, SNSB-BSPG 1976 VI 24; **c** vertical thin section with basal abscission scar; **d** close up of abscission scar with Howship’s lacunae; **e** distal view of the specimen prior to sectioning; **f** proximal view of the specimen with roundisch abscission scar prior to sectioning. Shed dichotomous antler of *Heteroprox eggeri*, SNSB-BSPG 1959 II 5258, Sandelzhausen (Germany), Middle Miocene (MN5); **g** longitudinal radiographic section of the antler’s base; **h** placement of **g** radiographic section in the specimen; **i** side view of specimen. Still attached antler of *Euprox furcatus*, SNSB-BSPG1950 I 30, Massenhausen (Germany), Middle Miocene (MN8); **j** longitudinal radiographic section showing a fine, sub-sinus-shaped line of Howship’s lacunae directly below the burr what coincides with the junction between a pedicle and an antler in modern cervids just before antler casting (see Li 2013: Fig. 5); **k** side view of specimen, blue frame indicating region depicted in **j**. HL, Howship’s lacunae. Red arrows indicate location of HL.

The pedicles included in the study exhibit the general long bone zonation with cortical tissue, intermediate trabecular tissue, and central medullary regions (Francillon-Vieillot et al. 1990) (Figures 7-8; Online Resource 3). Cortical osteons are arranged longitudinally. Those pedicles still attached as part of the *Os frontale* evidence continuity of internal tissues between both organ regions (Figure 8; Online Resources 14-16, 18, 23-24, 28-29, 31). We note an allometric effect which links a higher portion of trabecular bone and medullary cavities with larger-sized pedicles / diameters and leaves smaller-sized pedicles without zonation (Figure 8d, j; Online Resources 15, 23, 30). Medullary regions housing larger cavities are restricted to the proximal part of the pedicle only. Haversian bone, trabecular bone, and peripheral lamellar bone of the cortex are remodelled to different grades (depending on the specimen and place) and were vascularised in a reticular pattern according to few scattered primary and predominantly secondary osteons. Radially oriented Sharpey’s fibres are frequent (and can be quite coarse) in the periphery of the pedicle cortex (Online Resources 8B-D, 9B, 10C). An early Miocene cervid skull of *Procervulus dichotomus* (SMNS 45140) with fully erupted slightly worn permanent dentition, i.e. the individual died at young adult age, has pedicles with convex distal ends consisting of fully compact bone (Online Resource 38). Convex pedicle ends match concave abscission scars of many fossil antlers (Heckeberg 2017b, Figure 5-6). However, pedicle ends do not show indication of resorption, but rather bone formation, although the length of these pedicles is shorter than that of pedicles of the same species with attached fully grown antlers and strongly worn permanent dentition (Online Resource 24). The latter might hint to substantial pedicle length increase along with the repeated shedding processes of the antler generations, what is in contrast to modern cervids in which the pedicles become shorter with each shedding process.

**Figure 6:**
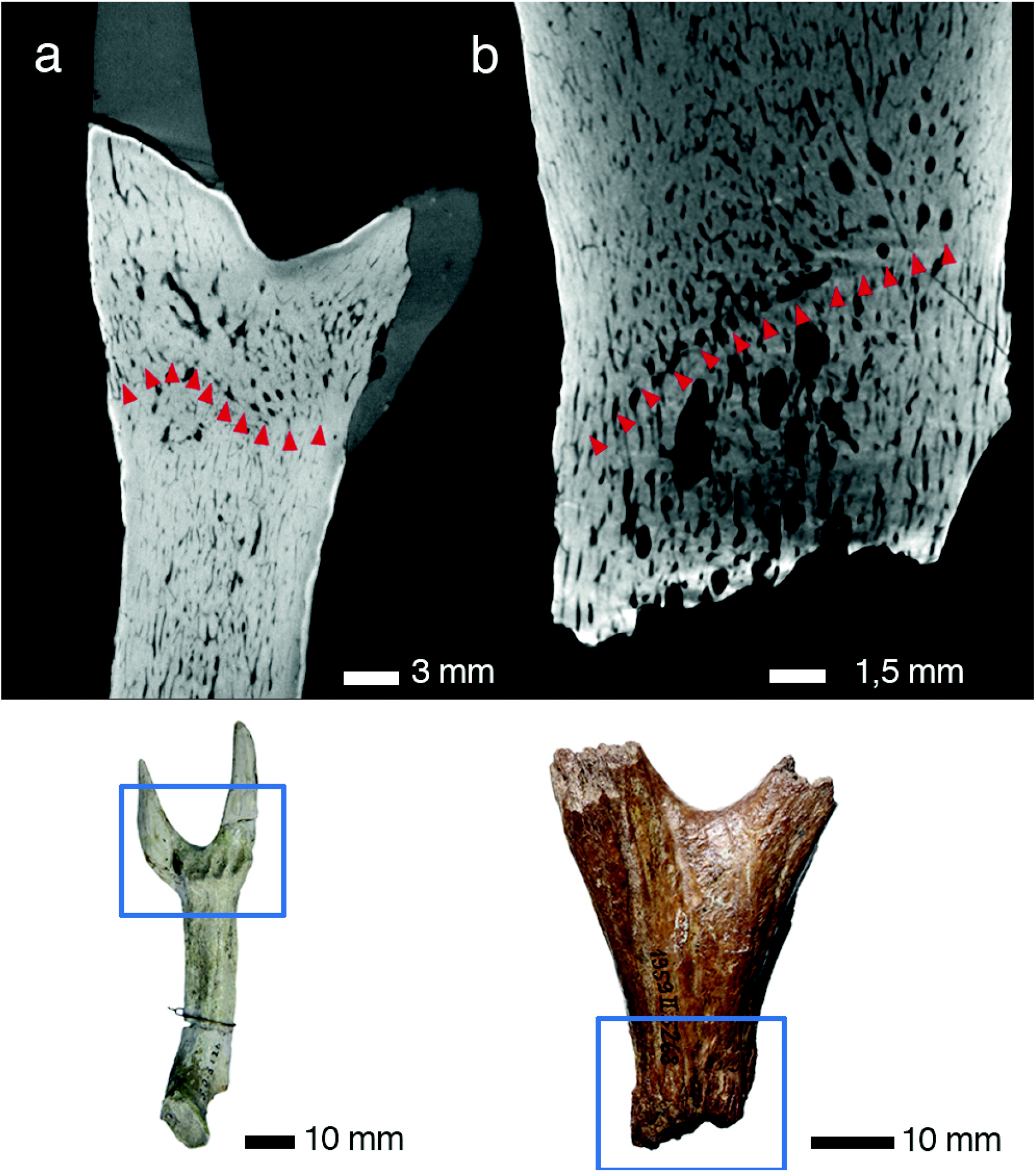
Regeneration in early antlers. At the transition from pedicle to antler two specimens show discontinuity of Haversian tissue appearing in the form of a transversal, concave towards distal, seam in a longitudinal section. It indicates not only disruption of life processes in the antler, but also reinduced growth. **a**, **b** Left attached dichotomous antler of *Acteocemas infans*, NMB S.O.3126, Chilleurs (France), Early Miocene (MN3); **a** longitudinal radiographic section, **b** lateral view of specimen. Shed dichotomous antler of *Heteroprox eggeri*, SNSB-BSPG 1956 II 5268, Sandelzhausen (Germany), Middle Miocene (MN5); **c** longitudinal radiographic section with discontinuity appearing clearly distal to the abscission scar, indicating a previous growth disruption and repeated shedding; **d** lateral view of specimen. Red arrows indicate lines of discontinuity.

**Figure 7:**
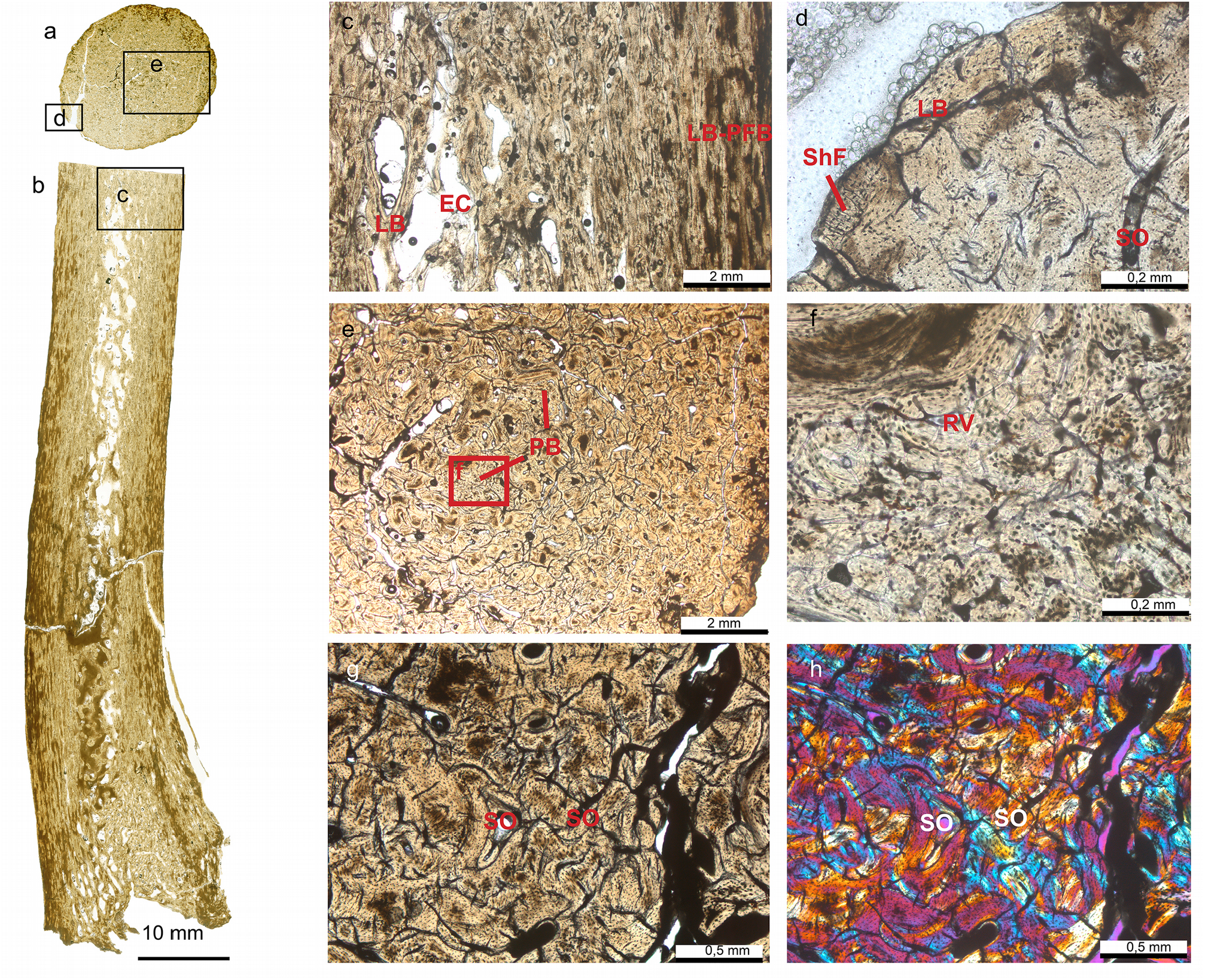
Pedicle of early antlers. Detailed histology of the pedicle of *Euprox furcatus*, NMB Sth. 12, Steinheim (Germany), Middle Miocene (MN7), in transversal (**a**) and longitudinal thin section (**b**). Images in **a-e** are in normal transmitted light, **f** in cross-polarised light using lambda compensator. **c** Close-up of distal portion of pedicle, just below the abscission area (see **b**), showing interior largely remodelled trabecular bone and a compact cortex. **d** Peripheral lamellar bone of the cortex, vascularised by few scattered primary and secondary osteons. Note presence of Sharpey’s fibres. **e** Patches of primary bone tissue with reticular vascularisation within largely remodelled Haversian bone tissue. **f** Close-up of **e**, patch of primary bone. **g**, **h** Close-up of the multiple generations of secondary osteons forming dense Haversian bone. EC, erosion cavity; LB-PFB, lamellar bone to parallel-fibred bone; PB, patches of primary bone tissue; RV, reticular vascularisation pattern; ShF, Sharpey’s fibres; SO, secondary osteon.

**Figure 8:**
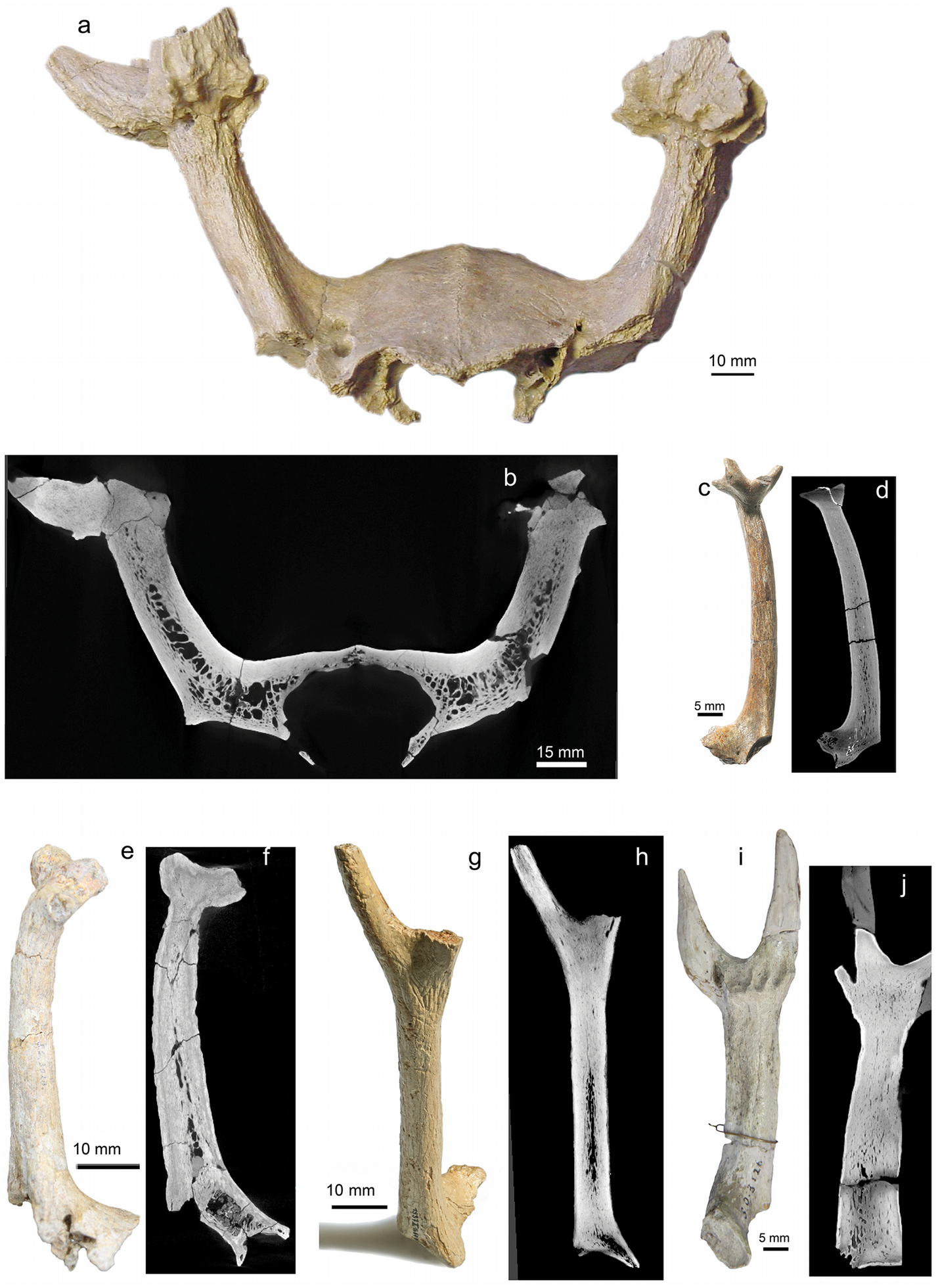
Overall internal structuring in early antlers. Radiographic sections of overall internal structuring in some examples of early antlers and associated pedicles and frontal bones. Compact Haversian bone is the most widely distributed tissue, exclusively composing the antlers (most proximal branched portion). Within pedicles trabecular bone is present in the centre relative to the overall size of the specimen (the more the larger) and there is apophyseal tissue continuity from frontal bone into pedicle base. Skull roof with both pedicles and bases of attached palmated antlers of *Paradicrocerus elegantulus*, holotype, NMA 79-5004/761, Stätzling (Germany), Middle Miocene (MN6); **a** rostral view of specimen, **b** transversal radiograph. Left palmated antler on pedicle of *Lagomeryx ruetimeyeri*, holotype of type species, SNSB-BSPG 1881 IX 55m, Reisensburg (Germany), Early Miocene? (MN4?); **c** anterior view of specimen, **d** longitudinal radiograph. Base of trichotomous antler on pedicle of *Ligeromeryx praestans*, NMB S.O. 3020, lectotype, Chitenay (France), Early Miocene (MN3); **e** anterior view of specimen, **f** longitudinal radiograph. Right dichotomous antler on pedicle of *Procervulus praelucidus*, SNSB-BSPG1937 II 16841, Wintershof-West (Germany), Early Miocene (MN3); **g** anterolateral view of specimen, **h** longitudinal radiograph. Left dichotomous antler on pedicle of *Acteocemas infans*, NMB S.O.3126, holotype, Chilleurs-aux-Bois (France), Early Miocene (MN3). **i** lateral view of specimen, **j** longitudinal radiograph. **e**, **f**, **g**, **h**, **I**, and **j** depict the oldest antlers known.

At the transition from pedicle to antler, some specimens show discontinuity of Haversian tissue appearing in the form of a transversal seam in a longitudinal section, concave towards distal (Figure 6), indicating not only disruption of life processes in the antler, but also reinduced growth. In one of these specimens, what was shed, these irregularities appear clearly distal to the abscission scar (Figure 6C) and, hence, indicate a previous growth disruption. Accordingly the specimen may document repeated shedding. The seam is in accordance with results from neohistological studies on processes during antler regeneration (Li et al. 2005: fig. 7), and as such it indicates epimorphic regeneration (*de novo* formation of a lost appendage distal to the level of amputation, Li et al. 2005).

The two shed antler specimens of living *Muntiacus muntjak* (ZSM 1966 237b) are built from compact Haversian bone only, but exhibit somewhat decreasing lamellar bone from periphery to centre. Haversian canals run through the abscission scars and Volkmann’s canals to the external surface. The burr is a result of proliferated osteonal bone, and branching into dichotomous antlers happened through growth centre bifurcation directly above the antler’s base (Online Resource 37).

## Discussion

The complexity of the modern antler cycle comprising periodic growth, necrosis, and abscission challenges several fields of biological sciences. It demonstrates principal capabilities of not only *in toto* organ replacement, but *in toto* apparatus replacement in mammals. However, the not yet fully understood outstanding biology of antlers opens to another dimension when including the fossil record. Results from morphological and histological comparisons of antlers of extinct and extant deer have been considered to represent critical differences which document the stepwise modification of ‘antler-like organs’ towards the highly derived antler biology of the modern world. Hence, the available evidence from the deep past triggered several hypotheses on the evolutionary history of antlers and initial stages, favouring permanent initial organs and / or gradually acquired modern antler characteristics including the antler cycle process (e.g., Lartet 1839, Brooke 1878, Dawkins 1881, Rütimeyer 1881, Filhol 1890, Lydekker 1898, Schlosser 1924, Zdansky 1925, Kraglievich 1932, Frick 1937, Matthew 1908, Pocock 1923, Stehlin 1928, 1937, 1939, Teilhard de Chardin 1939, Teilhard de Chardin and Trassaert 1937; Colbert 1936, Pilgrim 1941, Cabrera 1941, Dehm 1944, Simpson 1945; Crusafont 1952; Young 1964; Barrette 1977, Leinders 1983; Bubenik 1990, Dong 1993, Gentry 1994, Azanza & Ginsburg 1997, Groves 2007, Azanza et al. 2011).

Central to these discussions is the pronounced and vastly studied annual antler cycle of holarctic cervids living in temperate and cold zones including *Alces*, *Rangifer*, *Capreolus*, *Odocoileus*, *Cervus*, *Dama*, and the extinct *Megaloceros*, while neglecting other clade representatives from warmer climates or with small, simple antlers. This may have obscured relevant physiological aspects to the understanding of fundamental antler biology and in consequence of antler evolution.

In that context, the present study provides the so far most extensive insight into hard tissue traits of fossil antlers. As such our palaeohistological findings allow for a profound interpretation of growth patterns and related physiological aspects in the onset of antler evolution. Given that both modern antler histology and development have received intensified attention during the last two decades (see Kierdorf et al. 2013, Li & Suttie 2012, Li 2013, Landete-Castillejos et al. 2019 and references therein), there is a good substrate to interpret fossil structures. In the following, we compare our results with this evidence from modern antlerogenesis, and, in addition, discuss further topics relevant to an overall valid hypothesis on antler evolution.

### Pedicles and antlers

Modern antler biology provides much evidence for the functional entity of pedicles and antlers (Kierdorf et al. 2013) and as such antlers are the ‘regenerated apices of the pedicle’ (Bubenik 1990:8). An increase in testosterone levels initiates pedicle and first antler growth which originate from proliferation and differentiation of the cellular layer cells in the antlerogenic periosteum (Goss 1983, Bubenik et al. 1991). The latter overlies the crest on frontal bones of living prepubertal cervid individuals. Pedicles develop through three ossification stages: first intramembranous ossification up to 1.0 cm in height (palpable pedicle), followed by transitional ossification between 1.0 cm and 2.5 cm in pedicle height (visible pedicle), and finally pedicle endochondral ossification to complete the rest of the pedicle tissue formation (2.5–3.0 cm in height) (Li 2013) and form the antler. This transformation in ossification is *per se* an extraordinary phenomenon, as pedicles and antlers are skull appendages and skull bones derive from intramembraneous ossification, which could otherwise not grow pedicles and antlers due to insufficient vascularisation (Ham 1969, Stockwell 1979, Banks & Newbery 1982). Li et al. (1995) identified mechanical compression of stretched skin fibres during growth as the driving force in the change of ossification mode. Moreover, cells of the pedicle periosteum were identified as antler stem cells in living cervids (Li et al. 2005). The distal pedicle periosteum, however, is different, but similar to the antler’s periosteum, in that no clear demarcation between the internal cellular layer and the external fibrous layer can be readily detected (Li 2013).

Pedicles of early evolutionary antlers are characterized by their position directly on the roof of the orbit (Figures 1, 8), a place where in modern prepubertal cervids the supraorbital process of the frontal crest is located (Li & Suttie 2012: 1B). They are mostly directed almost upright and with a considerable length equal to or largely exceeding the length of the antler itself (Figures 1, 8; Online Resources 15, 24, 29, 34). In contrast, modern pedicles grow caudal to the orbit close to the parietofrontal suture, are strongly inclined caudad or laterad, and are mostly substantially shorter in comparison to the antler’s length. Our palaeohistological findings show apophyseal tissue continuity from frontal bone into pedicle base, and, hence, propose equal histological conditions with highly vascularized endochondral ossification arising from an antlerogenic periosteum homologue on the orbital roof.

Alike long bones of the appendicular skeleton modern pedicles are composed of compact cortical bone and trabecular bone in the centre, but in contrast lack medullary cavities (Li & Suttie 1998, Rolf & Enderle 1999: 2C, Kierdorf et al. 2013). The studied fossil pedicles coincide with the exception of present medullary cavities in their more proximal portions and no internal zonal patterning at all in those with a very small diameter; the latter is interpreted to be associated with allometry (Figure 7-8). Recorded frequent strong Sharpey’s fibres in external cortex tissue of stem cervid pedicles (Online Resources 8B-D, 9B, 10C) coincide with anatomy in modern pedicles (Li et al. 1995, Fig. 2C; Li 2013: figs 5C and 5D; Kierdorf et al. 2013: figs 4a, e, f). Kierdorf et al. (2013) documented evidence of extensive pedicle histogenetic remodelling in the context of the modern antler cycle. Our palaeohistological results confirm extensive tissue remodelling during the lifetime of ancient pedicles.

### Antler bone and growth

Histology of modern antlers resembles long bone tissue of the appendicular skeleton by following general principles in endochondral ossification: successive forming and replacing of preosseous (circumferential periosteum / perichondrium, cartilage, osseocartilaginous tissue) and osseous tissues (Nickel et al. 1992). Generally, osteogenesis of antlers can be equated with endochondral bone formation as in growing skeletal epiphyses (Wislocki et al. 1947), yet a contiguous growth plate and secondary ossification centres of the latter are not present (Gruber 1937, Banks 1974, Banks and Newbrey 1983).

Alike long bones, antlers are composed of compact cortical bone and trabecular bone in the centre, but in contrast lack medullary cavities and extensive circumferential growth (e.g., Chapman 1975, Bubenik 1990, Rolf and Enderle 1999, Kraus et al. 2009, Sridevu et al. 2014, Landete-Castillejos et al. 2019: fig. 8). Internal structure and histology of mature modern antler beams typically show concentrically organised bone differentiation with decreasing lamellar bone versus trabecular bone towards the centre. The shape-providing cortex is composed of dense lamellar bone, whereas the central spongiosa is built from trabecular bone scaffold only, i.e. increase of spacious bone tissue towards the centre. The inner cortex of mature Haversian bone differs from the very thin outer cortex (subvelvet zone) of incomplete primary osteons. The transitional zone between cortex and spongiosa often appears to be the zone with the greatest portion (Wislocki 1947, Chapman 1975, Bubenik 1990, Rolf and Enderle 1999, Price et al. 2005, Chen et al. 2009, Kraus et al. 2009, Kierdorf et al. 2013, Sridevu et al. 2014,). Beam tips and tines are entirely formed of compact Haversian bone (Chen et al. 2009, Kierdorf et al. 2013). However, cortical thickness and bone density appear to be dependent on species and antler size (Chapman 1975, Acharjyo and Bubenik 1983, Kierdorf et al. 2013). The interior of the small antlers of living *Muntiacus* resembles beam tips and tines of larger antlers, consisting of Haversian bone with only little transitional zone in the centre (Online Resource 37), Azanza et al. 2011). Differences with long bone histology and life-long persistent cranial appendages have been outlined (Rolf and Enderle 1999, Paral et al. 2007, Sridevu et al. 2014). Prior to the rut an increase in testosterone level causes intense ossification (Landete-Castillejos et al. 2019 and references therein).

Our palaeohistological data reveal osteonal / Haversian bone to be the predominant tissue in early antlers. Name-giving Haversian canals are the central structures of the secondary osteons housing vascularisation, and, hence, are considered the major morphophysiological unit (Francillon-Vieillot et al., 1990). In fossil antlers, bone differentiation into cortex and centrally decreasing lamellar bone, similar to the transitional zone in modern antlers, is restricted to areas where tines split, i.e. regions with largest dimensions or enlarged space in only some of the specimens / morphotypes (Figure 3, Online Resources 29, 34). We found no evidence at all for purely trabecular bone (spongiosa). However, shape and regional density of osteons are species specific. Basal transversal antler extensions and tines arising from them, both not present in modern antlers, never exhibit internal differentiation. In general, tine tips never hold internal differentiation (Figure 3; Online Resources 3, 5, 7, 10, 16, 23-24, 27, 29-31, 34-35) alike in modern antlers (Kierdorf et al. 2013). Thus, species specifics (morphotype and size) seem to have been relevant to concentric antler bone differentiation.

Generally, Haversian bone in the fossil antlers indicates rapidly proliferated tissue due to high vascularisation providing sufficient nutrient supply for high metabolic demands in the process of endochondral ossification. This fundamental histogenetic cascade is known from findings in antlerogenesis of living cervids (Ham 1969, Stockwell 1979, Banks & Newbrey 1982, Gomez et al. 2013, Kierdorf et al. 2013). Growth in antlers of living cervids is observed to take place via rapidly proliferating mesenchymal cells (Wislocki, 1942; Banks, 1974; Kierdorf et al. 1995a, 2007; Price et al. 1996, 2005; Szuwart et al. 1998; Colitti et al. 2005; Cegielski et al. 2009, Gomez et al. 2013) at the tips of beam and tines. These differentiate further proximally first to chondroblasts and then to osteoblasts forming a scaffold of longitudinal, ramifying trabeculae surrounding blood vessels. Thus, and according to the short life time of an antler, modern antler histology from tip to base shows a gradual change from an early ontogenetic stage to a more advanced ontogenetic stage or ossification grade (see Wislocki et al. 1947, Price et al 2005). Since the fundamental process of growth is the same at all ontogenetic stages, the mode of growth can be traced from periphery to the centre along an antler from distal to proximal (Wislocki et al. 1947, Price et al. 2005, Sridevu et al. 2014). Osteons in Haversian bone trace morphogenesis via preosseous tissue proliferation from the pedicle or antler’s base to tine tips or protrusion apexes. Our histological findings in fossils suggest a similar process. Yet, one of the unshed and dichotomously branched fossil specimens is built from primary trabecular scaffold only (Figure 2; Online Resources 19) and, hence, may represent an early stage of ossification before apposition of lamellar bone and a rapid growth.

Recently, Krauss et al. (2011) found evidence that fast longitudinal growth in antlers profit essentially from a mineralized tubular cartilage framework prior to osteon / bone formation in the cortex along the main antler axis what is unknown from long bones. This aspect is also of importance for biomechanical strength of hard antlers in intraspecific fighting, during which they are subjected to high impact loading and large bending moments (Chen et al. 2009; Currey et al. 2009; Launey et al. 2010). Our palaeohistological results similarly exhibit laminar bone matrix with a longitudinal tubular structure in tines (Figures 2-4, Online Resources 15, 16, 19, 23-24, 26-31, 35). Although the geometry of these antlers did not comprise extreme longitudinal elements, the tubular bone matrix was present, already supporting rapid growth and being of potential advantage in intraspecific combat use.

As opposed to Chapman (1975), ‘secondary osteons’ are revealed to be a substantial part of the fossil antlers interspersing the initial ossified framework. In living cervids, the formation of osteons successive to a first phase of osteon formation appears regularly during antlerogenesis (Gomez et al. 2013; Wislocki 1942, Krauss et al. 2009: fig. 1c; Krauss et al. 2011: figs 2c, 3e-f, 4; Kierdorf et al. 2013: figs 6, 7). They impregnate intertrabecular, non-bone compartments and only to a lesser extent replace the original bony framework, improving the strength of the forming antler. Depending on the author, these are termed ‘primary osteons’ (Kierdorf et al. 2013) or ‘secondary osteons’ (Krauss et al. 2011, Gomez et al. 2013). Due to the longitudinal growth of antlers and ossification from periphery to the centre, this impregnation with successive osteons is widest distributed in the most mature proximal part of the inner cortex with decreasing density towards the tine tips and exhibits a zonation of different ossification grade from proximal to distal. This is in accordance with what we found for the fossil antlers under study (Figure 4; Online Resources 4, 7, 12). Further, Kierdorf et al. (2013) delimit the term ‘secondary osteons’ to structures completely replacing previous antler bone, being formed late during antler growth, and, hence are comparably rare. Though, such a sequence of different grade osteons is hard to decipher in detail in palaeohistological sections of the fossil antlers, however, there is compelling evidence on successive addition, as well as replacement of bone tissue through widely distributed successive, non-first-phase, osteons similar to modern annually shed antlers. Some unshed specimens contain open erosion cavities in places and may represent ontogenetic tissue stages directly after resorption and prior to refilling with secondary osteons in the sense of Kierdorf et al. 2013 (Online Resources 5E, 7, 12B). Lacking ‘secondary osteons’ in ornamentation protuberances in fossil antlers may hint at their formation during late growth states and is again in line with findings from living cervids’ antlers (Bubenik 1966).

At the periphery of the fossil antlers, thin primary bone is deposited consisting of lamellar parallel-fibred bone with the exception of distal tine regions and ornamentation protuberances (Figure 4d, f; Online Resources 4I-L, 7C-H; 13I). This peripheral bone layer does not contain resorption spaces or any other secondary feature in contrast to the Haversian bone, and, thus, is more immature, i.e. was deposited during late growth. It may be a homologue of the outer cortex (subvelvet zone), a thin bony sleeve, deposited along the periphery of forming modern antlers in late growth by periosteal appositional (intramembranous) ossification and absent in tine tips (Li et al. 1995, Li and Suttie 1998, Krauss et al. 2011, Kierdorf et al. 2013, and references therein).

Apoptosis has been revealed to be an essential trigger for the rapid growth, morphogenesis, and tissue remodelling of extant antlers (Colitti et al. 2005), and there is no reason why it should not had worked the same way in the geological past when considering the fundamental consistency in fossil and extant antler histology.

The pattern of bone differentiation (outer cortex, inner cortex, transitional zone, spongiosa), across antlers of all extinct and extant cervids considered, hints at allometric scaling. As all the studied antler fossils have clearly smaller-sized dimensions than the modern antlers referred to in the latter works, the amount of trabecular portion appears to be related to antler size, i.e. the smaller the antler diameter, the less the trabecular portion. This is consistent with findings in modern antler beam tips and tines, whose diameters are smaller than the one of beams, as well as in modern *Muntiacus* (Online Resource 37), whose antlers are closer in size to the studied fossils. Yet, internal bone differentiation is not only about antler size, but also reflects growth patterns, as fossil antlers with basal transversal extensions do not exhibit zonation at all, but simple bifurcated antlers do. Similarly, the burr and entire antler base in living cervids is composed of Haversian bone only (Rolf and Enderle 2009, Li et al. 1995, Kierdorf et al. 2013). The biological advantage of incomplete ossification (spongiosa and transitional zone) in antlers is obvious: it reduces duration of antlerogenesis and lessens weight when size increases.

Burr formation represents transversal mesenchymal cell proliferation in addition to longitudinal growth at the onset of antler formation. The ring shaped protuberance around the base of crown cervid antlers is known to appear with the first regeneration of antlers (second antler generation of an individual) (see examples of primordial antlers without burrs in Stehlin 1937: fig. 8, Acharjyo and Bubenik 1983: fig. 3, Bubenik 1990: fig. 30, Heckeberg 2017b: fig. 9).

### Velvet

Alike bone formation, antler formation indicates the presence of skin, the so-called velvet. The antler velvet is a specialized skin transformed from pedicle epidermis, most likely due to a mix of chemical and mechanical induction (e.g., Li and Suttie 2000, Li 2013). Velvet is richly supplied with arteries and veins and, hence, provides the mayor nutritional source for antler formation (Wislocki 1942, Waldo et al. 1949). Unlike the skin covering the pedicle, it contains hair follicles that lack arrector pili muscles and are connected to extremely large sebaceous glands. The velvet lacks sweat glands and is thickened in comparison to pedicle epidermis. The underlying subcutaneous loose connective tissue is flattened into a thin layer, merging almost completely with the periosteum (Davis et al. 2011). (In contrast, a membrane insertion experiment demonstrated that antler regeneration could take place without pedicle skin participation, resulting in a skin-less antler, Li 2013.)

Sharpey’s fibres connect periosteum to bone, and hence, in fossil antlers, indicate former location of a matrix of connective tissue. In non-shed and shed antlers (Figures 4d, f; Online Resources 7E, G; 13G) of our study sample, Sharpey’s fibres underscore anchorage of periosteum / velvet to the antler bone alike in pedicles, although less frequent and strong than in the latter. However, we found no report on velvet liaison to bone in modern antlers for comparison.

### Necrosis, abscission

The least studied phase of the antler cycle are processes related to necrosis in the antler prior to abscission. In modern antlers longitudinal growth and mineralisation of matrix terminate with the cut of the antler’s blood supply through intensified ossification (reduction of Haversian system) caused by seasonal rise of testosterone level prior to the onset of rut (Landete-Castillejos et al. 2019 and references therein). Also, that is the time when beam and tine’ tips turn into sharp ends from rounded growth buds (Davis et al. 2009). Blood supply principally happens through arteries housed in the vascular layer of the velvet. Injection experiments evidenced that total cessation of blood circulation above the pedicle (Wislocki 1942, Waldo & Wislocki 1949) causes first death and shedding of the velvet, and then leaves the bare antler and dried-up vascularisation in the Haversian bone of the peripheral cortex (Li & Suttie 2012 and references therein). In consequence, there is necrosis of osteoblasts (Wislocki 1942) leaving dead antlerogenic tissue with vascularisation canals still opened up to the external surface. The cause of the obvious blood vessel closure and consequential cascade of velvet and antler bone necrosis is still unknown (Li and Suttie 2012). Our CT scans of shed modern *Muntiacus* antlers confirm open Volkmann’s canals (Online Resource 37). Axial canals and micro-cavities located in the antler core prompted Acharjyo and Bubenik (1983) to speculate that antlers from some deer species remained still alive through these vascular systems after velvet shedding. Also, reported blood filled vascular system and spongiosal tissue were taken as evidence of bare antlers remaining highly vascularised until just days before shedding (Rolf and Enderle 1999, Rolf et al. 2001). However, Waldo & Wislocki (1951) could not discover ‘growth, reconstruction or any sort of vitality in the bare antler. Nor supported experiments on dehydration and mechanical properties (Currey et al. 2009) the hypothesis of living bare antlers. The recorded compact cortical bone with only some wider axial canals of studied shed or mature (due to sharp tips) fossil antlers as well as open Volkmann’s canals (Figure 4, Online Resources 3, 12, 16, 17, 21, 24-29, 31, 32 34, 35) are in agreement with the above described histological data.

With a drop in circulating testosterone at the end of rutting season the consecutive antler cycle process is abscission. At the very base of modern antlers (proximal of the burr), dense osteoclast development in the trabecular bone on both sides of the future abscission scars induces the resorption process (Kölliker 1873: pl. 8 figs 94, 95; Wislocki 1942: pl. 1; Waldo and Wislocki 1951: pl. 2; Goss 1983; Bubenik 1990; Goss et al. 1992). Simultaneously, at the beam and tine tips the Haversian system is well developed with underrepresented mineralisation (Kierdorf et al. 2017). The initial thin demarcation line between pedicle and antlers is extended into erosion cavities (Howship’s lacunae) from the periphery to the centre (Wislocki 1942: pl. 1 fig. 4; Goss et al. 1992: figs 1-3; Li and Suttie 2012: fig. 3; Li 2013: fig. 5) eventually causing drop of the antler due to its weight when trabecular remainders cannot hold anymore. Extension of the resorption zone along the entire pedicle is documented (e.g., Goss et al. 1992: fig. 5, Kierdorf et al. 2013: Fig. 2).

Abscission is recorded in the fossil antlers under study by a number of obviously shed specimens. They hold evidence of resorption at the abscission scars via presence of widely distributed Howship’s lacunae (Figure 5; Online Resources 2F, 3F, 13A-B, 21, 26-27), a prerequisite preceding osteoclast activity. Volkmann’s canals meet the external surface in still attached as well as shed fossil antler specimens (Figure 3; Online Resources 10A-B, 11A-C, 12A-B, 15, 19, 21, 23, 30). This is consistent with modern antlers (see above). There is an antler still attached to the pedicle holding a fine, sub-sinus-shaped line of micro-Howship’s lacunae directly below the burr (Figures 5j-k, Online Resource 32), what coincides with the junction between a pedicle and an antler in modern cervids just before antler casting (see Li 2013: Fig. 5)

The abscission scars in modern cervids usually have a convex vaulting on the antler (Wislocki 1942, Waldo and Wislocki 1951: pls 1-2, Bubenik 1990: fig. 31, Heckeberg 2017b: figs 1A-D) and in pendant an watch-glass-like concave depression on the pedicle (e.g., Kierdorf et al. 2013: fig. 2a). This is in contrast to abscission scar geometry of early cervids, which we found to be mostly highly concavely vaulted in antlers, and a convex bulge in pedicles (Figures 5-6; Online Resources 2F, 3F, 13A-B, 21, 26-27, 32), often even not fully transversal but diagonal, and pointing to differences in the spatial distribution of osteoclastic activities. However, Bubenik (1966; 1990: Fig. 31) observed a change in the geometry of the antler abscission scars throughout the life of a red deer stag: convex from yearling up to prime-age, flat during the transitional years, and concave in older stags. The fossil record available does not allow for in detail reviewing of sets of antler generations, and, thus, we have to leave this issue for future investigations.

### Regeneration

Antler regeneration in living deer directly arises from highly organised wound healing processes within days or longer periods. Proliferation of periosteum and its derived tissue form growth centres for beam and brow tine, and were interpreted as hypertrophied scars accordingly (Goss 1972).

In the fossil antlers direct evidence of regeneration is provided by two specimens with a seam-like internal tissue inconsistency at the border between pedicle and antler across all diameter (Figure 6), reflecting non-continuous growth. One of the specimens is shed, but holds distal to its abscission region a similar seam as described above made up by assumed older (proximal) and assumed younger (distal) osteons (Figure 6c). The latter finding, led to the interpretation of a repeated abscission and that the abscission area was relocated towards proximal with every shedding process. Indeed, this is in congruence with modern phenomena where pedicles shorten over the lifetime of a cervid individual (Li 2013) and produce similar discontinuous osteonal or trabecular arrangements (Rolf and Enderele 1999: fig 2C).

However, two fossil skulls of *Procervulus dichtotomus* with antlers lacking a burr, one with an early adult dentition (complete, slightly worn) and pedicles with convex abscission scars (Online Resource 38) and one of later age (medium to heavily worn dentition) and clearly longer pedicles with attached antlers (Online Resource 24), indicate pedicle length increase with every antler regeneration. In this context, it is of interest that Kierdorf et al. (2003) reported on osteoblastic activities after abscission in modern antlers, which led to a partial restoration of the distal pedicle portion that was lost along with the shed antler. Although the portion of restoration does not exceed the portion lost with shedding, eventually resulting in pedicle reduction during the lifetime of an individual, the mere existence of this post-abscission pedicle reconstruction may represent a rudiment of an ancestral trait. It is obvious to assume that burr formation in modern antlers and pedicle elongation in fossil antlers may result from the same developmental growth stimulant. Evolutionary transitional stages may have had a more balanced pedicle loss and reconstruction as well as incipient burr formation.

### Consideration of previous studies

Earlier studies on internal organisation of stem cervid antlers (Bubenik 1990, Vislobokova et al. 1989, Vislobokova and Godina 1989, Vislobokova and Godina 1993, Azanza and Ginsburg 1997, Azanza et al. 2011) provided first insights via histological sections and conventional radiographs into these ancient organs, but they were restricted in taxonomic coverage, waiving of holotypes, and exclusively 2D imaging. Whereas Vislobokova et al. 1989 as well as Vislobokova and Godina (1989, 1993) focused on general structural differences and shared features among ruminants for systematic purposes, Bubenik (1990), Azanza and Ginsburg (1997), and Azanza et al. (2011) presented hypotheses on the evolution of antlers comprising gradual trait acquirement towards the modern antler cycle. Histological features of ancestral antlers were interpreted as results from long-term persistence and only occasional shedding / “spontaneous autotomy” as opposed to regular shedding (Bubenik 1990, Azanza 1993, Azanza and Ginsburg 1997, Azanza et al. 2011). Detected “morphostructural features” in different species were “correlated with differences in physiological processes” and interpreted to indicate “separate types of protoantlers” (Azanza and Ginsburg 1997).

In detail, relatively smooth surface and absence of a burr prompted interpretations of permanent skin-covered cranial appendages with a facultative perennial nature in *Ligeromeryx*, *Lagomeryx*, and *Procervulus* (Azanza and Ginsburg 1997, Bubenik 1990), although the authors could not provide coherent explanations. In addition, Azanza et al. (2011) suggested skin cover of shed specimens of *Procervulus* and *Heteroprox* because of the lack of a protective highly mineralized, compact zone between antler and pedicle as in modern cervids simultaneously with velvet shedding. The same was hypothesized by Bubenik (1990) for *Ligeromeryx*. Yet, Szuwart et al. (1998) clarified for living cervids that intense vascularization of the antler growth zone makes cell degeneration because of insufficient blood supply or rebuilt capillary canals respectively highly unlikely. “Sprouting”, ramification through exostoses of the cortex, was claimed for *Ligeromeryx* (Bubenik 1990, Azanza 1993, Azanza and Ginsburg 1997; mentioned also in Mennecart et al. 2016), but without histological / radiographic evidence. A highly active cortex up to post growth termination and related abscission of life-organs due to still existent velvet was suggested for *Dicrocerus* (Bubenik 1990), whereas Azanza et al. (2011) assumed velvet shedding before abscission of antlers in *Dicrocerus*, because of missing central trabecular area (i.e., loss of blood supply). Bubenik (1990) interpreted detected longitudinal central canals to the prong tips in a shed *Ligeromeryx* specimen (1990: Fig. 18B; but see our Online Resource 17 of the internal structure of the same specimen, NMB S.O. 5720, which proves the absence of central canals!) as an evidence of abscission of life-antlers due to insufficient mineralization to cut off blood supply from the pedicle, like observed with tines or distal parts of antlers in castrated deer. Azanza and Ginsburg (1997) argued that capillaries leading directly to the outer border of cast *Ligeromeryx* antlers indicate blood supply and consequently skin cover at the time of shedding. Indeed, already Wislocki (1942) observed high vascularisation penetrating from the velvet into the antler bone. However, Rolf and Enderle (1999) observed a widespread capillary system, directly after velvet shedding, throughout the four tissue zones and even extending the external border of an antler (fig. 3H). Pawlowska et al. (2014) found in a *Megaloceros* antler evidence for Volkmann’s canals after vascularisation loss as we did in *Muntiacus* (Online Resource 37). Hence, the evidence found by Azanza and Ginsburg (1997) is not in contradiction to the antler cycle in living deer, but rather shows correspondence.

In a macroevolutionary perspective, missing central spongeous bone in *Ligeromeryx* and *Dicrocerus* antlers where interpreted as immature antler bone (Azanza and Ginsburg 1997, Azanza et al. 2011). Bubenik (1990: Fig. 15.1) and Azanza et al. (2011) discussed evidence on centrifugal mineralization (from centre to the periphery) in *Dicrocerus* as opposed to the inverse ossification in modern antlers. Azanza et al. (2011: fig. 6 3d) even distinguished between a primary (external) and a secondary (deeper) cortex, not homologous with the outer and main cortex of modern antlers (Gomez et al. 2013, Kierdorf et al. 2013). Based on Chapman (1975), who considered secondary and tertiary Haversian system and interstitial lamellae to be absent from modern antlers due to their restricted life time in contrast to skeletal bones, Azanza and Ginsburg (1997) inferred from tissue remodelling in early fossil antlers on longer-lived organs with no regular shedding. The latter looks to be in accordance with lines of arrested growth (LAGs, continuous circumferential bands of the cortex caused by temporally extrinsically or intrinsically induced growth stops, see Kolb et al. 2015 and references therein) found in early antlers of *Dicrocerus elegans* (Azanza et al. 2011: fig. 6 2d). However, our reviewing of the depicted evidence led to the conclusion that these LAGs were misinterpretated circumferentially arranged osteons, grown during primary osteoblast activity during which trabeculae scaffold around blood vessels was formed, before infilling of mineralized matrix in interspaces (compare to Krauss et al. 2011: fig. 3). Indeed, in all ten extinct species and 34 specimens studied in the present paper, we neither found LAGs, nor any other feature indicating longer-lived organ duration in congruence with evidence from extant antlers (see discussion above). This is especially of interest, since LAGs have been found in long bone specimens of *Dicrocerus elegans* from the same fossil site (Amson et al. 2015) and were to be expected in antlers, in case they were longer-lived organs. On the other hand, the latter authors describe osteoporosis and cyclic bone remodelling, what indicates cyclic intervals of great demand for minerals (antler formation) in accordance with observations in modern *Odocoileus* bones (Meister 1956, Banks et al. 1968, Hillmann et al. 1973).

In addition, our results do not support centrifugal mineralization, but the general ossification pattern alike in modern antlers and provide indication for initial primary tissue only, both peripherally and centrally, that got increasingly remodelled. Obvious colour differences detected by Bubenik (1990: fig. 15.1) and Azanza et al. (2011) may represent taphonomical impregnation or alteration of the antler tissue.

### Terminological recommendations

Previous works attempted to find homologues for morphological elements of early fossil antlers in modern antlers (burr, brow antler, shaft, beam, sculpturing), and whereas some morphotypes were always recognised as fossil homologues of their modern antler successors, others went through odysseys of interpretations (e.g. *Procervulus*, *Lagomeryx*-related). Accordingly, the introduced terms “protoantler”, “true antlers”, “protoburr”, and “true burr” were meant to express morphological and physiological differences between modern antlers and their evolutionary forerunners (Bubenik 1990, Azanza & Ginsburg 1997, Geist 1998, Azanza et al. 2011, Azanza et al. 2013, Heckeberg 2017b). This retrospective terminology, however, on the one hand blurs the fundamental consistence (apophyseal, branched, deciduous organs) and uniqueness of modern and ancestral antlers. On the other hand, it simplifies the variability among modern antlers (shape, climatic dependence of antler cycle). In fact, in the geochronological perspective modern antlers appear to be highly specialised rather than “true” and their burrs are just one kind of a variety of basal extensions in antlers. Therefore, our recommendation is to avoid the terms above in favour of terms which consider the super- (e.g. antlers, basal extensions) or subordinary (e.g. modern antlers, beam-antlers, crown cervid antlers, burr, basal plate, palmation, ancestral antlers, stem cervid antlers etc.) nature in a macroevolutionary perspective.

### Discussion summary

Overall, evidence from 3D computed tomography and 2D thin-sections of early and middle Miocene antlers as well as their pedicles and comparative histology with modern homologues revealed several key aspects relevant to their evolutionary assessment. 1) Structural features of the osseous tissue reflects endochondral ossification and rapid growth. 2) Principle patterns of remodelling of the osseous tissue resemble those of annually shed antlers in living deer. 3) According to the internal bone structure and histology, there is no evidence of time recording (see Castanet 2006) alike in long bones in the studied fossil antlers. 4) Unequivocal histological evidence on abscission scars surfaces reveal resorption processes comparable to abscission in modern cervids. 5) Internal arrangement of trabeculae reflect repeated regeneration. 6) Internal zonation is dependent of place, size, and morphology, i.e. species-dependent, but independent of fundamental physiological processes of the antler cycle. 7) There is indication that relative pedicle length and burr formation / basal antler extension are histogeneticly linked: the less pedicle length, the more basal extension. For correlation of successive longitudinal growth stages in antlerogenesis of Miocene cervids with known processes in living cervids see Table 1.

**Table 1.** Correlation of successive longitudinal growth stages in antlerogenesis of Miocene cervids with known processes in living cervids based on the palaeohistological evidence.

| Growth stage | Palaeohistological evidence | Process |
| --- | --- | --- |
| Initial endochondral ossification, largely replacing the preexisting trabeculae of mineralized cartilage in the developing antler (velvet covering present) | deposition of primary and longitudinally oriented, loose bone scaffolding consisting of lamellar to woven bone with laminar to reticular vascularization | Osteoblast synthesising activity causing stabilisation of rapidly proliferating mesenchymal cells of antler tissue |
| Advanced endochondral ossification, compaction of antler bone and formation of dense cortical bone (velvet covering present) | infilling of vascular spaces in the primary bone scaffolding by primary osteonal bone (i.e., lamellar bone) forming tubular structures arranged in the longitudinal axis of the antler and differentiation between cortical and trabecular (if present) bone | Osteoblast synthesising activity causing progressive hardening and maturation of antler bone |
| Intensified ossification of antler bone at season of raised testosterone level (velvet covering present) | secondary osteons in cortical and trabecular (if present) bone as well as resorption spaces, resulting in predominantly dense tubular Haversian bone; primary bone remnants found either as interstitial pockets in more internal areas or in the cortical periphery, usually in form of a layer of lamellar bone | Osteoclast resorption activity and osteoblast synthesising activity causing bone remodelling and optimisation of antler bone strength |
| Shedding of antlers (past velvet shedding) | Howship's lacunae (resorption structures) in abscission scars at antler bases | Osteoclast resorption activity at antler-pedicle-demarcation line eventually leading to antler abscission |

## Conclusions

Our extensive exploration of internal structures of oldest fossil antlers and pedicles has produced a fairly large data set in contrast to what was known before and opens up the deepest and most detailed view into the evolutionary history of antlers we ever had. The study reveals intriguing consistence with histology in antlerogenesis of living cervids. We qualitatively investigated histology of 34 fossil antlers and comprehensive taxonomic coverage using micro-computed tomography as well as thin-sections. We compared to common knowledge on bone histology and evidence from living deer antler hard tissue. We found correspondence with histology of modern antlers recording apophyseal, rapid, longitudinal growth with growth centre splitting at branching points, progressive proximodistal centripetal ossification and remodelling, abscission as well as repeated regeneration. We found no indication of longevity, and, consequently, doubt on spontaneous autotomy, but have no reason to doubt on ephemerality alike in modern homologues. Accordingly, we cannot verify the hypothesis of a gradually acquired modern antler cycle (see Introduction), but have to conclude that characteristic physiological processes and mechanisms of the modern antler cycle, i.e. periodic cell depth, abscission, and regeneration, were fundamental to antlers with the onset of initial evolutionary stages.

Apart from the profound consistence between ancient and modern antler biology, size and shape differences correlate with differences in tissue organisation and histogenesis respectively. Lightweight constructions, with central trabecular bone, known from modern beam antlers of large cervid species (e.g., Picavet and Balligand 2016: fig. 2) are not represented among the ancient antlers studied and not in the small living *Muntiacus*. The internal structures observed point to either allometric, but also morphotype specifics. Moreover, ancient pedicle position on the orbital roof facilitated link with a medullary cavity system in the frontal bone unlike modern pedicles. Ancient pedicles’ relative length is longer than in modern pedicles and may be related to the grade of burr formation / basal extension. Geometry of ancient antler abscission scars is never convex, but concave.

Disparity in morphotype diversity characterises the difference between stem cervid antlers and crown cervid antlers. Whereas the latter are coined by the beam structure, the former hold a variety of basic morphotypes without shaft and / or beam but dichotomous, trichotomous, and palmated branching at the antler’s base exclusively (Figure 1a-h). That corresponds to the basic successive steps when it comes to establishment of novel features during evolution: variation and selection (e.g. Grant et al. 1976, Levinton 1983, Bégin & Roff 2004, Eldredge et al. 2005). An incremental integration of the beam antler in intraspecific social behaviour during rutting season (Clutton-Brock 1980, 1982) may have become essential for surviving an evolutionary bottleneck among cervids when it came to drastic Eurasian environmental and faunal turnovers during the so-called Vallesian crisis 10 million years ago in the early late Miocene (Agusti & Moyá-Solá 1990, Fortelius et al. 1996, Dong 1993, Gentry et al. 1999, Ataabadi et al. 2013, Azanza et al. 2013). In that context, the general tubular bony framework, described for modern antlers (Krauss et al. 2011) as well as for the studied fossil antlers, and a nanoscale toughening mechanism (inhomogenous fibril stretching) (Krauss et al. 2009) must have been beneficial to the rapidity of the growth process and became substantially advantageous for the evolution of a longitudinal geometry by development of a high fracture resistance.

The question why deer shed antlers, especially in face of the costly regeneration in large-sized cervids, has been answered with hypotheses on selective advantages (Whitehead 1972, Geist & Bromeley 1978). However, our results suggest that the antler cycle encompasses processes and mechanisms which are evolutionaryly and ontogeneticly deeply rooted, rather due to a common underlying developmental programm than of any functional importance (as supported by results from Metz et al. 2018). Accordingly, cervids simply have had to cope with the periodic loss and regain of their cranial appendages, and their evolutionary history was constantly accompanied by the competition between physiological costs and socio-reproductive success.

As antlers originated under totally different extrinsic conditions (ecological, faunistic, vegetational, climatic) than today (e.g. Rössner & Heissig 1999, Zachos et al. 2001, Böhme 2003, Kutzbach & Behling 2004, Pound et al. 2012, DeMiguel et al. 2013), this has to be considered when discussing the ancient antler cycle. Genetic, hormonal and photoperiodic control may be similar to modern cervids living in tropical or subtropical habitats. However, systematic studies on the irregular timing of the antler cycle of the latter species are rare (Mohr 1932, Morris 1935, Van Bemmel 1952, Ashdell 1964, Ables 1977, Loudon and Curlewis 1988, van Mourik and Stelmasiak 1990, Bubenik et al. 1991, Daud Samsudewa and Capitan 2011, Kavčić et al. 2019) and a synthesis is missing what hampers comprehensive assessment of the process in specific and in general. Yet, reports on annual antler shedding in tropical cervids living in temperate climate (Pohle 1989) and more than annual shedding in temperate cervids (Kierdorf & Kierdorf 1998) deviate from the common view.

The ultimate cause and conditions of the origination of pedicles with deciduous osseous apices remains a question to be solved. However, outcomes from histological studies exploring the origin of pedicle and antler development (Li 2013; Wang, Berg et al. 2019; and references therein) as well as principles in physiology, proteomics, and genetics controlling growth, ossification, demineralisation, and regeneration may help in solving this question (e.g., Davis et al. 2005 and references therein, Stéger et al. 2010, Hu et al. 2019; Zheng et al. 2019; Wang, Zhang et al. 2019).

## Material and Methods

34 specimens of ten species, representing either antler or pedicle or both, were selected aiming at a good taxonomic coverage over the early and middle Miocene (appr. 19 to 12 Ma) including holotypes and the oldest antlers known (*Ligeromeryx praestans* (Stehlin, 1937) from Chitenay (France), Azanza and Ginsburg 1997; *Procervulus praelucidus* (Obergfell, 1957) from Wintershof-West (Germany), Rössner 1995). We have mostly chosen fully-grown antlers to secure systematic assessment and comparability of results. Moreover, if possible, we examined multiple specimens of a species representing attached and shed specimens as well as different antler generations. There is the general issue that the fossil record does not provide series of fully-grown antlers of one individual, and hardly an entire set coming from one species. When studying geological earliest antlers, a further difficulty arises with the systematic association of not fully-grown antler morphologies. However, we intended to compensate these issues with a good sampling across systematics and ontogeny. Our investigations also considered the pedicles, as pedicle and antler form a functional entity (Li 2013). In addition, we investigated antlers of a modern *Muntiacus muntjak* to provide reference of an ancestral-type antler (long pedicle, short antler, simple branching pattern) with explorable biology. Online Resource 1 lists all specimens under study, their specifics, and applied methods.

Our methodological approach comprises high resolution X-ray computed tomography for most specimens as well as histological thin sections. Scanning was performed at the Bavarian Natural History Collections (SNSB) facilities using a phoenix|x-ray nanotom m (phoenix_x-ray, GE Sensing & Inspection Technologies GmbH, Wunstorf, Germany) and at the Biomaterials Science Center of the University of Basel (see single scanning parameters in Online Resource 1). Preparation of histological thin-sections followed standard petrographic thin-sectioning procedures as outlined by Chinsamy and Raath (1992). The antler and pedicle fossils were embedded in synthetic resin prior to cutting and polishing to prevent fracturing and loss of material, prior to being mounted on glass plates. Specimens were ground down manually to appropriate thicknesses (about 70 to 100 microns thick) using SiC powders of different grain-size (220, 500, 800) before being covered by a glass slip. The sections were then studied using a compound polarising microscope Leica DM 2500 M, equipped with a Leica DFC 420 C digital camera. Images were taken and processed using Adobe creative suite.

In order to interpret microstructural and palaeohistological findings, we applied general knowledge on bone histology from the literature (especially Francillon-Vieillot et al. 1990, Castanet et al. 1993, Castanet 2006, Kolb et al. 2015) and compared to specific results from published research on modern antlers (see below). In doing so, we often came across terminological conflicts between neontologists and palaeontologists which we tried to sort out with regard to our research question. For discussion we put our results in the context of modern antler biology (Kierdorf et al. 2013Li 2013, Li and Suttie 2012, Landete-Castillejos et al. 2019) to be able to identify fundamental traits and / or patterns.

## Abbreviations

NMA: Naturmuseum Augsburg, Germany
NMB: Naturhistorisches Museum Basel, Switzerland
SNSB-BSPG: Staatliche Naturwissenschaftliche Sammlungen Bayerns – Bayerische Staatssammlung für Paläontologie und Geologie, Munich, Germany
SNSB-ZSM: Naturwissenschaftliche Sammlungen Bayerns – Zoologische Staatssammlung München, Munich, Germany

## Acknowledgements

GER acknowledges financial support by the Deutsche Forschungsgemeinschaft (grant Ro 1197/7-1), LC by the Swiss National Science Foundation (grant200021_178853), and TMS by the Swiss National Science Foundation (grant 31003A_179401149506). We thank R. Ziegler and E. Heizmann (Staatliches Museum für Naturkunde, Stuttgart, Germany), and M. Rummel (Naturmuseum Augsburg, Germany) for access to specimens under their care and the permission for CT-scanning them. G. Schulz and B. Müller (Biomaterials Science Center, University of Basel) supported our study with CT-scans of some of the antlers. C. Kolb (Paläontologisches Institut und Museum Universität Zürich, Zurich, Switzerland) provided us with thin-sections. I. Vislobokova (Palaeontological Institute, Russian Academy of Sciences, Moscow, Russia) has helped with literature. M. Rummel, M. Schellenberger and M. Focke (both SNSB - Bayerische Staatssammlung für Paläontologie und Geologie, Munich, Germany) provided photographic and / or graphic work. C. Kolb as well as U. and H. Kierdorf (both University of Hildesheim, Germany) helped with advice on interpretation of histology and antler biology.

## Competing interests

GER declares on behalf of all authors that there are no competing interests.

## Notes

### Competing Interest Statement

The authors have declared no competing interest.

## References

Abel O (1919) Stämme der Wirbeltiere. Vereinigung wissenschaftlicher Verleger, Berlin, Leipzig

Ables ED (1977) The axis deer in Texas. Kleberg Studies in Natural Resources, Texas A & M University. ISBN-10: 0890961964, ISBN-13: 978-0890961964

Acharjyo LN, Bubenik AB (1983) The structural peculiarities of antler bone in genera *Axis*, *Rusa*, and *Rucervus*. In: Brown RD (ed) Antler development in Cervidae, Caesar Kleberg Wildlife Research Institute, Kingsville, Texas, pp 195–209. ISBN: 0-912229-04-7

Agusti J, Moyá-Solá S 1990 Mammal extinctions in the Vallesian (Upper Miocene). In: Kauffman EG, Walliser OH (eds) Extinction Events in Earth History, Lec Notes Earth Sci 30:425–432

Aiglstorfer M, Rössner GE, Böhme M (2014a) *Dorcatherium naui* and pecoran ruminants from the late Middle Miocene Gratkorn locality (Austria). Palaeobio Palaeoenviron 94:83–123. https://doi.org/10.1007/s12549-013-0141-9

Amson E, Kolb C, Scheyer TM, Sánchez-Villagra MR (2015) Growth and life history of Middle Miocene deer (Mammalia, Cervidae) based on bone histology. C R Palevol 14(8):637–645. https://doi.org/10.1016/j.crpv.2015.07.001

Asdell SA (1964) Patterns of Mammalian Reproduction. 2nd edn. Cornell University Press, Ithaca, N.Y.

Ataabadi MM, Liu L-P, Eronen JT, Bernor RL, Fortelius M (2013) Continental-Scale Patterns in Neogene Mammals Community Evolution and Biogeography: A Europe-Asia Perspective. In: Wang X, Flynn LJ, Fortelius M (eds) The Fossil mammals of Asia, Columbia University Press, New York, pp 629–655. ISBN: 9780231150125

Azanza B (1993) Sur la nature des appendices frontaux des cervidés (Artiodactyla, Mammalia) du Miocène inférieur et moyen. Remarques sur leur systématique et leur phylogénie. C R Acad Sci Paris, II, 316:717–723

Azanza B, DeMiguel D, Andres M (2011) The antler-like appendages of the primitive deer *Dicrocerus elegans*: morphology, growth cycle, ontogeny, and sexual dimorphism. Estud Geol 67(2):579–602. https://doi.org/10.3989/egeol.40559.207

Azanza B, Ginsburg L (1997) A revision of the large lagomerycid artiodactyls of Europe. Palaeont 40(2): 461–485

Azanza B, Menéndez E (1990) Los ciervos fósiles del Neógeno español. Pal Evol 23:75–82

Azanza B, Rössner GE, Ortiz-Jaureguizar E (2013) The early Turolian (Late Miocene) Cervidae (Artiodactyla, Mammalia) from the fossil site of Dorn-Dürkheim 1 (Germany) and implications on the origin of crown cervids. Palaeobiodiv Palaeoenv 93: 217–258. https://doi.org/10.1007/s12549-013-0118-8

Azanza Asensio B (2000) Los Cervidae (Artiodactyla, Mammalia) del Mioceno de las cuencas del Duero, Tajo, Calatayud-Teruel, y Levante. Mem Mus Pal Univ Zarag 8:1–376

Banks WJ (1974) The ossification process of the developing antler in the white-tailed deer (Odocoileus virginianus). Calc Tiss Res 14:257–274. https://doi.org/10.1007/BF02060300

Banks WJ, Epling GP, Kainer RA, Davis RW (1968) Antler growth and osteoporosis. I. Morphological and morphometric changes in the costal compacta during the antler growth cycle. Anat. Rec. 162:387–397. https://doi.org/10.1002/ar.1091620401

Banks WJ, Newbrey JW (1982) Antler development as a unique modification of mammalian endochondral ossification. In: Brown RD (ed) Antler Development in Cervidae, Caesar Kleberg Wildlife Research Institute, Kingsville, Texas, pp 279–306. ISBN: 0-912229-04-7

Barrette C (1977) Fighting behavior of muntjac and the evolution of antlers. Evol 31:169–176.

Bégin M, Roff DA (2004) From micro-to macroevolution through quantitative genetic variation: positive evidence from field crickets. Evol 58(10): 2287–2304. https://doi.org/10.1111/j.0014-3820.2004.tb01604.x

Böhme M (2003) The Miocene Climatic Optimum: evidence from ectothermic vertebrates of Central Europe. Palaeo Palaeo Palaeo 195:389–401. https://doi.org/10.1016/S0031-0182(03)00367-5

Böhme M, Aiglstorfer M, Uhl D, Kullmer O (2012) The antiquity of the Rhine River: stratigraphic coverage of the Dinotheriensande (Eppelsheim Formation) of the Mainz Basin (Germany). PloS ONE 7(5):e36817. https://dx.doi.org/10.1371/journal.pone.0036817

Bohlin B (1937) Eine tertiäre Säugetier-Fauna aus Tsaidam. Palaeont Sin XIV(1):1–111

Bruijn H de, Daams R, Daxner-Höck G, Fahlbusch V, Ginsburg L, Mein P, Morales J 1992 Report on the RCMNS working group on fossil mammals, Reisensburg 1990. Newsl Stratigr 26(2/3):65–118

Bubenik AB (1966) Das Geweih. Entwicklung, Aufbau und Ausformung der Geweihe und Gehörne, Paul Parey Verlag, Hamburg

Bubenik AB (1990) Epigenetical, morphological, physiological, and behavioral aspects of evolution of horns, pronghorns, and antlers. In: Bubenik GA, Bubenik AB (eds) Horns, Pronghorns, and Antlers, Springer, New York, pp 3–113. https://doi.org/10.1007/978-1-4613-8966-8, ISBN-10: 1461389682, ISBN-13: 978-1461389682

Bubenik GA, Brown RD, Schams D (1991) Antler cycle and endocrine parameters in male axis deer (*Axis axis*): seasonal levels of LH, FSH, testosterone, and prolactin and results of GnRH and ACTH challenge tests. Comp Biochem Physiol A 99(4):645–650

Caecero F, Villagrán M, Gambín-Pozo P, García AJ, Cappelli J, Ungerfeld R (2019) Better antlers when surrounded by females? The social context influence antler mineralization in pampas deer (*Ozotozeros bezoarticus*). Ethol Ecol Evol 31(4):358–368. https://doi.org/10.1080/03949370.2019.1620340

Cappelli J, Ceacero F, Landete-Castillejos T, Gallego L, Garc A (2020) Smaller does not mean worse: variation of roe deer antlers from two distant populations in their mechanical and structural properties and mineral profile. J Zool 311: 66–75. https://doi.org/10.1111/jzo.12764

Cegielski M, Izykowska I, Podhorska-Okolow M, Gworys B, Zabel M, Dziegiel P (2009) Histological studies of growing and mature antlers of red deer stags (*Cervus elaphus*). Anat Histol Embryol 38(3):184–188. https://doi.org/10.1111/j.1439-0264.2008.00906.x

Chen PY, Stokes AG, McKittrick J (2009) Comparison of the structure and mechanical properties of bovine femur bone and antler of the North American elk (*Cervus elaphus canadensis*). Acta Biomatter 5:693–706. https://doi.org/10.1016/j.actbio.2008.09.011

Chinsamy A, Raath MA (1992) Preparation of fossil bone for histological examination. Palaeont Afric 29: 39–44

Chow B, Shih M (1978) A skull of *Lagomeryx* from Middle Miocene of Linchu, Shantung. Vertebrata Palasiat 16(2):111–122

Clutton-Brock TH (1982) The functions of antlers. Behaviour 79(2-4): 108–124. https://doi.org/10.1163/156853982X00201

Clutton-Brock TH, Albon SD, Harvey PH (1980) Antlers, body size, and breeding group size in Cervidae. Nature 285:565–567. https://doi.org/10.1038/285565a0

Colbert EH (1936) Tertiary deer discovered by the American Museum Asiatic Expeditions. Amer Mus Nov 854:1–21

Colitti M, Allen SP, Price JS (2005) Programmed cell death in the regenerating deer antler. Jour Anat 207: 339–351. https://doi.org/10.1111/j.1469-7580.2005.00464.x

Crusafont M (1952) Los Jiráfidos fósiles de España. Mem Com Inst Geol, Disp Prov Barcelona 8:1–239

Currey JD (1979) Mechanical properties of bone with greatly differing functions. J. Biomech. 12:313–319. https://doi.org/10.1016/0021-9290(79)90073-3

Currey JD, Landete-Castillejos T, Estevez J, Ceacero F, Olguin A, Garcia A, Gallego L (2009) The mechanical properties of red deer antler bone when used in fighting. J Exp Biol 212:3985–3993. https://doi.org/10.1242/jeb.032292

Darwin C (1871) The descent of man and selection in relation to sex. John Murray, London.

Daud Samsudewa, Capitan SS (2011) Reproductive behaviour of timor deer (*Rusa timorensis*). Wartazoa 21(3):108–113. http://dx.doi.org/10.14334/wartazoa.v21i3.76

Davis EB, Brakora KA, Lee AH (2011) Evolution of ruminant headgear: a review. Proc R Soc B 278, 2857–2865. https://doi.org/10.1098/rspb.2011.0938

Dawkins WB (1881) On the evolution of antlers in the Ruminants. Nature 25:84–86.

Dehm R (1944) Frühe Hirschgeweihe aus dem Miocän Süddeutschlands. N Jb Min Geol Paläont, Mh 8(4): 81–98

DeMiguel D, Azanza B, Morales J (2014) Key innovations in ruminant evolution: a palaeontological perspective. Integ Zool 9(4):412–433. https://doi.org/10.1111/1749-4877.12080

Donaldson JC, Doutt JK (1965) Antlers in female white-tailed deer: a 4-year study. J Wildl Managem 29(4):699–705. https://doi.org/10.2307/3798545

Dong W (1993) The fossil record of deer in China. In: Ohtaishi N, Sheng H-l (eds) Deer of China: Biology and Management: Proceedings of the International Symposium on Deer of China Held in Shanghai, China, 21-23 November 1992. Developments in Animal & Veterinary Sciences 26:95–102. ISBN-13: 978-0444815408, ISBN-10: 0444815406

Dong W (2008) A review on morphology and evolution of antlers. In: Dong W (ed) Proceedings of the Eleventh Annual Meeting of the Chinese Society of Vertebrate Palaeontology, China Ocean Press, Beijing, pp 127–144. ISBN: 9787502770716

Dong Z, Coates D, Liu Q, Li C (2019) Quantitative proteomic analysis of deer antler stem cells as a model of mammalian organ regeneration. J Proteomics 195:98–113. https://doi.org/10.1016/j.jprot.2019.01.004

Eisenberg JF (1987) The evolutionary history of the Cervidae with special reference to the South American radiation. In: Wemmer C (ed) Biology and Management of the Cervidae, Smithsonian Institution, Washington D.C., pp 60–64. ISBN-10: 0874749816, ISBN-13: 978-0874749816

Eldredge N, Thompson JN, Brakefield PM, Gavrilets S, Jablonski D, Jackson JBC, Lenski RE, Lieberman BS, McPeek MA, Miller W III (2005) The dynamics of evolutionary stasis. Paleobiol 31(S2):133–145. https://doi.org/10.1666/0094-8373(2005)031[0133:TDOES]2.0.CO;2

Fahlbusch V (1977) Die obermiozäne Fossil-Lagerstätte Sandelzhausen 11. Ein neues Zwerghirsch-Geweih: *Lagomeryx pumilio*? Mitt Bayer Staatsslg Paläont hist Geol 17:227–233

Filhol H (1891) Études sur les mammifères fossiles de Sansan. Ann Sci Geol France XXI(1), pp 1–319

Fortelius M, Werdelin L, Andrews P, Bernor RL, Gentry A, Humphrey L, Mittmann H-W, Viranta S 1996 Provinciality, Diversity, Turnover, and Paleoecology in Land Mammal Faunas of the Later Miocene of Western Eurasia. In: Bernor RL, Fahlbusch V, Mittmann H-W (eds) The Evolution of Western Eurasian Mammals Faunas, Columbia University Press, New York, pp 449–470. ISBN-13: 978-0231082464, ISBN-10: 0231082460

Fraas O (1862) Die tertiären Hirsche von Steinheim. Württemb Naturwiss Jh 18:113–131

Francillon-Vieillot H, Buffrénil V d,Castanet J,Géraudie J, Meunier, Sire JY, Zylberberg L, Ricqlès A d (1990) Microstructure and mineralization of vertebrate skeletal tissues. In: Carter JG (ed) Skeletal Biomineralization: Patterns, Processes and Evolutionary Trends, Van Nostrand Reinhold, New York, pp 471–530. ISBN-10: 0442006209, ISBN-13: 978-0442006204

Fröbisch NB, Bickelmann C, Olori JC, Witzmann F (2015) Deep-time evolution of regeneration and preaxial polarity in tetrapod limb development. Nature 527: 231–234. https://doi.org/10.1038/nature15397

Fröbisch NB, Bickelmann C, Witzmann F. (2014) Early evolution of limb regeneration in tetrapods: evidence from a 300-million-year-old amphibian. Proc. R. Soc. B. 281:20141550. http://dx.doi.org/10.1098/rspb.2014.1550

Gardiner HM (2005) Ontogenetic Decline of Regenerative Ability and the Stimulation of Human Regeneration. Rejuv res 8(3):141–153. https://doi.org/10.1089/rej.2005.8.141

Gaudry A (1878) Les Echainements du monde animal dans les temps géologiques. Mammifères tertiaires. Paris (Savy)

Geist V (1989) Deer of the world: Their evolution, behaviour, and ecology. Stackpole Books, Mechanicsburg, PA. ISBN-10: 1840370947, ISBN-13: 978-1840370942

Geist V, Bromley PT (1978) Why deer shed antlers. Z Säugetierk 43:223–231

Gentry AW (1994) The Miocene differentiation of Old World Pecora (Mammalia). Hist Biol 7:115–158. https://doi.org/10.1080/10292389409380449

Gentry AW, Heizmann EPJ (1993) Case 2882 *Lagomeryx* Roger, 1904 (Mammalia, Artiodactyla): proposed designation of *L. ruetimeyeri* Thenius, 1948 as the type species. Bull Zool Nomencl. 50(2): 97–135.

Gentry AW, Rössner GE, Heizmann EPJ (1999) Suborder Ruminantia. In: Rössner GE, Heissig K (eds) The Miocene Land Mammals of Europe, Verlag Dr. Friedrich Pfeil, München, pp 225–258. ISBN: 978-3-931516-50-5

Gervais MP (1849) Sur la répartition des mammifères fossiles entre les différents âges tertiaires qui composent le sol de la France. C. r. hebd. Acad. Sci. 28:546–552

Giersch S (2004) Die Fauna aus den mittelmiozänen Krokodilschichten der Bohlinger Schlucht. Carolinea 62:5–50

Gilbert C, Ropiquet A, Hassanin A (2006) Mitochondrial and nuclear phylogenies of Cervidae (Mammalia, Ruminantia): Systematics, morphology, and biogeography. Mol Phylogenet Evol 40:101–117. https://doi.org/10.1016/j.ympev.2006.02.017

Ginsburg L (1985) Essai de phylogénie des Eupecora (Ruminantia, Artiodactyla, Mammalia). C R Acad Sci Paris 301:1255–1257

Ginsburg L, Azanza B (1991) Présence de bois chez les femelles du cervidé miocène *Dicrocerus elegans* et remarques sur le Problème de lòrigine du dimorphisme sexuel sur les appendices frontaux des Cervidés. C R Acad Sci Paris (II) 313:121–126

Goldfuss GA (1820) Handbuch der Zoologie, 2. Abtheilung. In: Schubert GH (ed) Handbuch der Naturgeschichte zum Gebrauch bei Vorlesungen, 3. Theil, 2. Abtheilung. Verlag Johann Leonhard Schrag, pp XXIV+510

Gomez S, Garcia AJ, Luna S, Kierdorf U, Kierdorf H, Gallego L, Landete-Castillejos T (2013) Labeling studies on cortical bone formation in the antlers of red deer (*Cervus elaphus*). Bone 52:506–515. https://doi.org/10.1016/j.bone.2012.09.015

Goss RJ (1968) Inhibition of growth and shedding of antlers by sex hormones. Nature 220:83–5. https://doi.org/10.1038/220083a0

Goss RJ (1970) Problems of antlerogenesis, Clin. Orthop. Rel. Res. 69:227–238.

Goss RJ (1972) Wound healing and antler regeneration. In: Maibach HI, Rovee DT (eds) Epidermal Wound Healing, Year Book Med. Publ., Inc., Chicago, pp. 219–228

Goss RJ (1983) Deer antlers: regeneration, function, and evolution. Academic Press, New York. ISBN-10: 0124120741, ISBN-13: 978-0124120747

Goss RJ, Praagh van A, Brewer P. (1992) The mechanism of antler casting in the fallow deer. J. Exp. Zool. 264:429–436. https://doi.org/10.1002/jez.1402640408

Grant PR, Grant BR, Smith JNM, Abbotti J, Abbotti LK (1976) Darwin’s finches: Population variation and natural selection. Proc Nat Acad Sci USA 73(1):257–261. https://doi.org/10.1073/pnas.73.1.257

Groves CP (2007) Family Cervidae. In: Prothero DR, Foss SE (eds) The evolution of artiodactyls, Johns Hopkins University Press, Baltimore, pp 249–256. ISBN: 9780801887352

Gruber GB (1937) Morphobiologische Untersuchungen am Cerviden-Geweih. Werden, Wechsel und Wesen des Rehgehörns. Nachr Ges Wiss Göttingen Math Phys Kl NF Fachgr VI Biol 3:9–63

Ham AW (1969) Histology. J.B. Lippincott Company, Philadelphia. ISBN 10: 039752062XISBN 13: 9780397520626

Han M, Yang X, Taylor G, Burdsal CA, Anderson RA, Muneoka K (2005) Limb regeneration in higher vertebrates: Developing a roadmap. Anat Rec 287B:14–24. https://doi.org/10.1002/ar.b.20082

Hartl DL, Clark AG (1997) Principles of population genetics. Sinauer Associates, Inc. Publishers; Sunderland, Massachusetts. ISBN-10: 0878933085, ISBN-13: 978-0878933082

Heckeberg NS (2017a) A Comprehensive Approach towards the Phylogeny and Evolution of Cervidae. Dissertation, Ludwig-Maximilians-Universität München. Morphobank project 1021 https://morphobank.org/index.php/Projects/Matrices/project_id/1021

Heckeberg NS (2017b) Evolution of antlerogenesis. J Morph 278:182–202. https://doi.org/10.1002/jmor.20628

Hensel R (1859) Über einen fossilen Muntjak aus Schlesien. Z Dt geol Ges 11:251–279

Hillmann JR, Davis RW, Abdelbaki YZ (1973) Cyclic Bone Remodeling in Deer. Calc Tiss Res 12:323–330. https://doi.org/10.1007/BF02013745

Hilzheimer M (1922) Über die Systematik einiger fossilen Cerviden. Centralbl Min etc 23:741–749

Holand Ø, Gjøstein H, Losvar A, Kumpula J, Smith ME, Røed KH, Nieminen M, Weladij RB (2004) Social rank in female reindeer (*Rangifer tarandus*): effects of body mass, antler size and age. J Zool 263(4):365–372. https://doi.org/10.1017/S0952836904005382

Hou S (2015) A new species of *Euprox* (Cervidae, Artiodactyla) from the upper Miocene of the Linxia Basin, Gansu Province, China, with interpretation of its paleoenvironment. Zootaxa 3911:43–62. http://dx.doi.org/10.11646/zootaxa.3911.1.2

Hu P, Wang T, Liu H, Xu J, Wang L, Zhao P, Xing X (2019) Full-length transcriptome and microRNA sequencing reveal the specific gene-regulation network of velvet antler in sika deer with extremely different velvet antler weight. Mol Gen Genom 294(2):431–443. https://doi.org/10.1007/s00438-018-1520-8

Janis CM (1990) Correlation of reproductive and digestive strategies in the evolution of cranial appendages, In: Bubenik GA, Bubenik AB (eds) Horns, Pronghorns, and Antlers, Springer, New York, pp 114–133. ISBN-10: 1461389682, ISBN-13: 978-1461389682

Kavčić K, Safner T, Rezić A, Ugarković D, Konjević D, Oršanić M, Šprem N (2019) Can antler stage represent an activity driver in axis deer *Axis axis*? Wildl Biol 2019(1):1–7. https://doi.org/10.2981/wlb.00516

Kierdorf H, Kierdorf U (1998) Wiederholte zweimalige Geweihbildung innerhalb eines Jahres bei einem freilebenden Rothirsch *(Cervus elaphus* L.). Z Jagdwiss 44:178–183. https://doi.org/10.1007/BF02250744

Kierdorf H, Kierdorf U, Szuwart T, Glemen C (1995) A light microscopic study of primary antler development in fallow deer (*Dama dama*). Ann Anat 177(6):525–532. https://doi.org/10.1016/S0940-9602(11)80085-3

Kierdorf U, Kierdorf H, Szuwart T (2007) Deer antler regeneration: cells, concepts, and controversies. J Morph 268:726–738. https://doi.org/10.1002/jmor.10546

Kierdorf U, Flohr S, Gomez S, Landete-Castillejos T, Kierdorf H (2013) The structure of pedicle and hard antler bone in the European roe deer (*Capreolus capreolus*): a light microscope and backscattered electron imaging study. J Anat 223:364–384. https://doi.org/10.1111/joa.12091

Kierdorf U, Li C, Price JS (2009) Improbable appendages: Deer antler renewal as a unique case of mammalian regeneration. Sem Cell Develop Biol 20:535–542. https://doi.org/10.1016/j.semcdb.2008.11.011

Kölliker A (1873) Die normale Resorption des Knochengewebes und ihre Bedeutung für die Entstehung der typischen Knochenformen, F.C.W. Vogel, Leipzig. ISBN 10: 375031991XISBN13: 9783750319912

Kolb C, Scheyer TM, Veitschegger K, Forasiepi AM, Amson E, Van der Geer AAE, Van den Hoek Ostende LW, Hayashi S, Sánchez-Villagra MR (2015) Mammalian bone palaeohistology: a survey and new data with emphasis on island forms. PeerJ 3:e1358. https://doi.org/10.7717/peerj.1358

Krauss S, Fratzl P, Seto J, Currey JD, Estevez JA, Funari SS, Gupta HS (2009) Inhomogeneous fibril stretching in antler starts after macroscopic yielding: Indication for a nanoscale toughening mechanism. Bone 44(6):1105–1110. https://doi.org/10.1016/j.bone.2009.02.009

Krauss S, Wagermaier W, Estevez JA, Currey JD, Fratzl P (2011) Tubular frameworks guiding orderly bone formation in the antler of the red deer (*Cervus elaphus*). J Struct Biol 175:457–464. https://doi.org/10.1016/j.jsb.2011.06.005

Kraglievich JL (1932) Contribución al conocimiento de los ciervos fósiles del Uruguay. Anal Mus Hist Nat Montevideo 2(3):355–438

Kutzbach JE, Behling P (2004) Comparison of simulated changes of climate in Asia for two scenarios: Early Miocene to present, and present to future enhanced greenhouse. Glob Planet Change 41:157–165. https://doi.org/10.1016/j.gloplacha.2004.01.015

Landete-Castillejos T, Currey JD, Estevez JA, Gaspar-López E, Garcia A, Gallego L (2007) Influence of physiological effort of growth and chemical composition on antler bone mechanical properties. Bone 41:794–803. https://doi.org/10.1016/j.bone.2007.07.013

Landete-Castillejos T, Estevez JA, Martínez A, Ceacero F, Garcia A, Gallego L (2007) Does chemical composition of antler bone reflect the physiological effort made to grow it? Bone 40:1095–1102. https://doi.org/10.1016/j.bone.2006.11.022

Landete-Castillejos T, Kierdorf H, Gomeze S, Lunae S, García AJ, Cappelli J, Pérez-Serrano M, Pérez-Barbería J, Gallegoa L, Kierdorf U (2019) Antlers - Evolution, development, structure, composition, and biomechanics of an outstanding type of bone, Bone 128:115046. https://doi.org/10.1016/j.bone.2019.115046

Lartet É (1837) Notice sur les ossements fossiles des terrains tertiaires de Simorre, de Sansan, etc., et sur la Découverte récente d’une mâchoire de singe fossile. C R hebd Acad Sci Paris 4:1–583.

Lartet, É (1839) Nouvelles espèces fossiles découvertes dans le département du Gers. C R hebd Acad Sci Paris 9: 1–66.

Lartet, É. 1851. Notice sur la colline de Sansan. Suivie d’une récapitulation de diverses espèces d’animaux vertébrés fossiles trouvés soit à Sansan, soit dans d’autres gisements du terrain tertiaire miocène dans le bassin sous-pyrénéen. Auch (Portes)

LeBlanc ARH, MacDougall MJ, Haridy Y, Scott D, Reisz RR (2018) Caudal autotomy as anti-predatory behaviour in Palaeozoic reptiles. Sci Rep 8:3328. https://doi.org/10.1038/s41598-018-21526-3

Leinders JJM (1983) Hoplitomerycidae fam. nov. (Ruminantia, Mammalia) from Neogene fissure fillings in Gargano (Italy). Part 1, the cranial osteology of *Hoplitomeryx* gen. nov. and a discussion on the classification of pecoran families. Scri Geol 70:1–68

Levinton JS (1983) Stasis in Progress: The Empirical Basis of Macroevolution. Ann Rev Ecol Syst 14:103–137

Li C (2013) Histogenetic aspects of deer antler development. Front Biosci E5(1):479–489

Li C, Suttie JM (1998) Electron Microscopic Studies of Antlerogenic Cells From Five Developmental Stages During Pedicle and Early Antler Formation in Red Deer (*Cervus elaphus*). Anat Rec 252:587–599.

Li C, Suttie JM (2000) Histological studies of pedicle skin formation and its transformation to antler velvet in red deer (*Cervus elaphus*). Anat Rec 260:62–71. https://doi.org/10.1002/1097-0185(20000901)260:1%3C62::AID-AR70%3E3.0.CO;2-4

Li C, Suttie JM (2012) Morphogenetic aspects of deer antler development. Front Biosci E4:1836–1842. doi: 10.2741/505

Li C, Suttie JM, Clark DE (2005) Histological Examination of Antler Regeneration in Red Deer (*Cervus elaphus*). Anat Rec 282A:163–174. https://doi.org/10.1002/ar.a.20148

Li C, Waldrup KA, Corson ID, Littlejohn RP, Suttie JM (1995) Histogenesis of antlerogenic tissues cultivated in diffusion chambers *in vivo* in red deer (*Cervus elaphus*). J Exp Zool 272(5):345–55. https://doi.org/10.1002/jez.1402720504

Lister AM (1987) Diversity and evolution of antler form in Quaternary deer. In: Wemmer CM (ed) Biology and management of the Cervidae, Smithsonian, Washington D.C., pp 81–98. ISBN-10: 0874749816, ISBN-13: 978-0874749816

Lister AM (1994) The evolution of the giant deer, *Megaloceros giganteus* (Blumenbach). Zool J Lin Soc 112(1-2):65–100. https://doi.org/10.1111/j.1096-3642.1994.tb00312.x

Loudon ASI, Curlewis JD (1988) Cycles of antler and testicular growth in an aseasonal tropical deer (*Axis axis*). J Repr Fert 83:729–738.

Lydekker R (1898) The deer of all lands. Ward, London.

Macewen W (1920) The growth and shedding of the antler of the deer; the Histological Phenomena and Their Relation to the Growth of Bone. Maclehose, Jackson & Co., Glasgow. ISBN-13: 978-1297624575, ISBN-10: 1297624572

Maginnis TL (2006) The costs of autotomy and regeneration in animals: a review and framework for future research. Behav Ecol 17(5):857–872. https://doi.org/10.1093/beheco/arl010

Matthew WD (1904) A complete skeleton of *Merycodus*. Bull Am Mus Nat Hist 20:101–129.

Matthew WD (1908) Osteology of *Blastomeryx* and phylogeny of American Cervidae. Bull Am Mus Nat Hist 24:535–562.

Mattioli S (2011) Family Cervidae (Deer), In: Wilson DE, Mittermaier RA (eds) Handbook of the Mammals of the World, 2, Hoofed Mammals, Lynx Ediciones, Barcelona, pp 350–428. ISBN-10: 8496553779, ISBN-13: 978-8496553774

McFarland WN, Pough FH, Cade TJ, Heiser JB (1985) Vertebrate Life, 2nd ed, MacMIllan Publishers, New York.

Meister WW (1956) Changes in histological structure of the long bones of white tailed deer during the growth of the antlers. Anat. Rec. 124:709–721. http://dx.doi.org/10.1002/ar.1091240407

Mennecart B, Rössner GE, Métais G, DeMiguel D, Schulz G, Müller B, Costeur L (2016) The petrosal bone and bony labyrinth of Early to Middle Miocene European deer (Mammalia, Cervidae) reveal their phylogeny. Journal of Morphology 277(10): 1329–1338. https://doi.org/10.1002/jmor.20579

Merino ML, Rossi RV (2010) Origin, systematics and morphological radiation. In: Duarte JMB, González S (eds), Neotropical Cervidology: Biology and medicine of Latin America deer, Switzerland: IUCN, FUNEP, Jabotical, Brasil & Gland, pp 2–11. ISBN: 9788578050467

Metz MC, Emlen DJ, Stahler DR, MacNulty DR, Smith DW, Hebblewhite M (2018) Predation shapes the evolutionary traits of cervid weapons. Nat Ecol Evol 2:1619–1625. https://doi.org/10.1038/s41559-018-0657-5

Mohr E (1932) Materialien über die Hirschzuchten des ehemaligen Hamburger Zoo. Zool. Garten 3:3–14

Morris RC (1935) Growth and shedding of antlers in sambar (*Cervus unicolor*) and cheetal (*Axis axis*) in south India. J Bombay Nat Hist Soc 37:484.

Nickel R, Schummer A, Seiferle E (1992) Lehrbuch der Anatomie der Haustiere, I (6th ed) Bewegungsapparat, Verlag Paul Parey, Berlin, Hamburg. ISBN: 9783489580164

Nickel R, Schummer A, Seiferle E (1992) Lehrbuch der Anatomie der Haustiere, III (6th ed) Kreislauf, Haut und Hautorgane, Verlag Paul Parey, Berlin, Hamburg

Nowak RM (1999) Walker’s mammals of the world. 6th ed, The Johns Hopkins University Press, Baltimore. ISBN-13: 978-0801857898, ISBN-10: 9780801857898

Obergfell FA (1957) Vergleichende Untersuchungen an den Dentitionen und Dentale altburdigaler Cerviden von Wintershof-West in Bayern und rezenter Cerviden (eine phylogenetische Studie). Palaeontographica A 109(3/6):71–166

Paral V, Witter K, Tonar Z (2007) Microscopic examination of ground sections - a simple method for distinguishing between bone and antler? Intern J Osteoarch 17:627–634. https://doi.org/10.1002/oa.912

Pawlowska K, Stefaniak K, Nowakowski D (2004) Healed antler fracture from a giant deer (*Megalocerus giganteus*) from the Pleistocene in Poland. Palaeont Electr 17(1):23A.

Picavet PP, Balligand M (2016) Organic and mechanical properties of Cervidae antlers: a review. Veterinary Res Communic 40(3-4):141–147. DOI 10.1007/s11259-016-9663-8

Pilgrim GEB (1941) The dispersal of Artiodactyla. Biol Rev 16:134–163

Pocock RI (1923) On the External Characters of *Elaphurus, Hydropotes, Pudu*, and other Cervidæ. Proceedings of the Zoological Society of London 93(2):181–207. https://doi.org/10.1111/j.1096-3642.1923.tb02183.x

Pohle C (1989) Geburt eines Schopfhirsches im Tierpark Berlin sowie Angaben zu Gewicht und Geweihwechsel von *Elaphodus cephalophus*. Zool Garten 59:188–194

Pound MJ, Haywood AM, Salzmann U, Riding JB (2012) Global vegetation dynamics and latitudinal temperature gradients during the Mid to Late Miocene (15.97-5.33 Ma). Earth-Sci Rev 112(1-2):1–22. https://doi.org/10.1016/j.earscirev.2012.02.005

Price JS, Allen S, Faucheux C, Althnaian T, Mount JG (2005) Deer antlers: a zoological curiosity or the key to understanding organ regeneration in mammals? J Anat 207(5):603–618. https://doi.org/10.1111/j.1469-7580.2005.00478.x

Price JS, Oyajobi BO, Nalin AM, Frazer A, Graham R, Russell G, Sandell LJ (1996) Chondrogenesis in the regenerating antler tip in red deer: expression of collagen types I, IIA, IIB, and X demonstrated by in situ nucleic acid hybridization and immunocytochemistry. Develop Dyn 205(3):332–347. https://doi.org/10.1002/(SICI)1097-0177(199603)205:3%3C332::AID-AJA12%3E3.0.CO;2-6

Rörig A (1900) Über Geweihentwicklung und Geweihbildung. Archiv für Entwicklungsmechanik der Organismen 10:525–617. https://doi.org/10.1007/BF02318640

Rössner GE (1995 Odontologische und schädelanatomische Untersuchungen an *Procervulus* (Cervidae, Mammalia). Münchner geowiss. Abh. A 29:1–127

Rössner GE (2010) Systematics and palaeoecology of Ruminantia (Artiodactyla, Mammalia) from the Miocene of Sandelzhausen (southern Germany, Northern Alpine Foreland Basin). Paläontol Z 84(1), 123–162. https://doi.org/10.1007/s12542-010-0052-2

Rössner GE, Heissig K 1999 The Miocene Land Mammals of Europe. Verlag Dr. Friedrich Pfeil, München.

Rolf HJ, Enderele A (1999) Hard Fallow Deer Antler: A Living Bone Till Antler Casting? Anat Rec 255:69–77.

Rolf HJ, Fischer K, Duwel FW, Kauer F, Enderle A (2001) Histomorphology and physiology of “living” hardantlers: Evidence for substance transport into polished antlers via the vascular system. In: Sim JS, Sunwo HH, Hudson RJ, Jeon BT (eds) Antler Science and Product Technolog, ASPTRC, Edmonton, pp 97–108

Roger O (1898) Wirbelthierreste aus dem Dinotheriensande der bayerisch-schwäbischen Hochebene. Ber Naturwissenschaftl Ver Schwaben Neuburg 33:1–46

Roger O (1904) Wirbeltierreste aus dem Obermiocän der bayerisch-schwäbischen Hochebene. V. Teil. Ber Naturwiss Ver Schwaben Neuburg 36:1–19.

Rütimeyer L (1881) Beiträge zu einer natürlichen Geschichte der Hirsche. Abh schweiz paläont Ges 8:1–120.

Samejima Y, Matsuoka H (2020) A new viewpoint on antlers reveals the evolutionary history of deer (Cervidae, Mammalia). Scientific Reports 2020(10):8910. https://doi.org/10.1038/s41598-020-64555-7

Scherz M D, Daza J D, Köhler J, Vences M, Glaw F (2017) Off the scale: a new species of fish-scale gecko (Squamata: Gekkonidae: *Geckolepis*) with exceptionally large scales. PeerJ 5:e2955. https://doi.org/10.7717/peerj.2955

Schilling A-M, Rössner GE (2017) The (sleeping) Beauty in the Beast - a review on the water deer, *Hydropotes inermis.* Hystrix - It J Mam 28(2):121–133. https://doi.org/10.4404/hystrix-28.2-12362

Schlosser M (1924) Über die systematische Stellung jungtertiärer Cerviden. Centralbl Min Geol Paläont 20: 634–640.

Simpson GG (1945) Principles of classification and classification of mammals. Bull Am Mus Nat Hist 85:1–350

Sridevu PC, Prasad RV, Narayana Bhat M, Jayashankar MR, Ramjrishna V (2014) Histological studies on antler of spotted deer (*Axis axis*), horns of black buck (*Antilope cervicapra*) with that of horns of domestic cattle. J Cell Tiss Res 14(1):4131–4136

Stéger V, Molnár A, Borsy A, Gyurján I, Szabolcsi Z, Dancs G, Molnár J, Papp P, Nagy J, Puskás L, Barta E, Zomborszky Z, Horn P, Podani J, Semsey S, Lakatos P, Orosz L (2010) Antler development and coupled osteoporosis in the skeleton of red deer *Cervus elaphus*: expression dynamics for regulatory and effector genes. Molecular Genetics and Genomics 284: 273–287. https://doi.org/10.1007/s00438-010-0565-0

Stehlin HG (1928) Bemerkungen über die Hirsche von Steinheim am Aalbuch. Ecl Geol Helvetiae 21(1):245–256

Stehlin HG (1937) Bemerkungen über die miocaenen Hirschgenera *Stephanocemas* und *Lagomeryx*. Verh. Naturforsch. Ges. Basel 48:193–214

Stehlin HG (1939) *Dicroceros elegans* Lartet und sein Geweihwechsel. Ecl. Geol. Helvetiae 32(2):163–179

Stockwell R (1979) Biology of Cartilage Cells. Cambridge University Press, London. ISBN 10: 0521224101, ISBN 13: 9780521224109

Suraprasit K, Chaimanee Y, Bocherens H, Chavasseau O,Jaeger J-J. 2014. Systematics and phylogeny of middle Miocene Cervidae (Mammalia) from Mae Moh Basin (Thailand) and a paleoenvironmental estimate using enamel isotopy ofsympatric herbivore species. J Vert Paleontol 34:179–194. https://doi.org/10.1080/02724634.2013.789038

Szuwart T, Kierdorf H, Kierdorf U, Clemen G (1998) Ultrastructural aspects of cartilage formation, mineralization, and degeneration during primary antler growth in fallow deer (*Dama dama*). Ann Anat 180(6):501–510. https://doi.org/10.1016/S0940-9602(98)80055-1

Teilhard de Chardin P (1939) The Miocene cervids from Shantung. Bull Geol Soc China 19(3):269–278

Teilhard de Chardin P, Trassaert M (1937) The Pliocene Camelidae, Giraffidae and Cervidae of S. E. Shansi. Paläontolog. Sinica, NS C 102(1):1–56

Thenius E (1948a) Zur Kenntnis der fossilen Hirsche des Wiener Beckens, unter besonderer Berücksichtigung ihrer Stratigraphischen Bedeutung. Ann Naturhist Mus Wien 56:262–308

Thenius E (1948b) Über ein stammesgeschichtlich interessantes Stadium aus der Geschichte der Hirsche. Anz Österr Akad Wissensch Mathem-naturw Kl 14:219–254

Thenius E (1959) Wirbeltierfaunen. In: Papp A, Thenius E (eds) Tertiär, 2^nd^ sec. In: Lotze F (ed) Handbuch der stratigraphischen Geologie, Ferdinand Enke Verlag, Stuttgart

van Mourik S, Stelmasiak T (1990) Endocrine mechanisms and antler cycle in rusa deer, *Cervus rusa timorensis*. In: Bubenik GA, Bubenik AB (eds), Horns, Pronghorns and Antlers, Springer, New York, pp 413–421. ISBN-10: 1461389682, ISBN-13: 978-1461389682

Van Bemmel ACV (1952) Contribution to the knowledge of the genera *Muntiacus* and *Arctogalidia* in the Indoaustralian archipelago. Beaufortia 16:1–50

Vislobokova IA (1983) Fossil deer of Mongolia. Transact Paleont Inst 23:1–75 [in Russian]

Vislobokova IA (1990) Fossil deer of Eurasia. Transact Paleont Inst 240:1–204 [in Russian]

Vislobokova IA (2013) Morphology, taxonomy, and phylogeny of megacerines (Megacerini, Cervidae, Artiodactyla). Paleont J 47:833–950. https://doi.org/10.1134/S0031030113080017

Vislobokova IA, Godina AY (1993a) The use of the structural analysis of cranial appendages for the definition of the systematic position of fossil ruminants. Paleont Zh 3:86–98 [In Russian]

Vislobokova I A, Godina A Y (1993b) Application of the structural analysis of frontal appendages for determining the systematic position of fossil higher ruminants. Palaeont J 27(3):108–123

Vislobokova I., Hu Changkang, Sun Bo (1989) On the systematic position of the Lagomerycinae. Vertebrata PalAsiatica 28(2):128–132

Waldo CM, Wislocki GB (1951) Observations on the shedding of the antlers of Virginia deer (*Odocoileus virginianus borealis*). Developm Dynam 88:351–395. https://doi.org/10.1002/aja.1000880303

Wang D, Berg D, Ba H, Sun H, Wang Z Li C (2019) Deer antler stem cells are a novel type of cells that sustain full regeneration of a mammalian organ — deer antler. Cell Death Desease 10:443. https://doi.org/10.1038/s41419-019-1686-y

Wang X, Xie G, Dong WA (2009) A new species of crown-antlered deer *Stephanocemas* (Artiodactyla, Cervidae) from the middle Miocene of Qaidam Basin, northern Tibetan Plateau, China, and a preliminary evaluation of its phylogeny. Zool J Linnean Soc 156(3):680–695. https://doi.org/10.1002/aja.1000880303

Wang Y, Zhang C, Wang N, Li Z, Heller R, Liu R, Zhao Y, Han J, Pan X, Zheng Z, Dai X, Chen C, Dou M, Peng S, Chen X, Liu J, Li M, Wang K, Liu C, Lin Z, Chen L, Hao F, Zhu W, Song C, Zhao C, Zheng C, Wang J, Hu S, Li C, Yang H, Jiang L, Li G, Liu M, Sonstegard TS, Zhang G, Jiang Y, Wang W, Qiu Q (2019) Genetic basis of ruminant headgear and rapid antler regeneration. Science 364:eaav6335. https://doi.org/10.1126/science.aav6335

Whitehead GK (1972) Deer of the world. Constance, London. ISBN 10: 0094560307, ISBN 13: 9780094560307

Wislocki GB (1942) Studies on the growth of deer antlers. I. On the structure and histogenesis of antlers of the Virginian deer (*Odocoileus virginianus borealis*). Am J Anat 71:371–415. https://doi.org/10.1002/aja.1000710304

Wislocki GB, Weatherford HL, Singer M (1947) Osteogenesis of antlers investigated by histological and histochemical methods. Anat Rec 99(3):263–295. https://doi.org/10.1002/ar.1090990305

Wislocki GB (1956) Further Notes on Antlers in Female Deer of the Genus *Odocoileus*. J Mamm 37(2):231–235. https://doi.org/10.2307/1376682

Young CC 1937 On a Miocene mammalian fauna from Shantung. Bull Geol Soc China 17(2):210–238.

Young CC (1964) On a new *Lagomeryx* from Lantian, Shansi. Vertebr PalAsiatica 11:329–340

Zachos J, Pagani M, Sloan L, Thomas E, Billups K (2001) Trends, rhythms, and aberrations in global climate 65 Ma to present. Science 292:686–693. https://doi.org/10.1126/science.1059412

Zdansky O (1925) Fossile Hirsche Chinas. Pal Sin C2(3): 1–94

Zittel KA (1893) Vertebrata (Mammalia) Handbuch der Palaeontologie. In: Zittel KA (ed) Handbuch der Palaeontologie, 1st sec, Palaeozoologie, 4th vol, R. Oldenbourg, München, Leipzig

