## supplemental Table 1 and Figures 1-8 for "Antiquity and fundamental processes of the antler cycle in Cervidae (Mammalia)"

Online Resource 1. Specimens investigated. NMA = Naturmuseum Augsburg, NMB = Naturhistorisches Museum Basel, SMNS = Staatliches Museum für Naturkunde Stuttgart, SNSB – BSPG = Staatliche Naturwissenschaftliche Sammlungen Bayerns – Bayerische Staatssammlung für Paläontologie und Geologie, SNSB – ZSM = Staatliche Naturwissenschaftliche Sammlungen Bayerns – Zoologische Staatssammlung München. For biochronological assessment of sites of provenance see de Bruijn et al. (1992) and for geographical location Thenius (1959).

| **Catalogue and Figure Number** | **Species Identification** | **Site of Provenance** | **Geological Age** | **Specimen** | **Method of Examination** | **Literature Presence** |
|---|---|---|---|---|---|---|
| SNSB-ZSM 1966 237b (Online Resource 36) | *Muntiacus muntjak* (Zimmerman, 1780) | zoo specimen, Tierpark Hellabrunn München | Recent | Shed antler | µCT (110 kV, 0.05 mA, 0.1 mm Cu filter, 1400 images, voxel size 27.5 µm) | Heckeberg (2017b: Figs 1G-I) |
| SNSB-BSPG 1950 I 30 (Online Resource 35) | *Euprox furcatus* (Hensel, 1859) | Massenhausen, Germany | Middle Miocene, MN8 | Attached antler | µCT (120 kV, 0.06 mA, 0.2 mm Cu filter, 1440 images, voxel size 68.1 µm) | Gentry et al. (1999: Fig. 23.13) |
| SNSB-BSPG 1966 XIV 34 (Online Resource 34) | *Euprox furcatus* (Hensel, 1859) | Breitenbrunn, Germany | Middle Miocene, MN8 | Shed antler | µCT (120 kV, 0.06 mA, 0.2 mm Cu filter, 1440 images, voxel size 51.4 µm) | Gentry et al. (1999: Fig. 23.13); Heckeberg (2017b: Figs 6E-F) |
| NMB Sth. 12 (Online Resources 1, 2, 8) | *Euprox furcatus* (Hensel, 1859) | Steinheim, Germany | Middle Miocene, MN8 | pedicle without antlers | thin-sections | |
| SMNS (Online Resource 33) | *Heteroprox larteti* (Filhol, 1890) | Steinheim, Germany | Middle Miocene, MN7 | skull with attached antlers | µCT (160 kV, 0.07 mA, 1440 images, voxel size 66.7 µm) | |
| NMA 79-5004/761 **holotype** (Online Resource 32) | *Paradicrocerus elegantulus* (Roger, 1898) | Stätzling, Germany | Middle Miocene, MN6 | Attached antler | µCT (130 kV, 0.037 mA, 0.2 mm Cu filter, 1400 images, voxel size 66.7 µm) | Roger (1904: pl. III, fig. 1); Stehlin (1937: Fig. 1); Azanza & Menéndez (1990: Fig. 3.3) |
| SNSB-BSPG 1976 VI 24 (Online Resources 1, 2, 12) | *Paradicrocerus elegantulus* (Roger, 1898) | Thannhausen, Germany | Middle Miocene, MN6 | Shed antler | thin-sections | |
| SNSB-BSPG 1993 I 21 (Online Resource 31) | *Paradicrocerus elegantulus* (Roger, 1898) | Wollersdorf, Germany | Middle Miocene, MN5 | Shed antler | µCT (120 kV, 0.06 mA, 0.2 mm Cu filter, 1440 images, voxel size 51.6 µm) | Heckeberg (2017b: Figs 5E-F) |
| SNSB-BSPG 1993 I 35 (Online Resource 30) | *Dicrocerus elegans* Lartet, 1837 | Sansan, France | Middle Miocene, MN6 | Attached antler | µ-CT (130 kV, 0.060 mA, 0.2 mm Cu filter, 1400 images, voxel size 54.6 µm) | Gentry et al. (1999: Fig 23.12) |
| NMB San. 15061 (Online Resources 1, 2, 11) | *Dicrocerus elegans* Lartet, 1837 | Sansan, France | Middle Miocene, MN6 | Tine, proximal portion | thin-sections | |
| NMB San. 15062 (Online Resource 1, 2, 11) | *Dicrocerus elegans* Lartet, 1837 | Sansan, France | Middle Miocene, MN6 | Tine, distal portion | thin sections | |
| SNSB-BSPG 1959 II 678 (Online Resource 29) | *Lagomeryx parvulus* Roger, 1898 | Sandelzhausen, Germany | Middle Miocene, MN5 | attached antler | µCT (120 kV, 0.06 mA, 0.1 mm Cu filter, 1438 images, voxel size 11.4 µm) | Fahlbusch (1977: Pl. 16 Figs 1-7); Rössner (2010: Fig. 8Y) |

Created in Master PDF Editor

| | | | | | | |
|---|---|---|---|---|---|---|
| SNSB-BSPG 1959 II 4594 (Online Resources 1, 2, 9, 10) | *Lagomeryx parvulus* Roger, 1898 | Sandelzhausen, Germany | Middle Miocene, MN5 | Attached antler | thin-sections | Rössner (2010: Figs 8A, G) |
| SNSB-BSPG 1959 II 12314 (Online Resource 1, 2, 3, 7) | *Heteroprox eggeri* Rössner, 2010 | Sandelzhausen, Germany | Middle Miocene, MN5 | Attached antler | thin-sections | |
| SNSB-BSPG 1959 II 5270 (Online Resource 1, 2, 3, 6) | *Heteroprox eggeri* Rössner, 2010 | Sandelzhausen, Germany | Middle Miocene, MN5 | Shed antler | thin-sections | |
| SNSB-BSPG 1959 II 5249 **holotype** (Online Resource 28) | *Heteroprox eggeri* Rössner, 2010 | SandelzhausenGermany | Middle Miocene, MN5 | Attached antler, fully grown, tips missing | µCT (120 kV, 0.06 mA, 0.1 mm Cu filter, 1650 images, voxel size 52.3 µm) | Rössner (2010: Fig. 6A) |
| SNSB-BSPG 1959 II 2502 (Online Resource 27) | *Heteroprox eggeri* Rössner, 2010 | Sandelzhausen, Germany | Middle Miocene, MN5 | Attached antler; unbranched, ?first generation | µCT (120 kV, 60 µA, 0.1 mm Cu filter, 1650 images, voxel size 45.4 µm) | Rössner (2010: Fig. 6B) |
| SNSB-BSPG 1959 II 5258 (Online Resource 26) | *Heteroprox eggeri* Rössner, 2010 | Sandelzhausen, Germany | Middle Miocene, MN5 | Shed antler, base preserved only | µCT (120 kV, 60 µA, 0.1 mm Cu filter, 1400 images, voxel size 27.1 µm) | Rössner (2010: Fig. 6E); Heckeberg (2017b: Figs 2E-F) |
| SNSB-BSPG 1959 II 5268 (Online Resource 25) | *Heteroprox eggeri* Rössner, 2010 | Sandelzhausen, Germany | Middle Miocene, MN5 | Shed antler, base preserved only | µCT (130 kV, 30 µA, 0.1 mm Cu filter, 1400 images, voxel size 25.8 µm) | Rössner (2010: Fig. 6C) |
| SNSB-BSPG 1976 XXI 64 (Online Resource 24) | *Procervulus dichotomus* (Gervais, 1849) | Langenau 2, Germany | Early Miocene, MN4 | Shed antler | µCT (120 kV, 50 µA, 0.1 mm Cu filter, 1700 images, voxel size 33.3 µm) | Heckeberg (2017b: Figs 3K-L) |
| SMNS 45140 (Online Resource 37) | *Procervulus dichotomus* (Gervais, 1849) | Langenau, Germany | Early Miocene, MN4 | Skull with left and right pedicle bud and dentition with M3 erupted but unworn | µCT (130 kV, 30 µA, 0.2 mm Cu filter, 1400 images, voxel size 66.7 µm) | |
| SNSB-BSPG 1979 XV 555 (Online Resource 23) | *Procervulus dichotomus* (Gervais, 1849) | Rauscheröd, Germany | Early Miocene, MN4b | Skull with both cranial appendages attached and heavily worn dentition | µCT (120 kV, 100 µA, 0.1 mm Cu filter, 1500 images, voxel size 66.7 µm) | Rössner (1995: Pl. 7, Fig. 1), Gentry et al. (1999: Fig. 23.10); |
| SNSB-BSPG 1881 IX 55m **holotype** (Online Resource 22) | *Lagomeryx ruetimeyeri* Thenius, 1948 | Reisensburg, Germany | ?Early Miocene, ?MN4 | Attached antler | µCT (120 kV, 60 µA, 0.2 mm Cu filter, 1400 images voxel size 17.8 µm) | Rütimeyer (1881: Pl. 1, Figs 2-5); Stehlin (1937: Fig. 9); Gentry & Heizmann (1993); Bubenik (1990: Fig. 8.1); Gentry et al. (1999: Fig. 23.11) |
| SNSB-BSPG 1937 II 16845 (Online Resource 21) | *Procervulus praelucidus* (Obergfell, 1957) | Wintershof-West, Germany | EarLower Miocene, MN3b | Shed antler, incomplete | µCT (120 kV, 60 µA, 0.2 mm Cu filter, 1400 images, voxel size 22.2 µm) | Rössner (1995: Pl. 6, Fig. 8) |
| SNSB-BSPG 1937 II 16842 (Online Resource 20) | *Procervulus praelucidus* (Obergfell, 1957) | Wintershof-West, Germany | Lower Miocene, MN3b | Shed antler | µCT (120 kV, 60 µA, 0.1 mm Cu filter, 1650 images, voxel size 21.8 µm) | Rössner (1995: Taf. 6, Figs 4, 9) |
| SNSB-BSPG 1937 II 16841 (Online Resource 19) | *Procervulus praelucidus* (Obergfell, 1957) | Wintershof-West, Germany | Lower Miocene, MN3b | Attached antler | µCT (130 kV, 40 µA, 0.2 mm Cu filter, 1400 images, voxel size 54.6 µm) | Rössner (1995: Taf. 5, Fig. 3); Gentry et al. (1999: Fig. 23.8) |

Created in Master PDF Editor

| | | | | | | |
|---|---|---|---|---|---|---|
| SNSB-BSPG 1937 II 16810 (Online Resource 18) | *Procervulus praelucidus* (Obergfell, 1957) | Wintershof-West, Germany | Lower Miocene, MN3b | Antler broken from pedicle | µCT (122 kV, 40 µA, 0.1 mm Cu filter, 1400 images, voxel size 24.4 µm) | Heckeberg (2017b: Figs 3G-H) |
| SNSB-BSPG 1937 II 16787 (Online Resources 1, 2, 3, 4) | *Procervulus praelucidus* (Obergfell, 1957) | Wintershof-West, Germany | Lower Miocene, MN3b | Attached antler | thin-sections | |
| NMB S.O. 3020 **lectotype** (Online Resource 17) | *Ligeromeryx praestans* (Stehlin, 1937) | Chitenay, France | Lower Miocene, MN3b | Attached antler | µCT (130 kV, 30 µA, 0.2 mm Cu filter, 1400 images, voxel size 52,6 µm) | Stehlin (1937: Fig. 10); Azanza & Ginsburg (1997: Text-Fig. 2D-E); Heckeberg (2017b: fig 8E (cast of original)) |
| NMB S.O. 5720 **paralectotype** (Online Resource 16) | *Ligeromeryx praestans* (Stehlin, 1937) | Chitenay, France | Lower Miocene, MN3b | Shed antler | µCT (130 kV, 30 µA, 0.2 mm Cu filter, 1400 images, voxel size 42 µm ) | Stehlin (1937: Fig. 11); Bubenik (1990: 18B); Azanza & Ginsburg (1997: Text-Fig. 2A, 3H); Stehlin (1937: fig. 11) |
| NMB S.O. 2078 **paralectotype** (Online Resource 15) | *Ligeromeryx praestans* (Stehlin, 1937) | Chitenay, France | Lower Miocene, MN3b | Shed antler | µCT (130 kV, 30 µA, 0.2 mm Cu filter, 1400 images, voxel size 38,3 µm) | Stehlin (1937: Fig. 12); Azanza & Ginsburg (1997: Text-Fig. 3C); Heckeberg (2017b: Figs 8C-D (cast of original)) |
| NMB S.O. 2077 (Online Resources 1, 2, 3, 5) | ?*Ligeromeryx praestans* (Stehlin, 1937) | Chitenay, France | Lower Miocene, MN3b | piece of frontal bone with pedicle | thin-sections | |
| NMB S.O. 3126 **holotype** (Online Resource 14) | *Acteocemas infans* (Stehlin, 1939) | Chilleur, France | Lower Miocene, MN3b | Attached antler | µCT (130 kV, 30 µA, 0.2 mm Cu filter, 1400 images, voxel size 47,2 µm) | Stehlin (1939: Fig. 11); Azanza Asensio (2000: Fig. 18) |
| NMB S.O. 3024 (Online Resource 13) | ?*Ligeromeryx praestans* (Stehlin, 1937) | Chitenay, France | Lower Miocene, MN3b | Attached antler, questionable if juvenile or senile | µCT (130 kV, 30 µA, 0.2 mm Cu filter, 1350 images, voxel size 31 µm) | |

Created in Master PDF Editor

**Online Resource 2:** Specimens (antler fragments and pedicles) before histological sectioning. A, *Procervulus praelucidus* (SNSB - BSPG 1937 II 16787), bifurcate antler with pedicle from Wintershof-West near Eichstätt, southern Germany; lateral view. B, *Ligeromeryx praestans* (NMB S.O. 2077), base of the pedicle from Chitenay, central France; lateral view. C, *Heteroprox eggeri* (SNSB - BSPG 1959 II 5270), bifurcate antler from Sandelzhausen, southern Germany; lateral view. D, *Lagomeryx parvulus* (SNSB - BSPG 1959 II 4594), multipointed antler with incomplete pedicle from Sandelzhausen, southern Germany. E, F, *Paradicrocerus elegantulus* (SNSB - BSPG 1976 VI 24), base of shed dichotomous antler with abscission scar from Thannhausen, southern Germany; distal view and proximal view. G, *Euprox* furcatus (NMB Sth.12), full pedicle with distal abscission scar from Steinheim, Germany; laterofrontal view. H-I, *Dicrocerus* elegans (NMB San. 15062), incomplete tine (distal portion) of dichotomous antler from Sansan, southern France; frontal or posterior view and lateral view. J, *Dicrocerus elegans* (NMB San. 15061), incomplete tine (proximal portion) of dichotomous antler from Sansan, southern France; frontal view. K, *Heteroprox eggeri* (SNSB - BSPG 1959 II 12314), one-tipped antler with missing tip and with full pedicle from Sandelzhausen, southern Germany, laterofrontal view. Black arrowheads indicate position of sections taken in addition to longitudinal sections (exception: the *Paradicrocerus* specimen was only sectioned as indicated by black arrow heads). Labeling of indicated close-ups refer to further online resources.

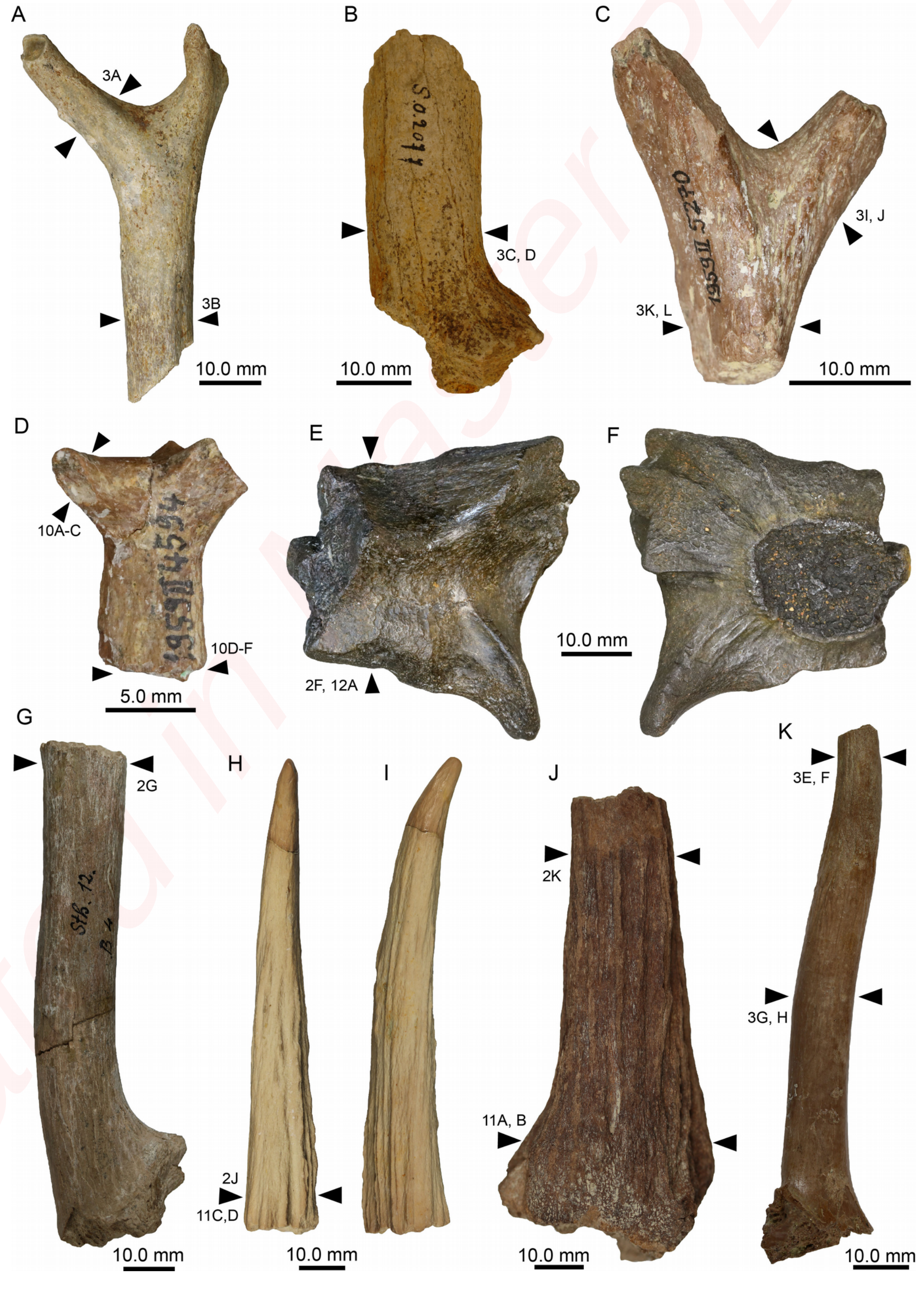


Created in Master PDF Editor

**Online Resource 3**: Sectioned specimens in normal transmitted light (all longitudinal sections except image B, a half cross-section, and images G, J, and K, which represent complete cross sections). A, *Procervulus praelucidus* (SNSB - BSPG 1937 II 16787). B, C, *Ligeromeryx praestans* (NMB S.O. 2077). D, *Heteroprox eggeri* (SNSB - BSPG 1959 II 5270). E, *Lagomeryx parvulus* (SNSB - BSPG 1959 II 4594). F, *Paradicrocerus elegantulus* (SNSB - BSPG 1976 VI 24). G, H, *Euprox* furcatus (NMB Sth. 12). I-J *Dicrocerus* elegans (NMB San.15062). K-L, *Dicrocerus* elegans (NMB San.15061). M, *Heteroprox eggeri* (SNSB - BSPG 1959 II 12314). Labeling of indicated close-ups refer to further online resources.

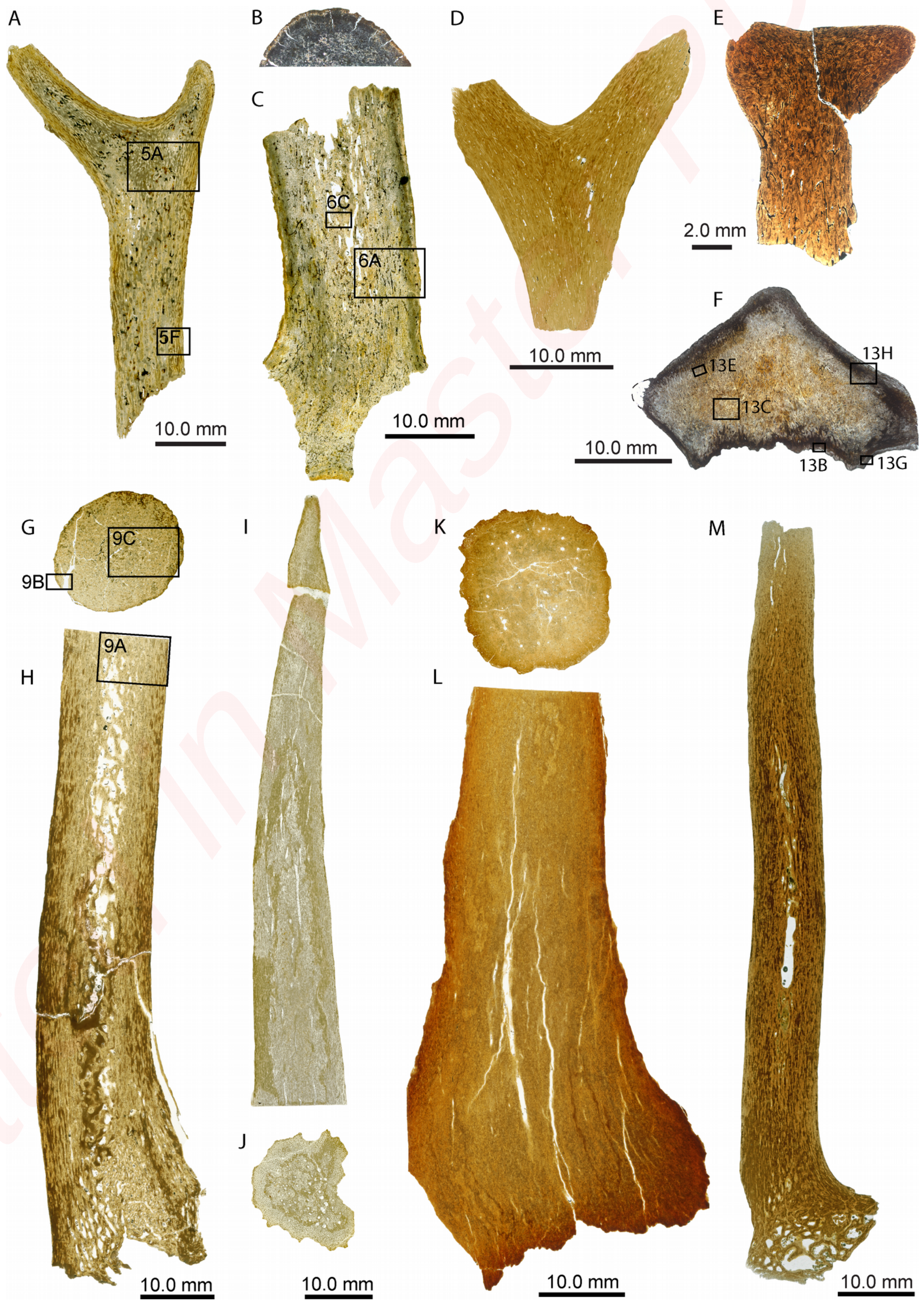


Created in Master PDF Editor

**Online Resource 4**: Comparison of cross sections through antlers' tines (A, E, F, I-L) with those through the pedicles (B-D, G and H) of *Procervulus praelucidus, Ligeromeryx praestans* and *Heteroprox eggeri*. Images in A-C, E, G, I, and K in normal transmitted light; images in D, F, H, J, and L in cross-polarised light using lambda compensator. A, B, *Procervulus praelucidus* (SNSB - BSPG 1937 II 16787). C, D, *Ligeromeryx praestans* (NMB S.O. 2077). E-H, *Heteroprox eggeri* (SNSB - BSPG 1959 II 12314). I-L, *Heteroprox eggeri* (SNSB - BSPG 1959 II 5270). Note that the histology of the antler (A) and pedicle (B) in *Procervulus praelucidus* is very similar, consisting mostly of secondary remodelled bone trabeculae and somewhat uniformly sized vascular spaces, whereas larger trabeculae and spaces in the central regions of the pedicle are absent (Online Resource Figure 2A). The pedicle of the larger *Ligeromeryx praestans* (C, D) holds similar bone structures, with the exception of larger central vascular spaces (see Online Resource Figure 2C). The cross-sections through the tines and pedicle of *Heteroprox eggeri* SNSB - BSPG 1959 II 12314 (E-H) are also compact with few larger vascular spaces (larger vascular spaces were found only within the proximal and mid-regions of the pedicle (see Online Resource Figure 2M). The cross-section through the antler part in SNSB - BSPG 1959 II 12314 reveals still more primary bone (with reticular or laminar organisation of primary osteons) and less dense remodelling in form of secondary osteons, compared to the section through the pedicle. The distal (I, J) and proximal (K, L) cross-sections of the shed *Heteroprox eggeri* antler SNSB - BSPG 1959 II 5270 both show densely remodelled bone interiorly, with remnants of primary bone tissue (lamellar/parallel-fibred bone). Remodelling appears more extensive in the proximal part of the antler compared with the more distal tine.

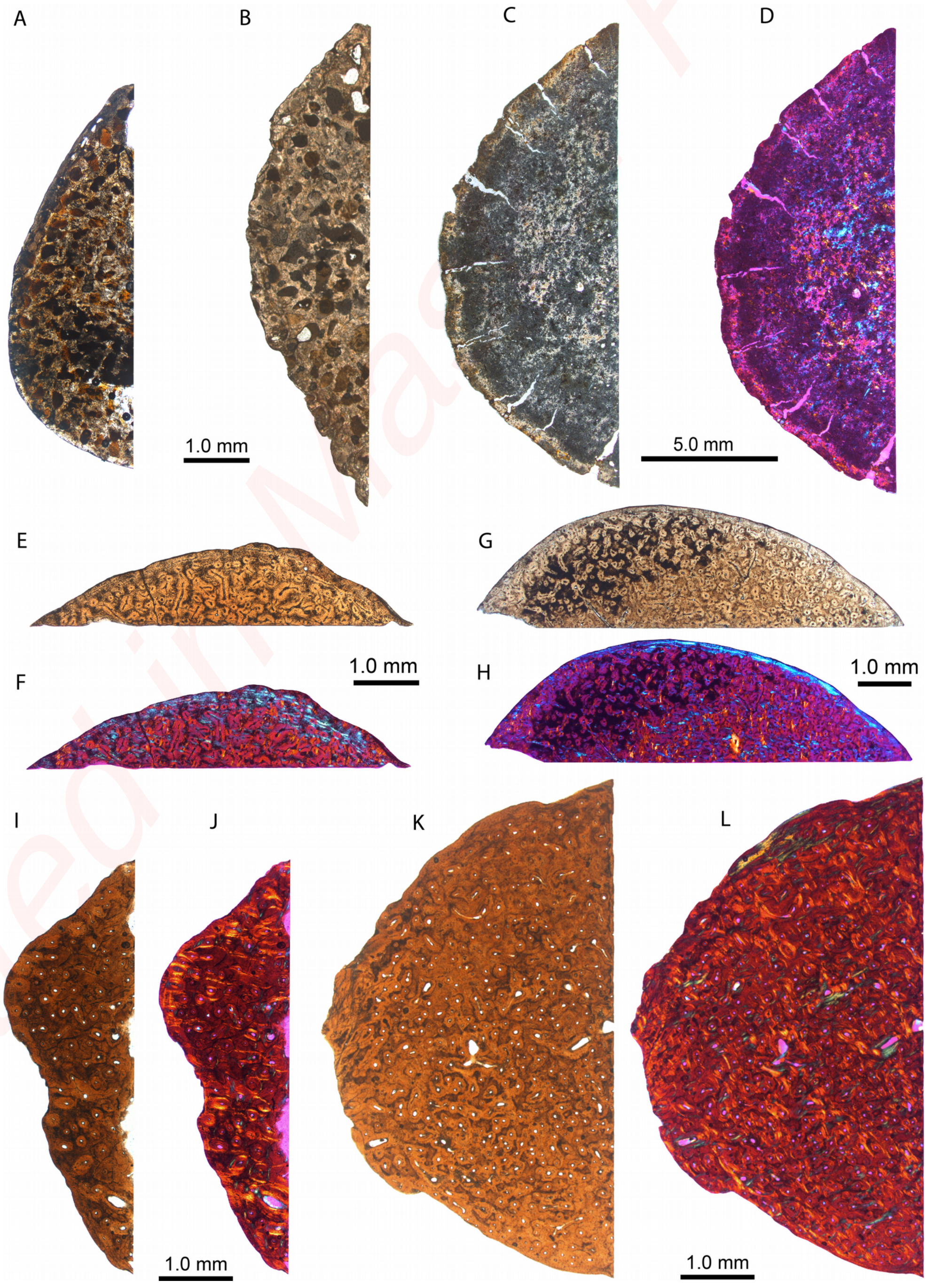


Created in Master PDF Editor

**Online Resource 5:** Detailed histology of antler attached to pedicle of *Procervulus praelucidus* (SNSB - BSPG 1937 II 16787). Images in A, C, E-G in normal transmitted light; B and D in cross-polarised light using lambda compensator. Position of longitudinal close-ups are indicated in Online Resource Figure 2A. A, B, Close-up of internal trabecular bone and peripheral primary bone of the cortex in the proximal part of the antler in longitudinal section (position of the close-up is indicated in Online Resource Figure 2A). C, D, Close-up of the smaller tine of the specimen in longitudinal section. Internal trabecular bone is framed by a laminar organisation of primary bone and vascularisation showing lamellar bone lining (i.e., primary osteons extending subparallel to the bone surface). Cell lacunae are more globular without canaliculi.  E, Close-up of the peripheral bone of the tine retaining primary osteons and larger erosion cavities in cross-section. F, Close-up of the peripheral bone of the pedicle in longitudinal section. G, Close-up of the peripheral bone of the pedicle in cross-section. Abbreviations: EC, erosion cavity; HC, Haversian canal of secondary osteon; LB, lamellar bone; PO, primary osteon; TB, trabecular bone.

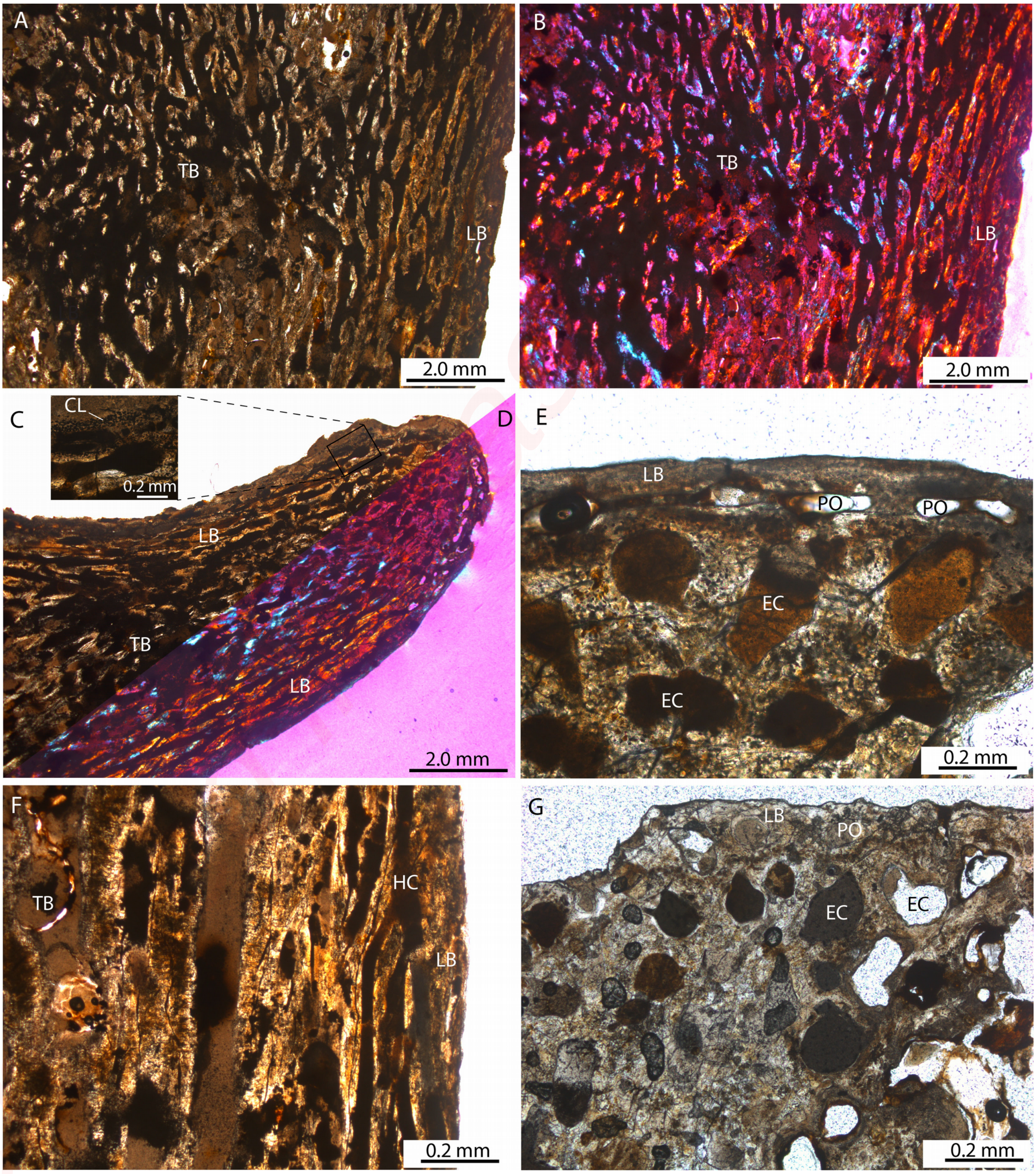

**Online Resource 6:** Detailed histology of a pedicle of *Ligeromeryx praestans* (NMB S.O. 2077) in longitudinal (A-D) and cross-section (E, F). The position of the longitudinal close-ups are indicated in Fig. 2C. Images in A, C and E in normal transmitted light; images in B, D and F in cross-polarised light. A, B, Interiorly the pedicle shows secondary trabecular bone composed of lamellar bone tissue, whereas the remainder of the tissue is dense Haversian tissue (composed of the lamellar bone of more or less longitudinally arranged secondary osteons). In the periphery of the pedicle, a thin layer of primary bone tissue is preserved (also consisting of lamellar bone). C, D, Close-up of the trabecular bone in the mid-region of the specimen. E, F, Close-up of the outer cortical bone composed mainly of secondary osteons and remnants of primary bone composed of lamellar bone tissue. Abbreviations: LB, lamellar bone; SO, secondary osteon; TR, trabecular bone.

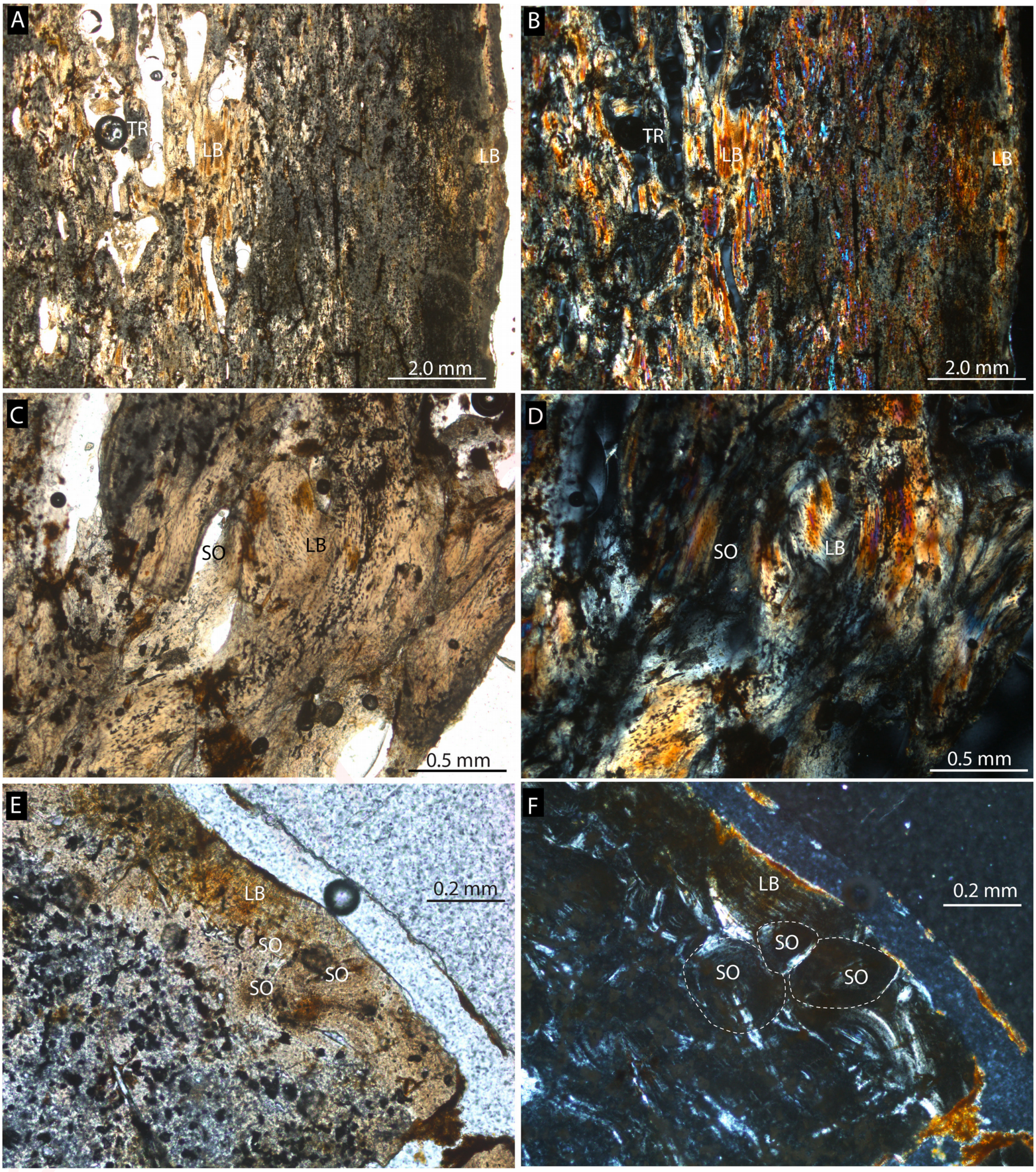

**Online Resource 7:** Detailed histology of a shed antler of *Heteroprox eggeri* (SNSB - BSPG 1959 II 5270) in longitudinal section (A-D) and cross-sections through the proximal antler (E, F) and through the distal portion of a tine (G, H). Images in C, E and G in normal transmitted light, D, F and H in cross-polarised light, and images in A and B in cross-polarised light using lambda compensator. A, B, Proximal and distal portions of the tines, being mainly composed of the lamellar bone tissue of longitudinally sectioned secondary osteons. Note divergence of bone fibres (indicated by the colour difference) where the two tines branch off. In this area, small irregular erosion cavities are found. C-F, Close-up of the compact bone of the proximal antler. Note thin primary bone in the cortical periphery consisting of lamellar/parallel-fibred bone (well visible in E; note also presence of Sharpey's fibres and the strongly remodelled interior bone largely consisting of dense Haversian bone. G, H, Focus on the bone tissue of the distal part of the tine. Here, most of the bone is also remodelled into dense Haversian tissue, and the external-most layer still consists of primary lamellar/parallel-fibred bone tissue, crossed by thin Sharpey's fibres. Abbreviations: EC, erosion cavity; LB, lamellar bone; ShF, Sharpey's fibres; SO, secondary osteon.

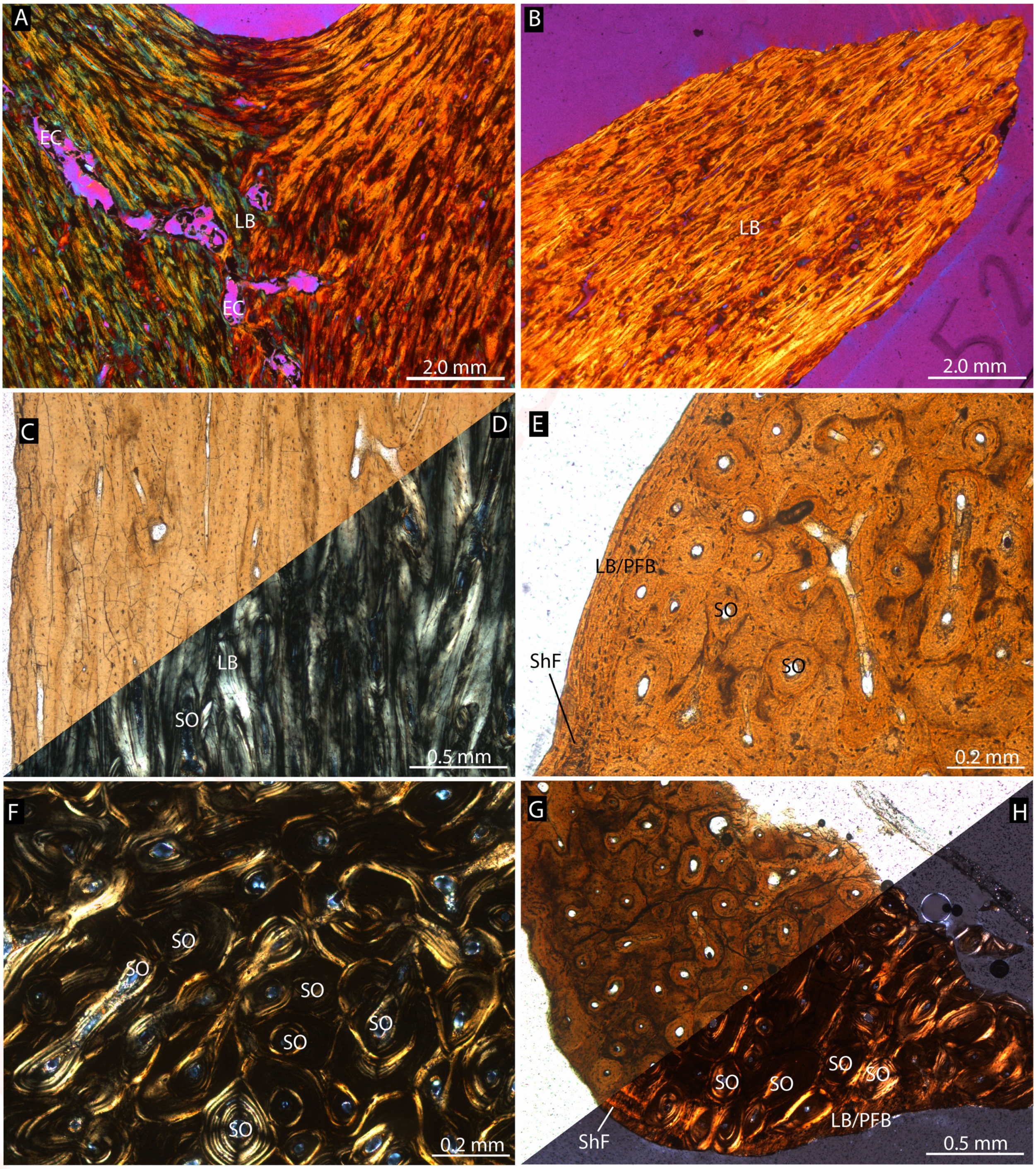

**Online Resource 8:** Detailed histology of cranial appendage with one-tipped antler of *Heteroprox eggeri* (SNSB - BSPG 1959 II 12314) in longitudinal (A, B) and cross sections (C, D). Images in A-C are in normal transmitted light, D in cross-polarised light using lambda compensator. A, Close-up of the distal portion of the antler. Note the absence of strong Sharpey's fibres here. B, Close-up of the mid-portion of the pedicle showing numerous strong (coarse) Sharpey's fibres. C, D, Cross-section of the mid-part of the pedicle showing mostly secondary remodelled bone tissue and a thin remnant of non-appositional primary bone consisting of lamellar bone, crossed by Sharpey's fibres. Abbreviations: HC, Haversian canal of secondary osteon; LB, lamellar bone; ShF, Sharpey's fibres; SO, secondary osteon.

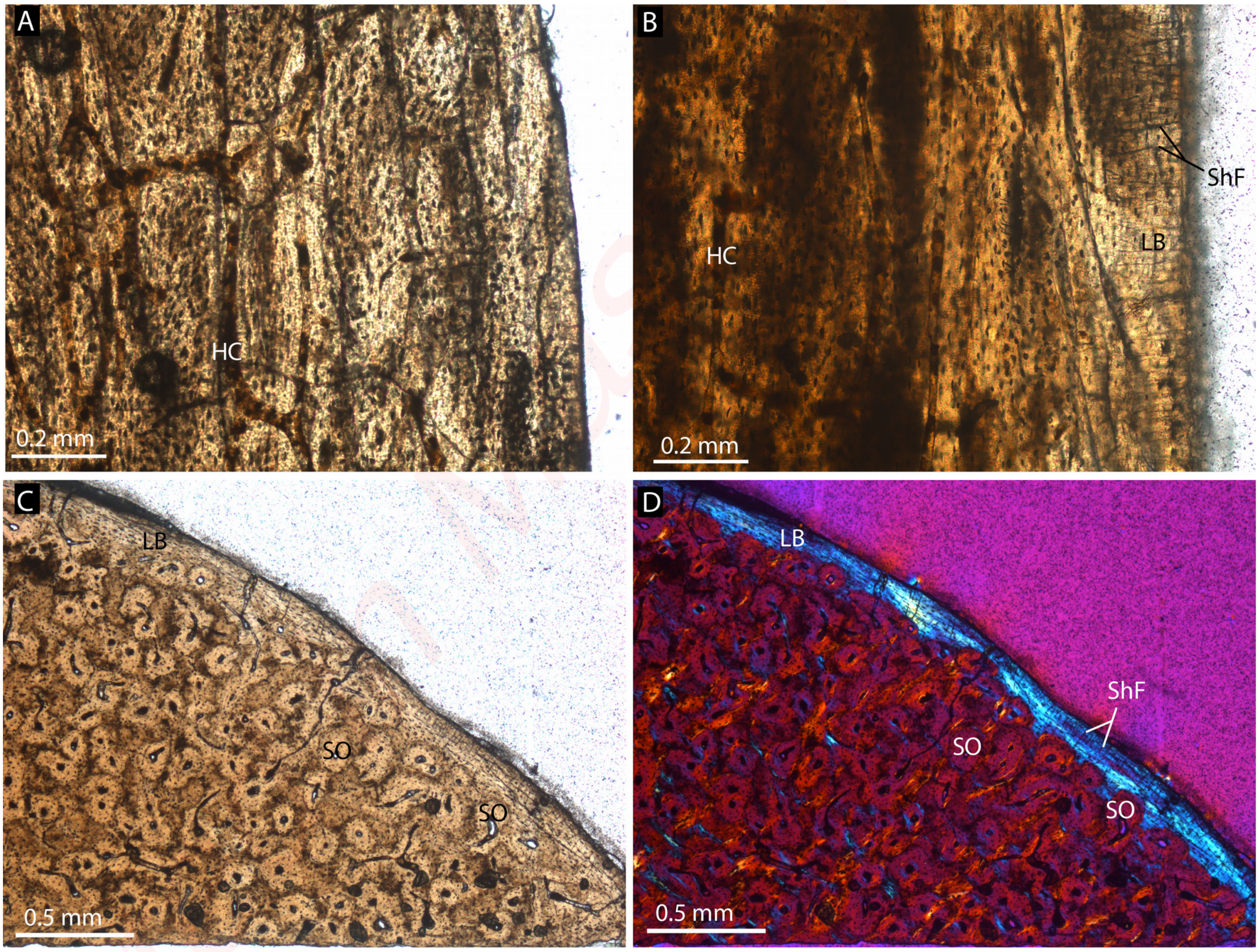
