## supplemental Figures 9-38 for "Antiquity and fundamental processes of the antler cycle in Cervidae (Mammalia)"

**Online Resource 9:** Detailed histology of the pedicle of *Euprox furcatus* (NMB Sth. 12) in longitudinal (A) and cross section (B-F). Images in A-E are in normal transmitted light, F in cross-polarised light using lambda compensator. A, Close-up of distal portion of pedicle, just below the abscission area (see Fig. 1G), showing interior trabecular bone, largely remodelled, and a compact cortex. Note the decrease in size of the vascular spaces towards the top of image, which represents the level of the cross-section depicted in B-F (position of focus areas are indicated in Fig. 2G, H). B, Peripheral lamellar bone of the compacta, vascularised by few scattered primary and secondary osteons. Note presence of Sharpey's fibres. C, Patches of primary bone tissue with reticular vascularisation, within largely remodelled Haversian bone tissue. D, Close-up of patch of primary bone. E, F, Close-up of the multiple generations of secondary osteons forming dense Haversian bone. Abbreviations: EC, erosion cavity; LB-PFB, lamellar bone to parallel-fibred bone; PB, patches of primary bone tissue; RV, reticular vascularisation pattern; ShF, Sharpey's fibres; SO, secondary osteon.

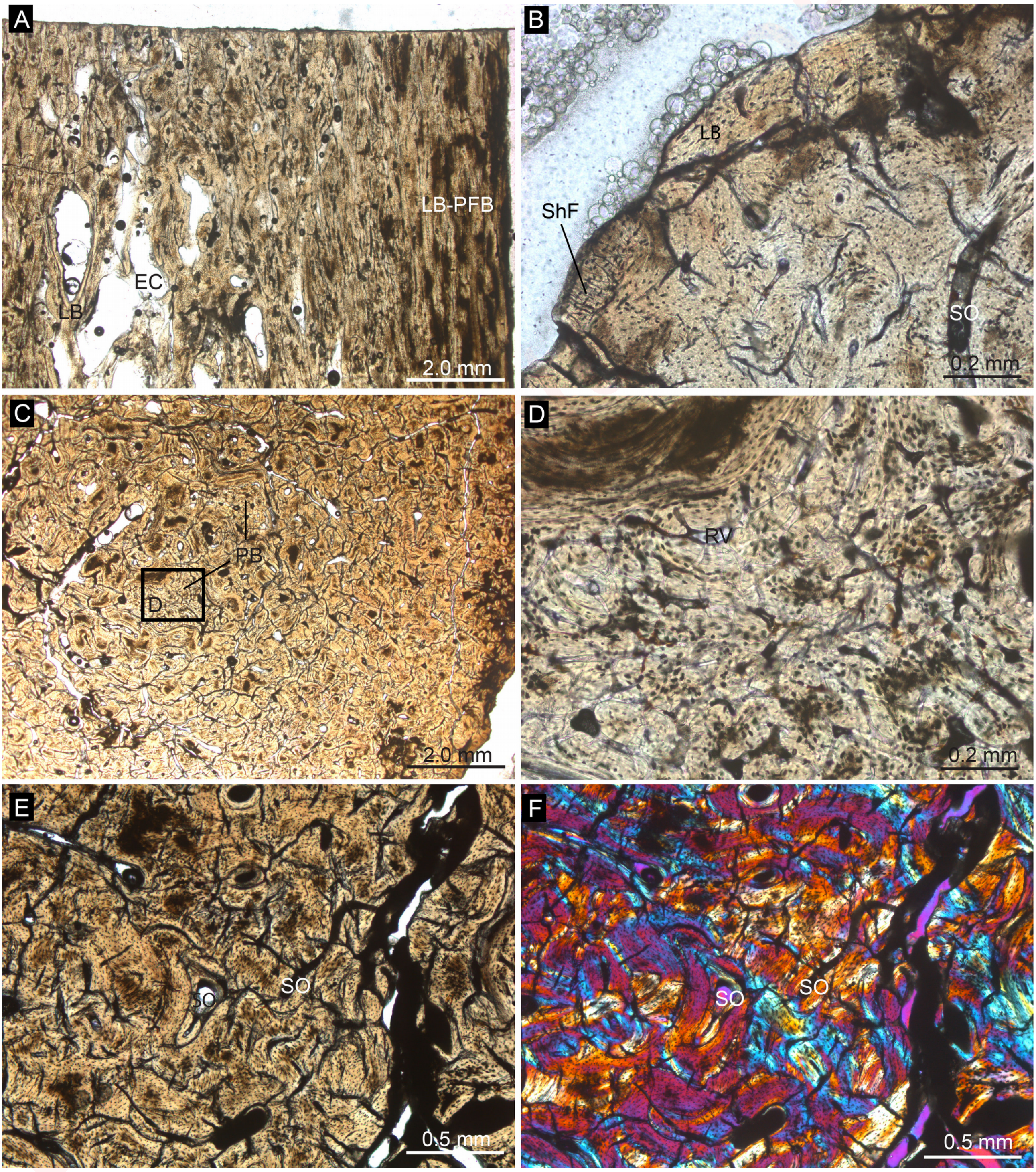

**Online Resource 10:** Detailed histology of a cranial appendage of *Lagomeryx parvulus* (SNSB - BSPG 1959 II 4594) in longitudinal section. Image C in normal transmitted light, images in A, D and F in cross-polarised light, and images B, E, and G in cross-polarised light using lambda compensator. A, B, Composite image of the complete longitudinal section of the specimen showing the deviating structures in the pedicle and the antler. C-E, Focus on the cortex and interior bone tissue just proximal of the antler's base (see inset in A). The bone is composed mainly of longitudinally extending and frequently branching secondary osteons, although a thin remnant of primary lamellar bone crossed by Sharpey's fibres is still preserved. The border between primary and secondary bone is marked by the white stippled line. F, G, Close-up of the cortical region of the antler (see inset in A for position). Note that remnants of primary bone are found in the form of primary osteons extending obliquely towards the antler bone surface. Abbreviations: HC, Haversian canal of secondary osteon; LB, lamellar bone; ShF, Sharpey's fibres; VC, Volkmann's canals.

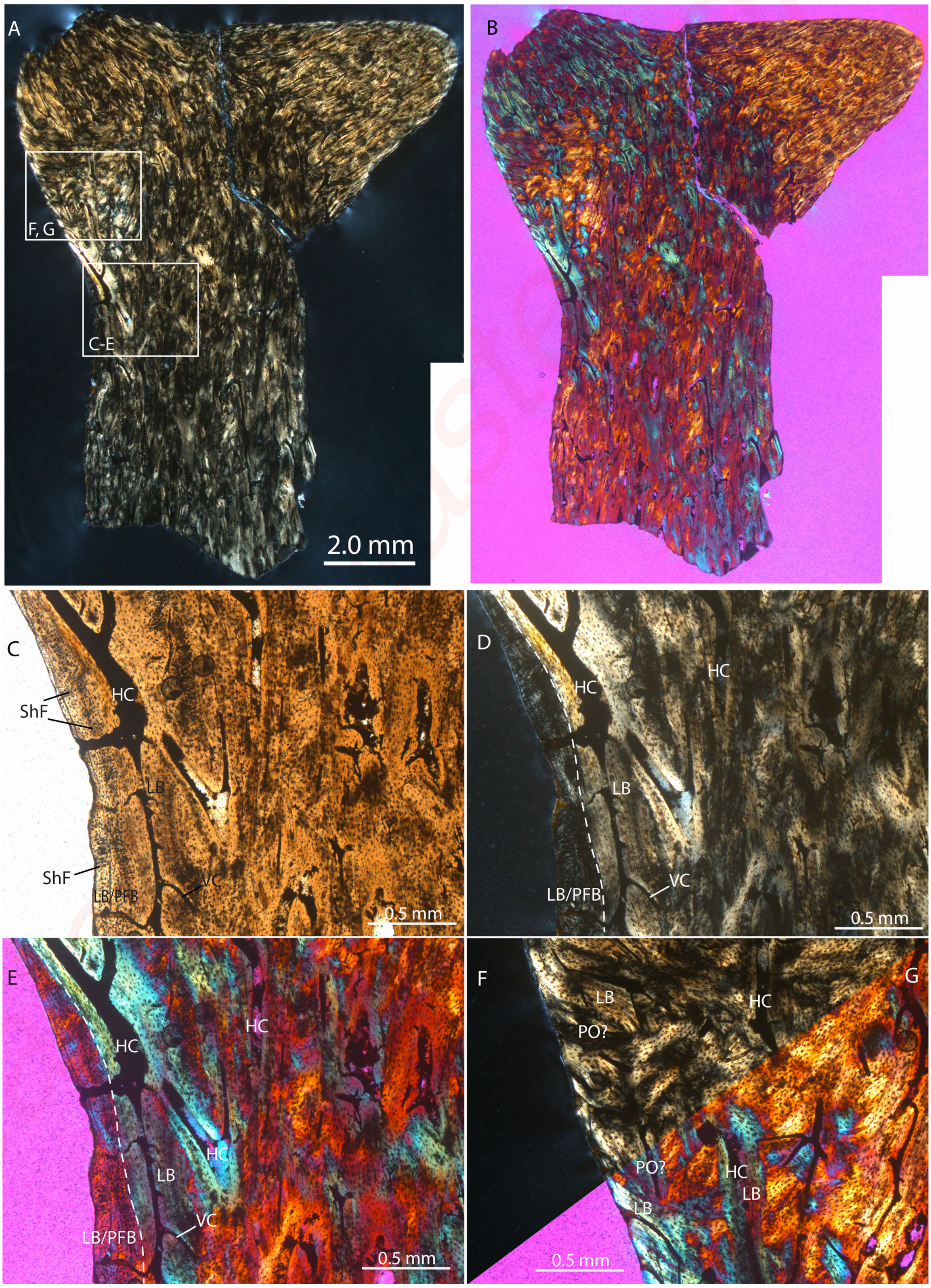

**Online Resource 11:** Detailed histology of a cranial appendage of *Lagomeryx parvulus* (SNSB - BSPG 1959 II 4594) in cross section. Approximate position of the planes of cross-section are indicated in figure 1D. Images A and D in normal transmitted light, images B and E, in cross-polarised light, and images C and F in cross-polarised light using lambda compensator. A-C, Close-up of a partial cross-section of an antler tine tip with more globular cell lacunae without canaliculi. D-F, Close-up of a partial cross-section of the distal part of the pedicle. Note that in both sections, secondary osteons forming dense Haversian bone make up most of the visible bone tissue, whereas patches of primary bone (remnants of primary osteons?) are present only adjacent to the bone surface. Most of the secondary osteons in the distal pedicle portion are oriented longitudinally, whereas they are angled towards the external bone surface in the sectioned tine. Abbreviations: PO, primary osteon; SO, secondary osteon.

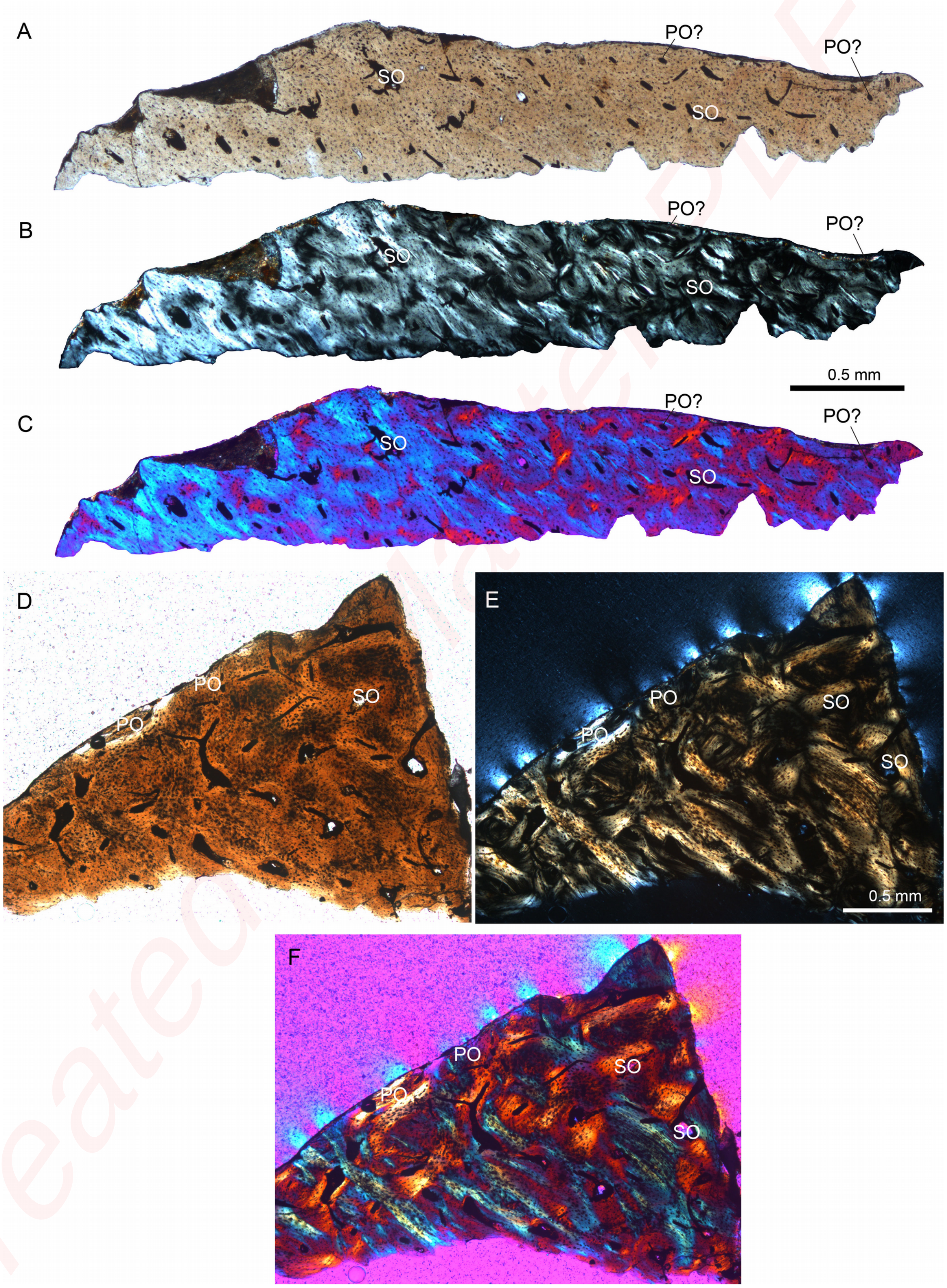


Created with a trial version of PDF Editor

**Online Resource 12:** Detailed histology of antler tines of *Dicrocerus elegans* in cross section (A, B: proximal portion of tine, NMB San.15061; C, D: close-up proximal portion of tine, NMB San.15061; see Online Resource 3 Figure K) and close-up of longitudinal section (E, F: distal portion of tine, NMB San.15062, see Online Resource 3 Figure I). Images C, E in normal transmitted light, image A in cross-polarised light, and images B, D, F in cross-polarised light using lambda compensator. C, D, Well-preserved patch of primary bone composed of fibro-lamellar bone tissue with reticular vascularisation, forming one of the external ridges of the tine.  E, F, Central portion of tine composed mainly of longitudinally arranged secondary osteons (visible by the central Haversian canals and Volkmann's canals branching off). A few small putative erosion cavities are visible. Abbreviations: FLB, fibro-lamellar bone; HC, Haversian canal of secondary osteon; VC, Volkmann's canal.

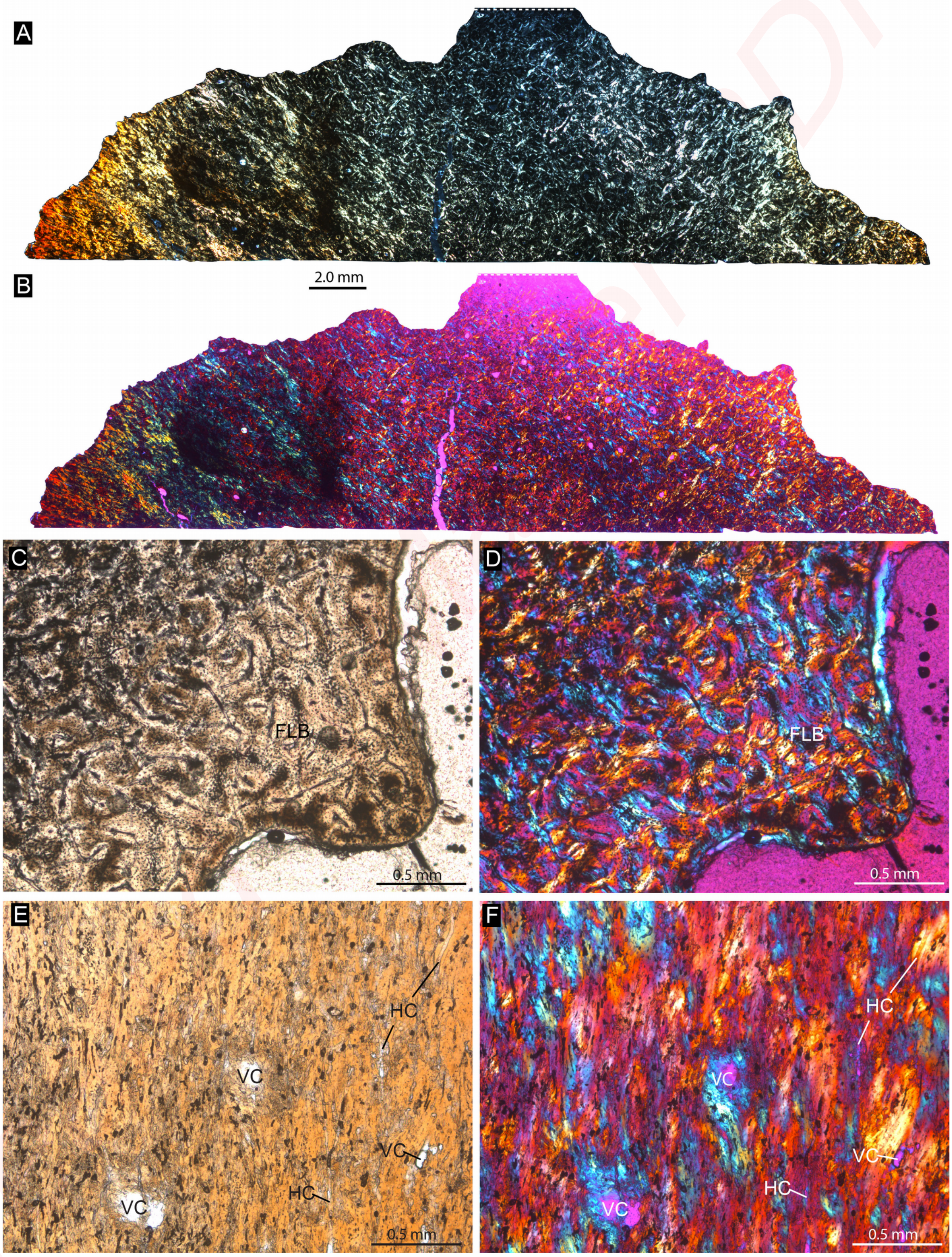

**Online Resource 13:** Detailed histology of an antler of *Paradicrocerus elegantulus* (SNSB - BSPG 1976 VI 24) in longitudinal section. Images in A, B, and D in normal transmitted light, image C in cross-polarised light. A, overview of complete section of antler. B, Close-up of antler bone tissue close to the abscission area. Note Howship's lacunae indicating osteoclast activity. C-F, Close-up of the internal bone tissue at different magnification, consisting of dense secondary osteons and interstitial remnants of primary bone (mostly represented by areas of parallel-fibred bone tissue in which the bone cell lacunae are more globular and more widely spaced). G, Close-up of cortical tissue in the proximal part of the antler, directly adjacent to the abscission area of the cast antler. Note presence of conspicuous Sharpey's fibres in this area. H, I, Close-up of cortical tissue of the distal part of the antler. The bone tissue comprises primary bone tissue (i.e, circumferentially arranged primary osteons, creating a fibrolamellar organisation of the bone tissue) close to the bone surface and secondary osteons with patches of interstitial primary bone tissue in the deeper areas inside the antler. Note absence of coarse or conspicuous Sharpey's fibres here. Abbreviations: HL, Howship's lacunae; LB, lamellar bone; PFB, parallel-fibred bone; PO, primary osteon; ShF, Sharpey's fibres; SO, secondary osteon.

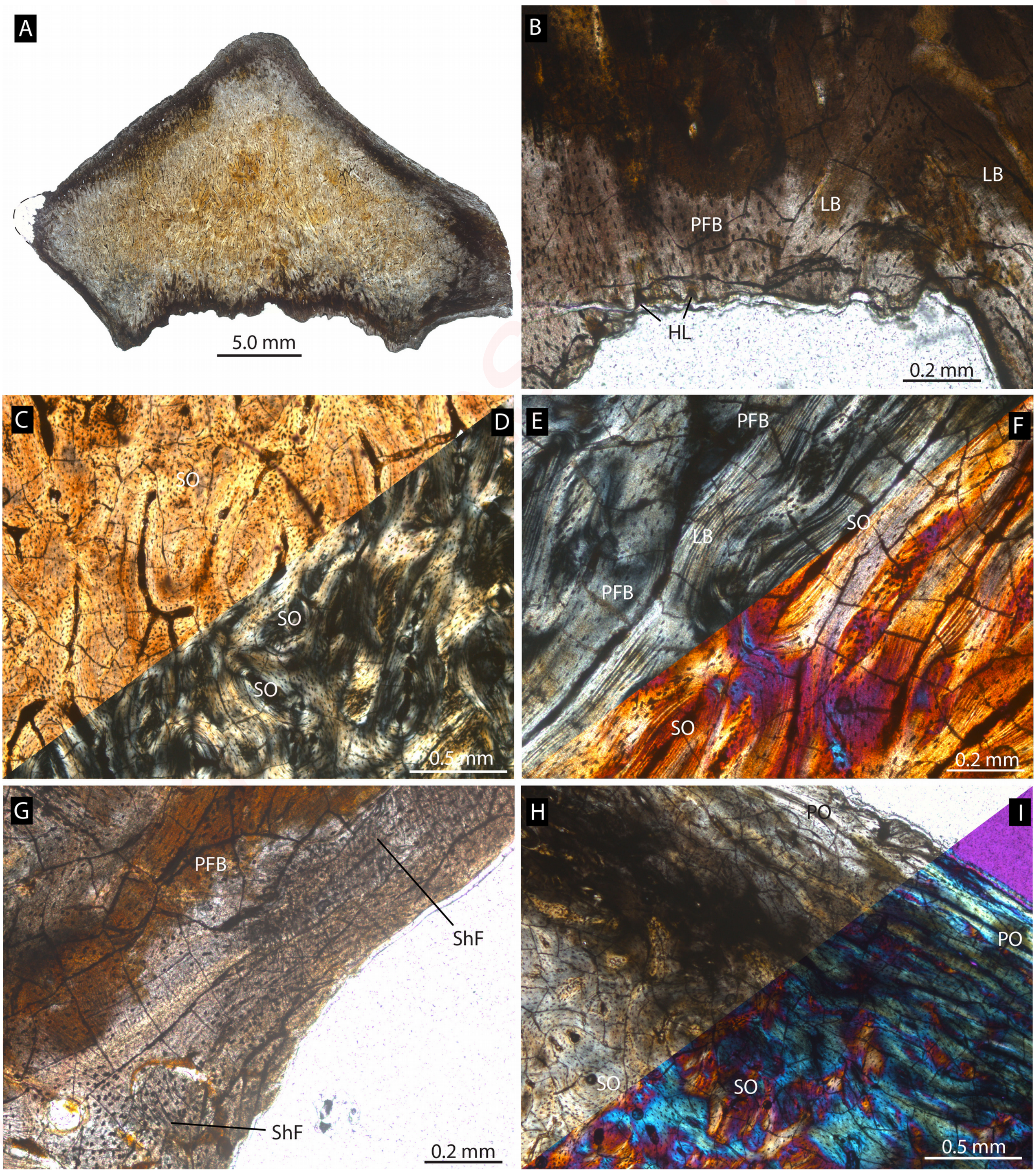

**Online Resource 14:** Radiographic sections of ?*Ligeromeryx praestans*, NMB S.O. 3024, Chitenay (France), Early Miocene (MN3).

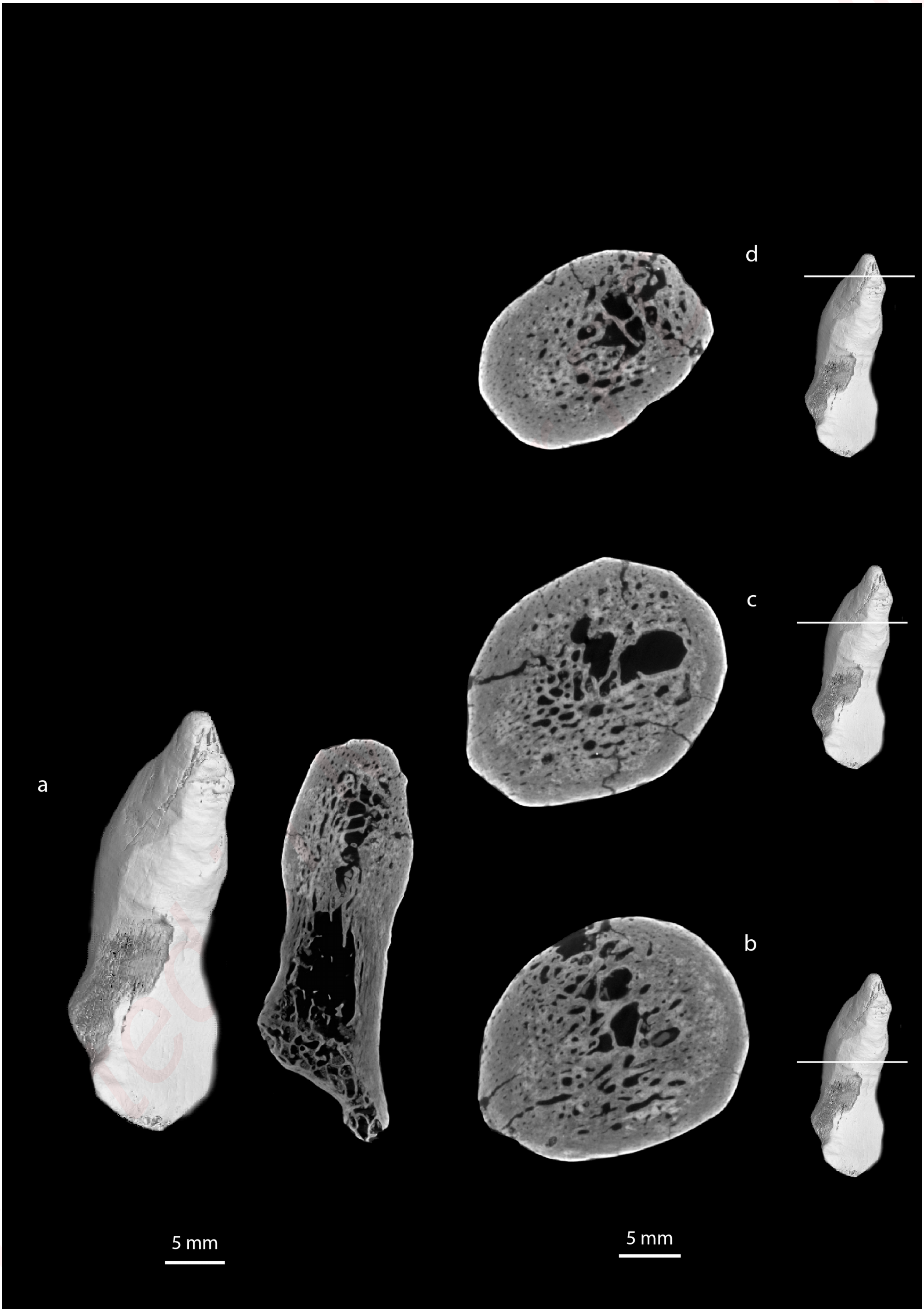

**Online Resource 15:** Radiographic sections of *Acteocemas infans*, holotype, NMB S.O. 3126, Chilleur (France), Early Miocene (MN3).

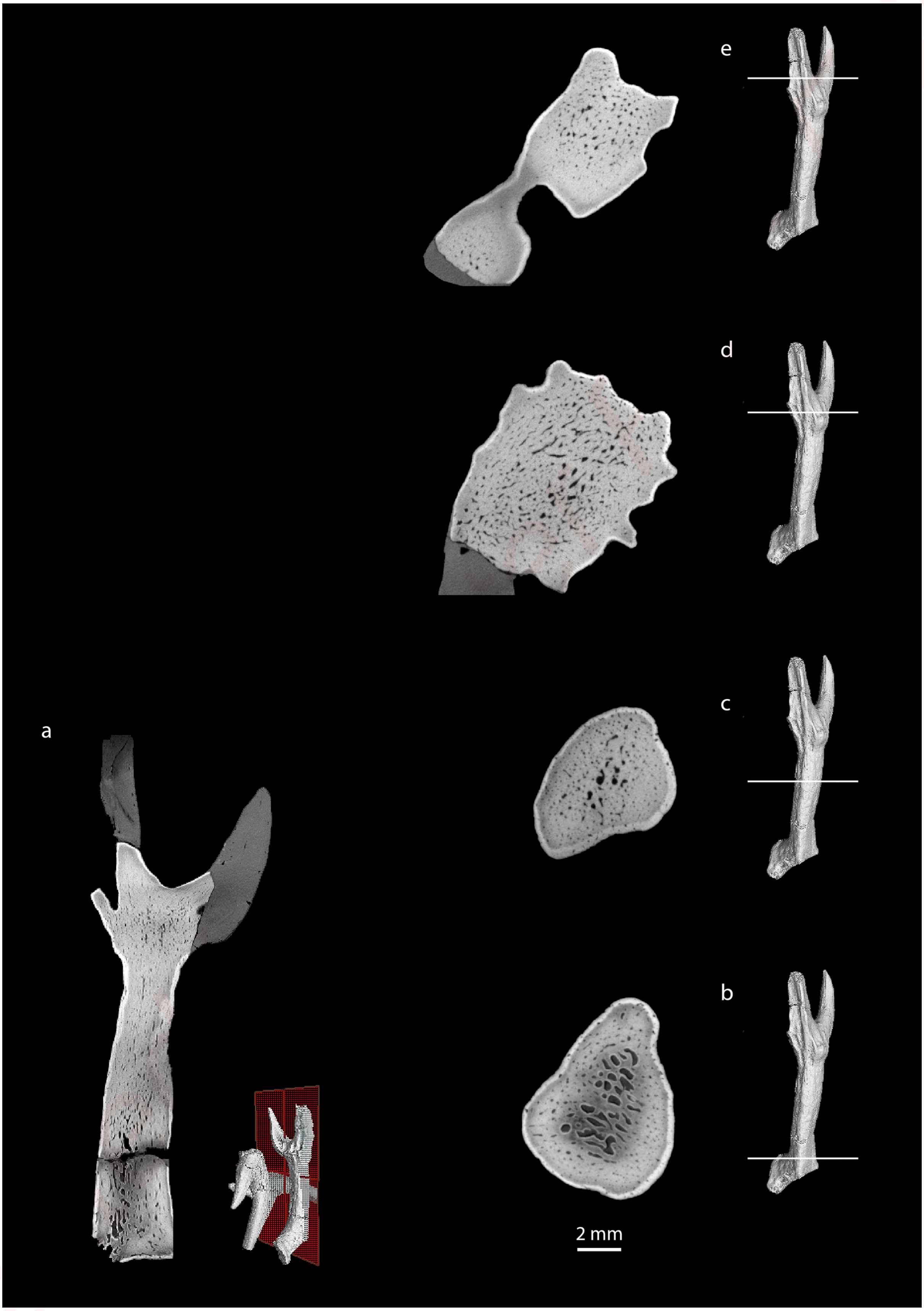

**Online Resource 16**: Radiographic sections of *Ligeromeryx praestans*, paralectotype, NMB S.O. 2078, Chitenay (France), Early Miocene (MN3).

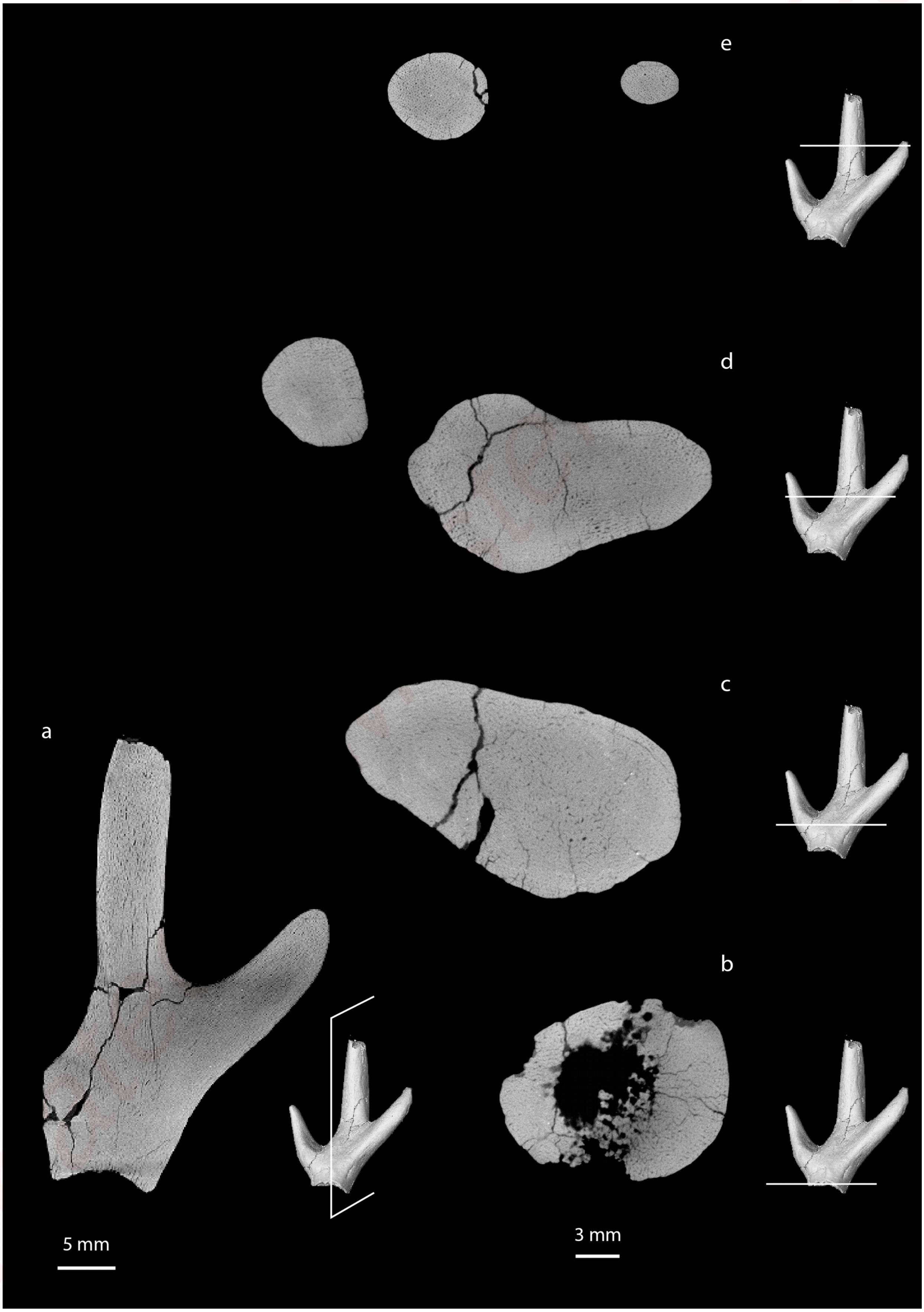

**Online Resource 17**: Radiographic sections of *Ligeromeryx praestans*, paralectotype, NMB S.O. 5720, Chitenay (France), Early Miocene (MN3).

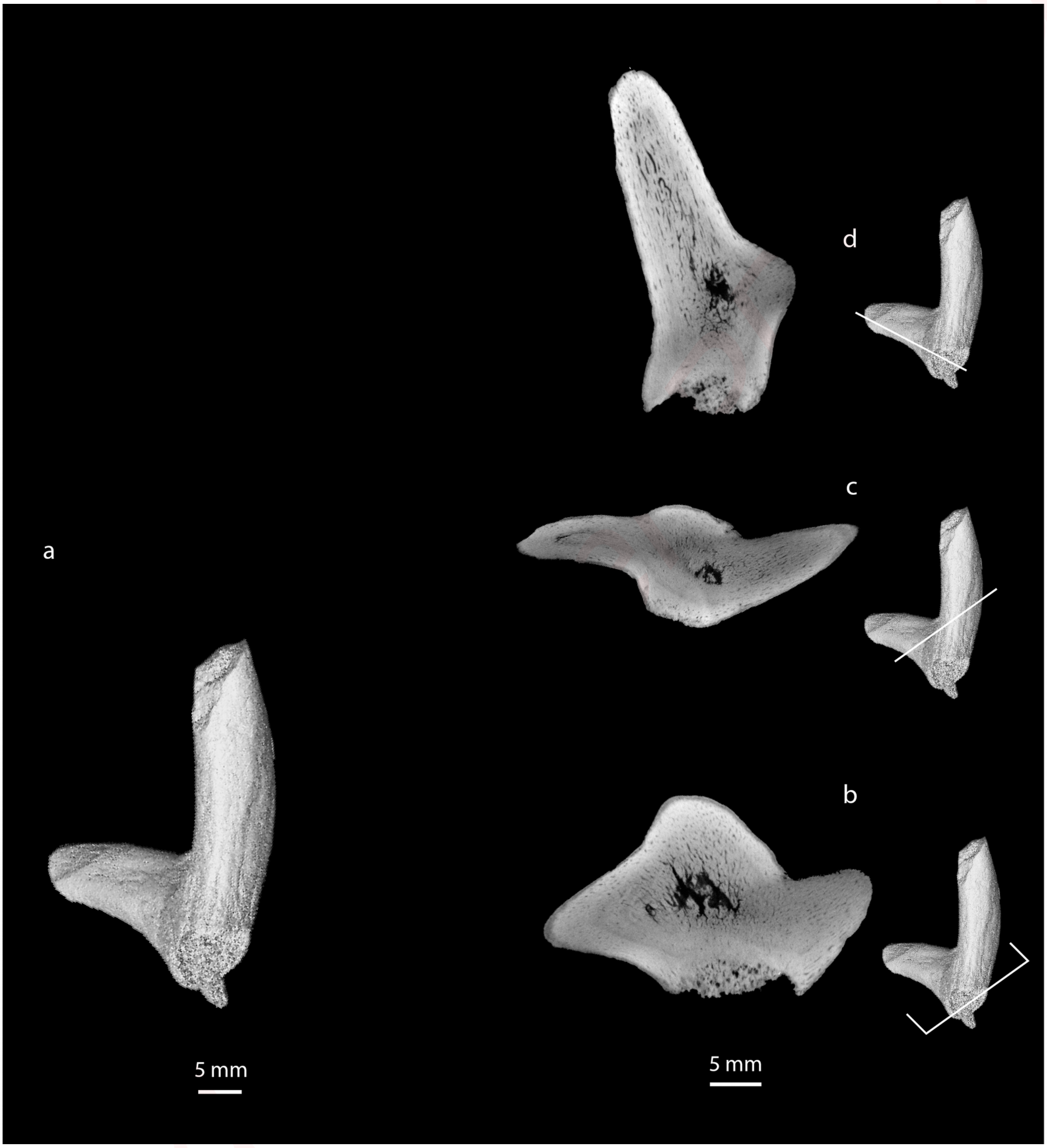

**Online Resource 18**: Radiographic sections of *Ligeromeryx praestans*, lectotype, NMB S.O. 3020, Chitenay (France), Early Miocene (MN3).

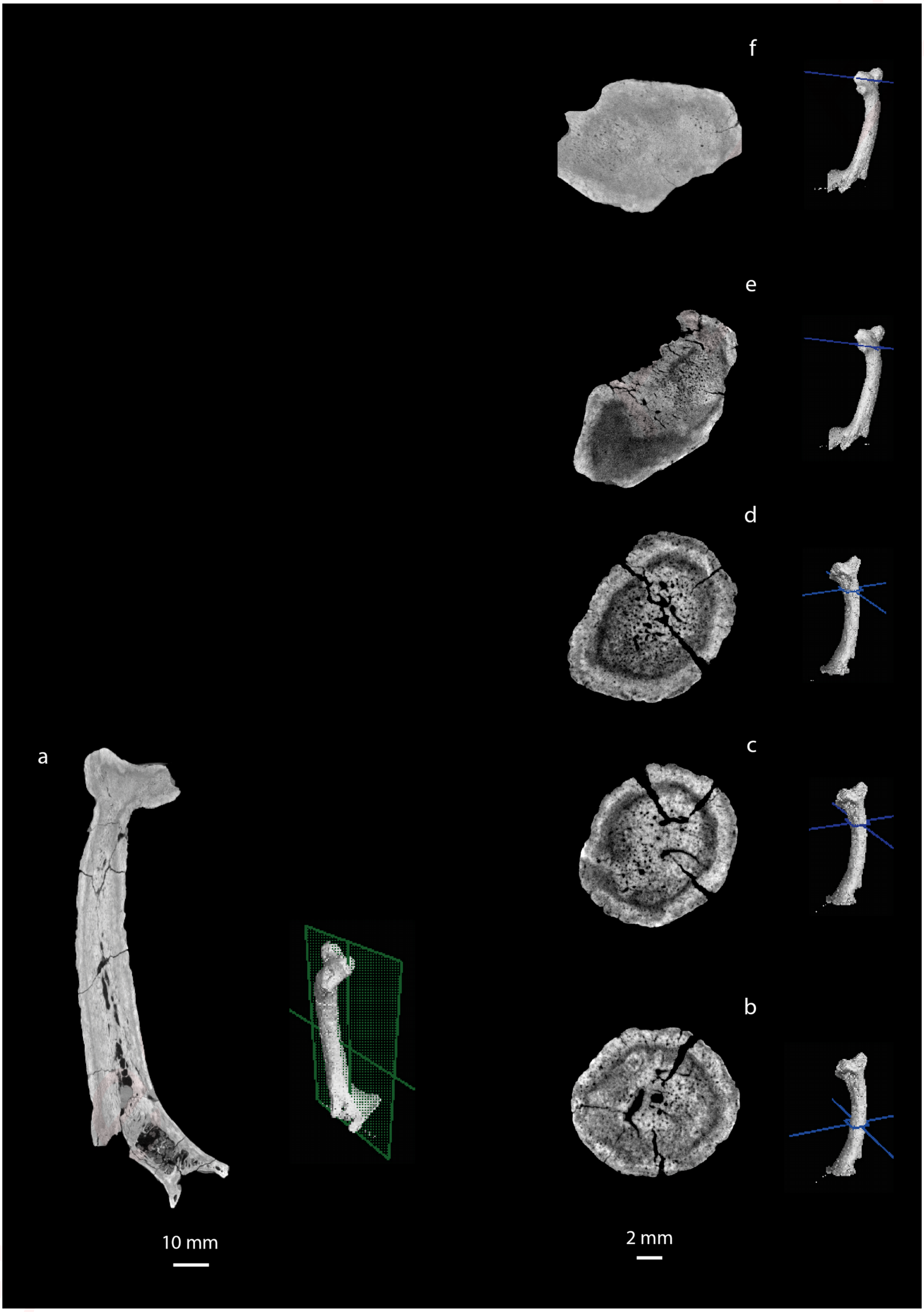

**Online Resource 19**:  Radiographic sections of *Procervulus praelucidus*, SNSB-BSPG 1937 II 16810, Wintershof-West (Germany), Early Miocene (MN3).

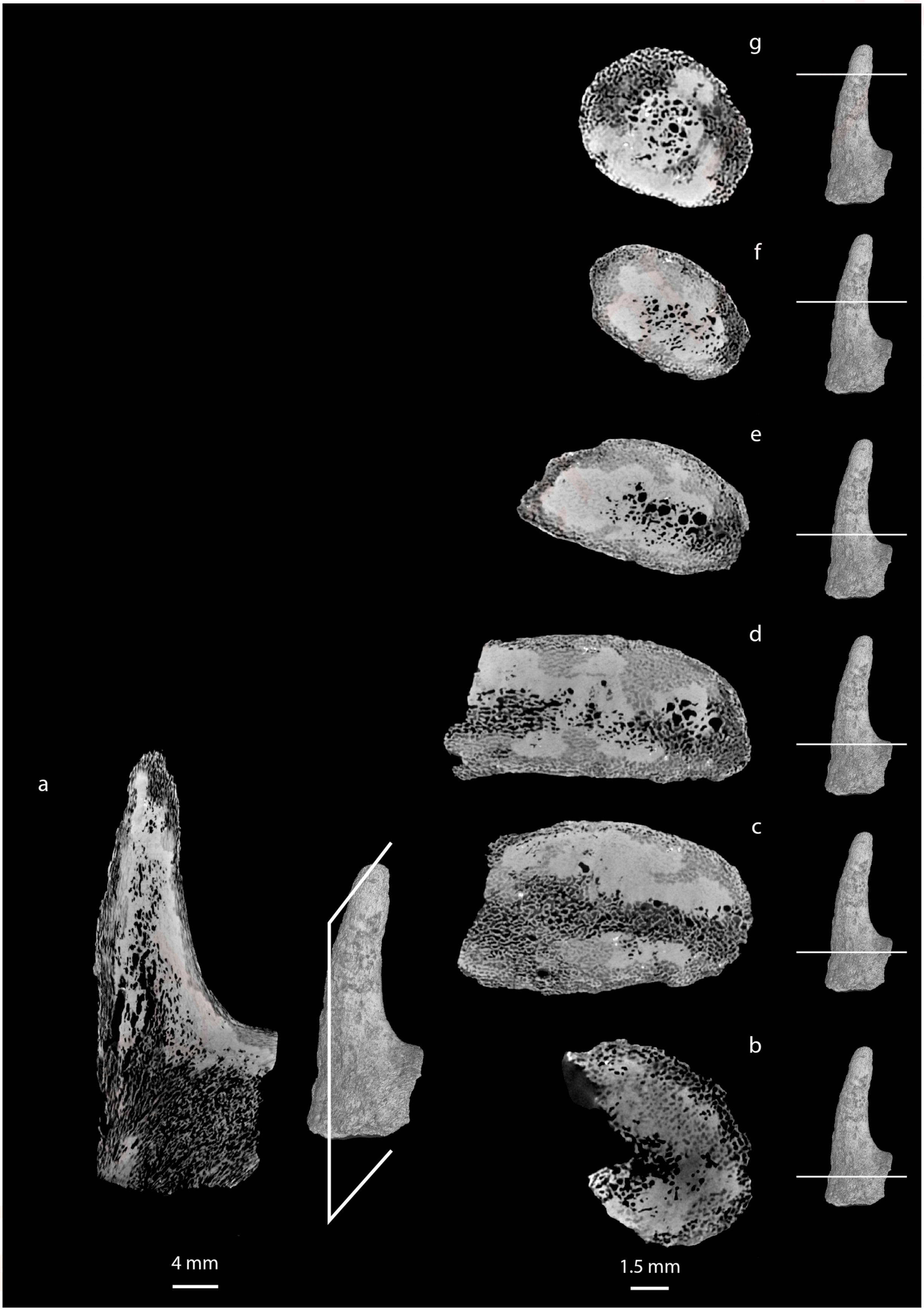

**Online Resource 20**: Radiographic sections of *Procervulus praelucidus*, SNSB-BSPG 1937 II 16841, Wintershof-West (Germany), Early Miocene (MN3).

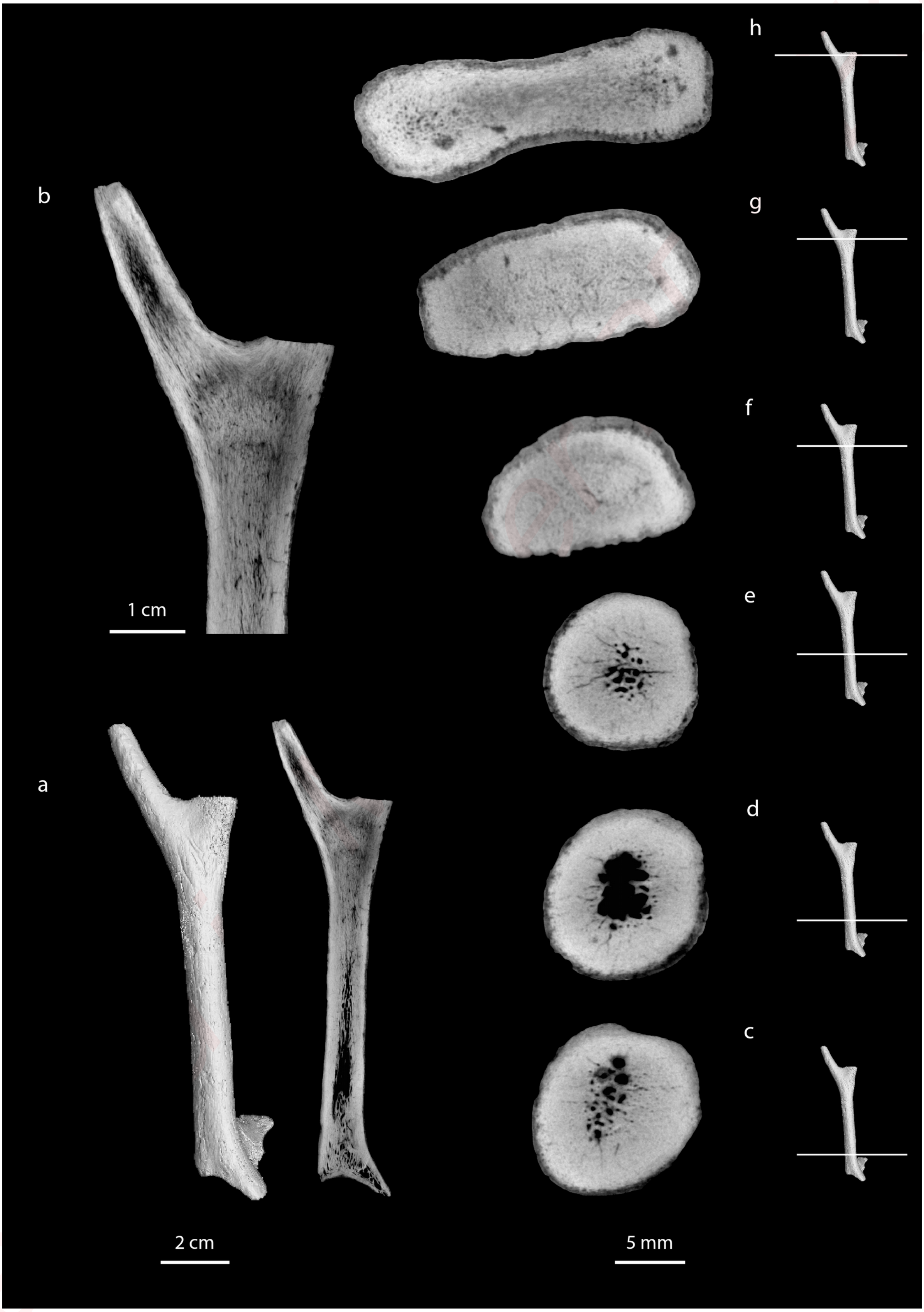

**Online Resource 21**: Radiographic sections of *Procervulus praelucidus*, SNSB-BSPG 1937 II 16842, Wintershof-West (Germany), Early Miocene (MN3).

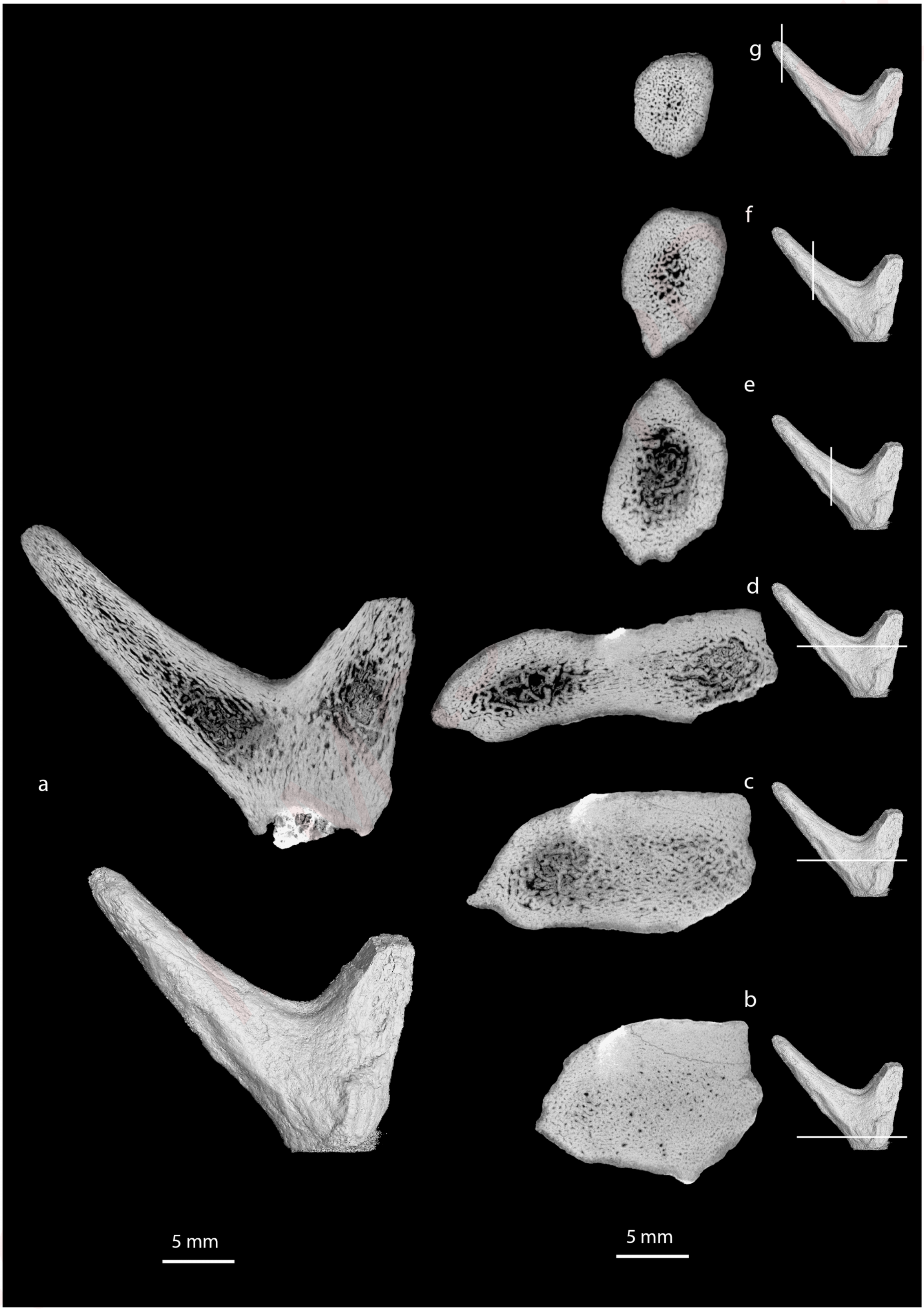

**Online Resource 22**: Radiographic sections of Procervulus praelucidus, SNSB-BSPG 1937 II 16845, Wintershof-West (Germany), Early Miocene (MN3).

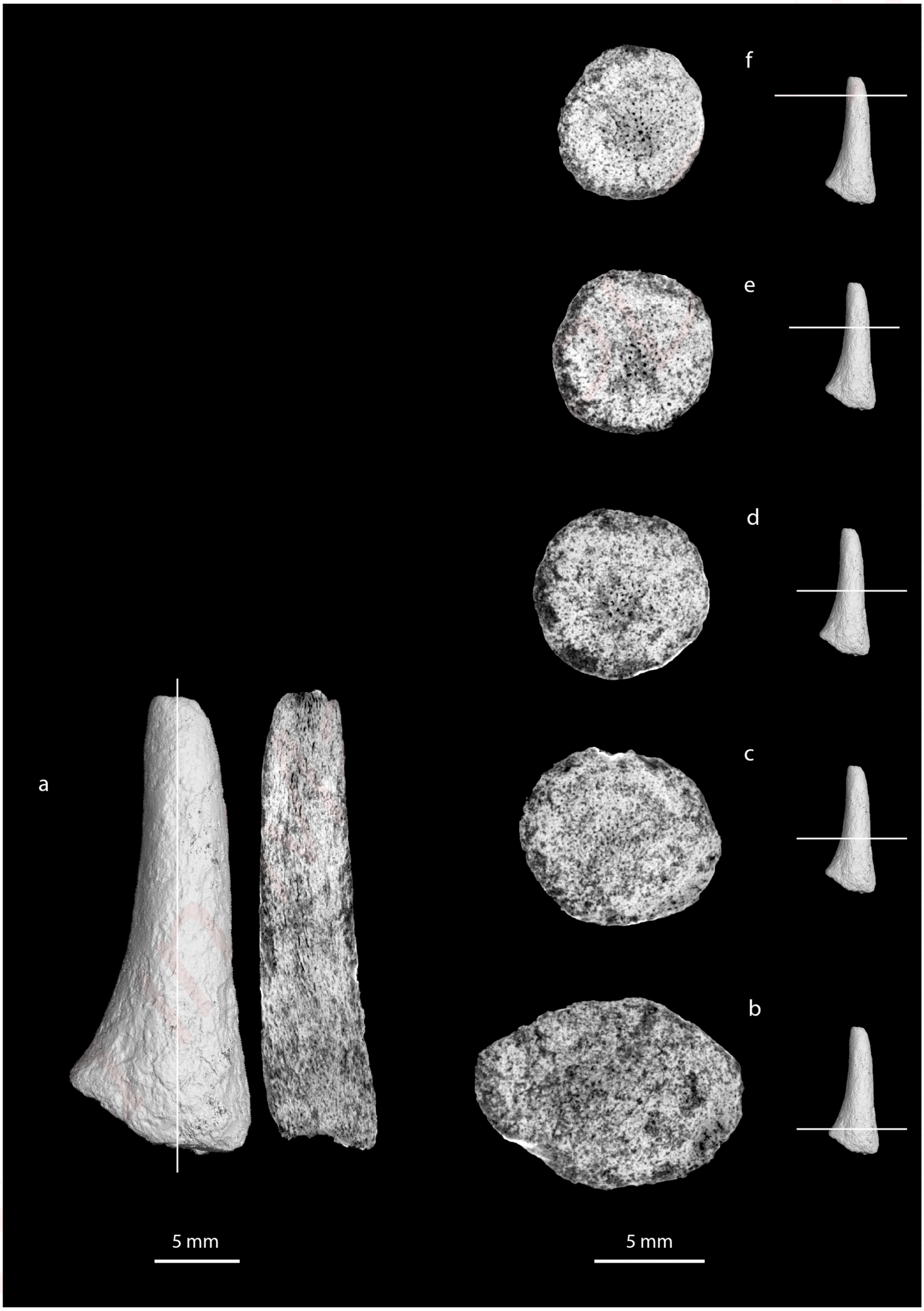

**Online Resource 23**: Radiographic sections of *Lagomeryx ruetimeyeri*, holotype of type species, SNSB-BSPG 1881 IX 55m, Reisensburg (Germany), Early Miocene (MN4).

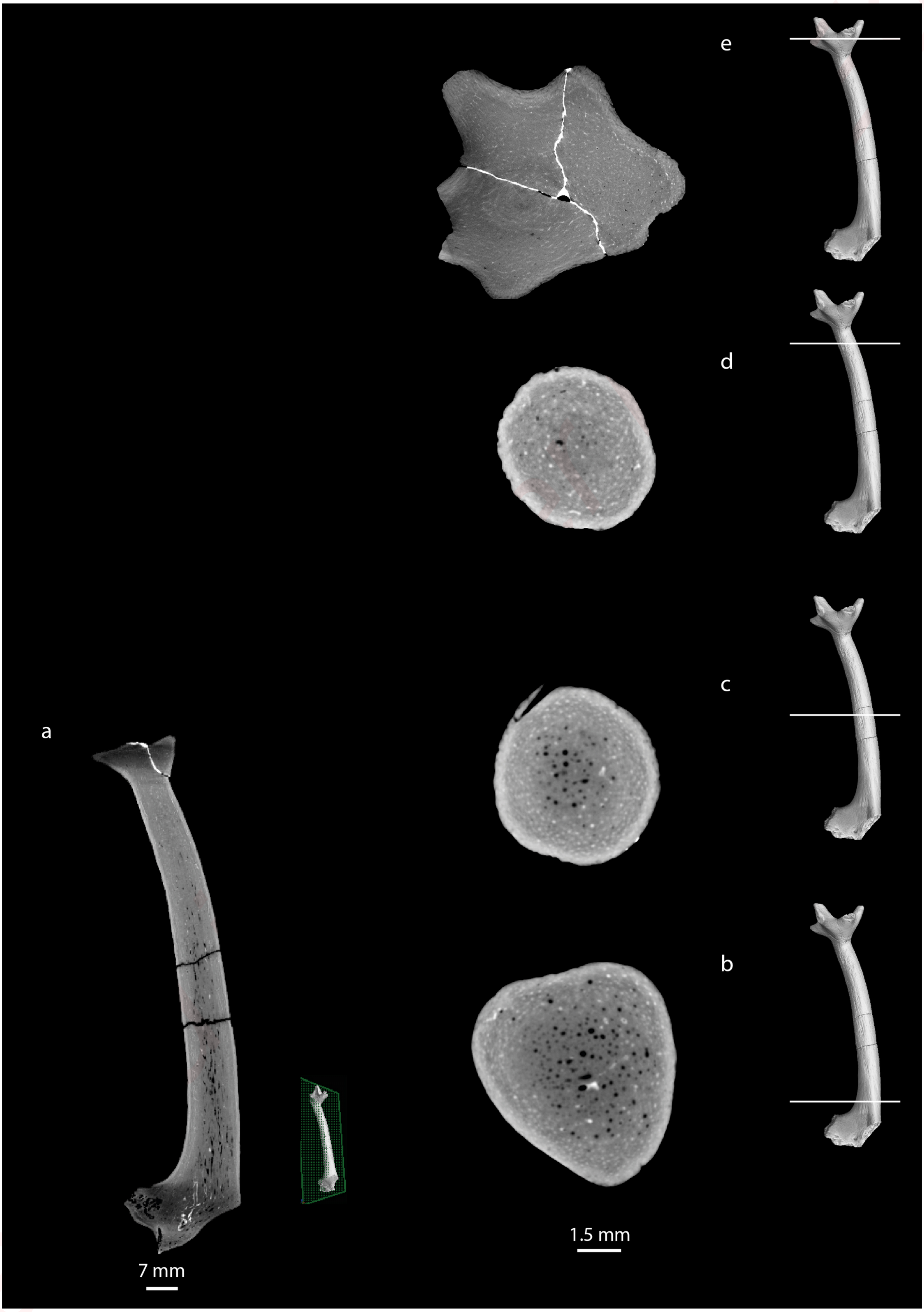

**Online Resource 24**: Radiographic sections of *Procervulus dichotomus*, SNSB-BSPG 1979 XV 555, Rauscheröd (Germany), Early Miocene (MN4).

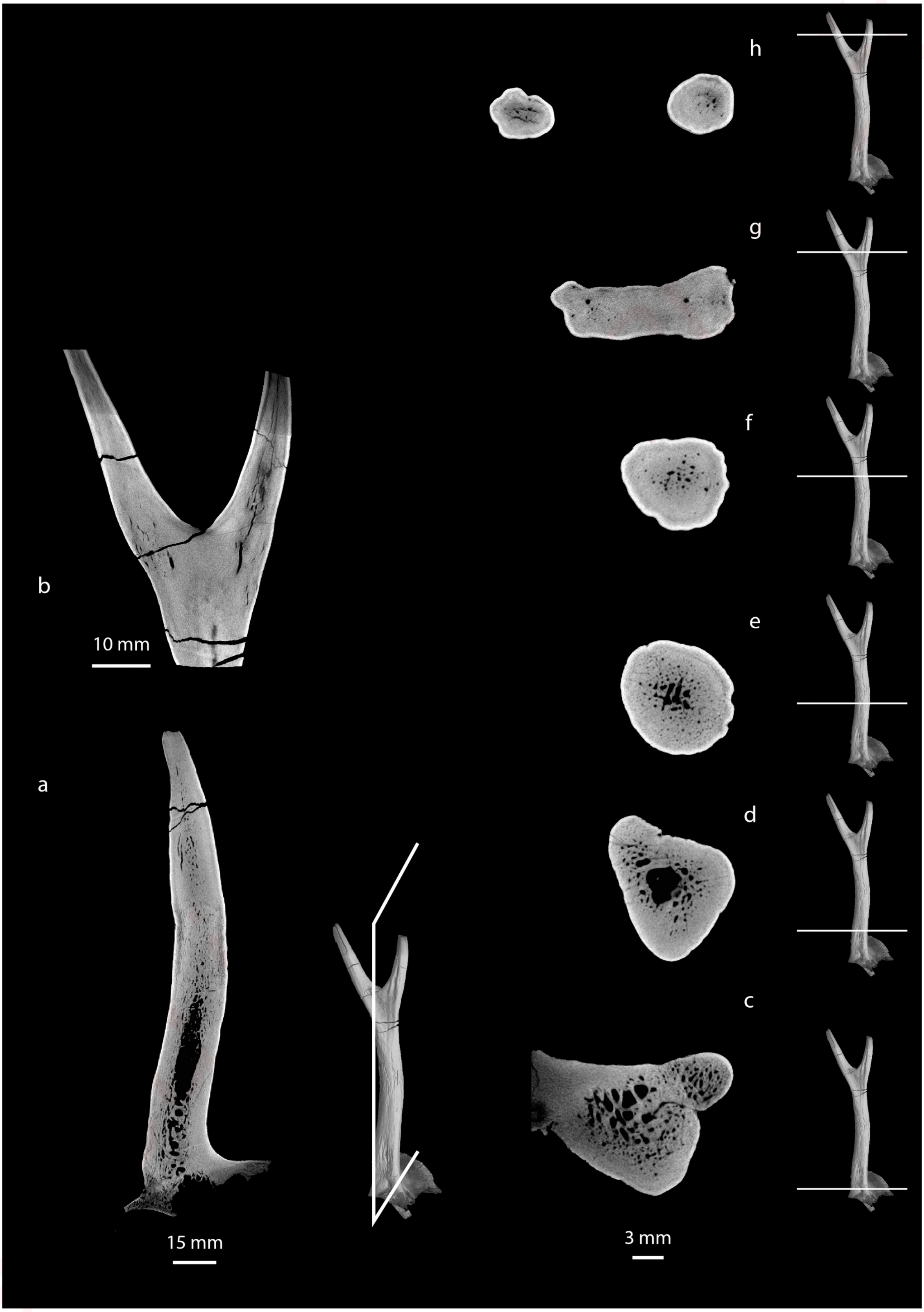

**Online Resource 25**: Radiographic sections of *Procervulus dichotomus*, SNSB-BSPG 1976 XXI 64, Langenau 2 (Germany), Early Miocene (MN4).

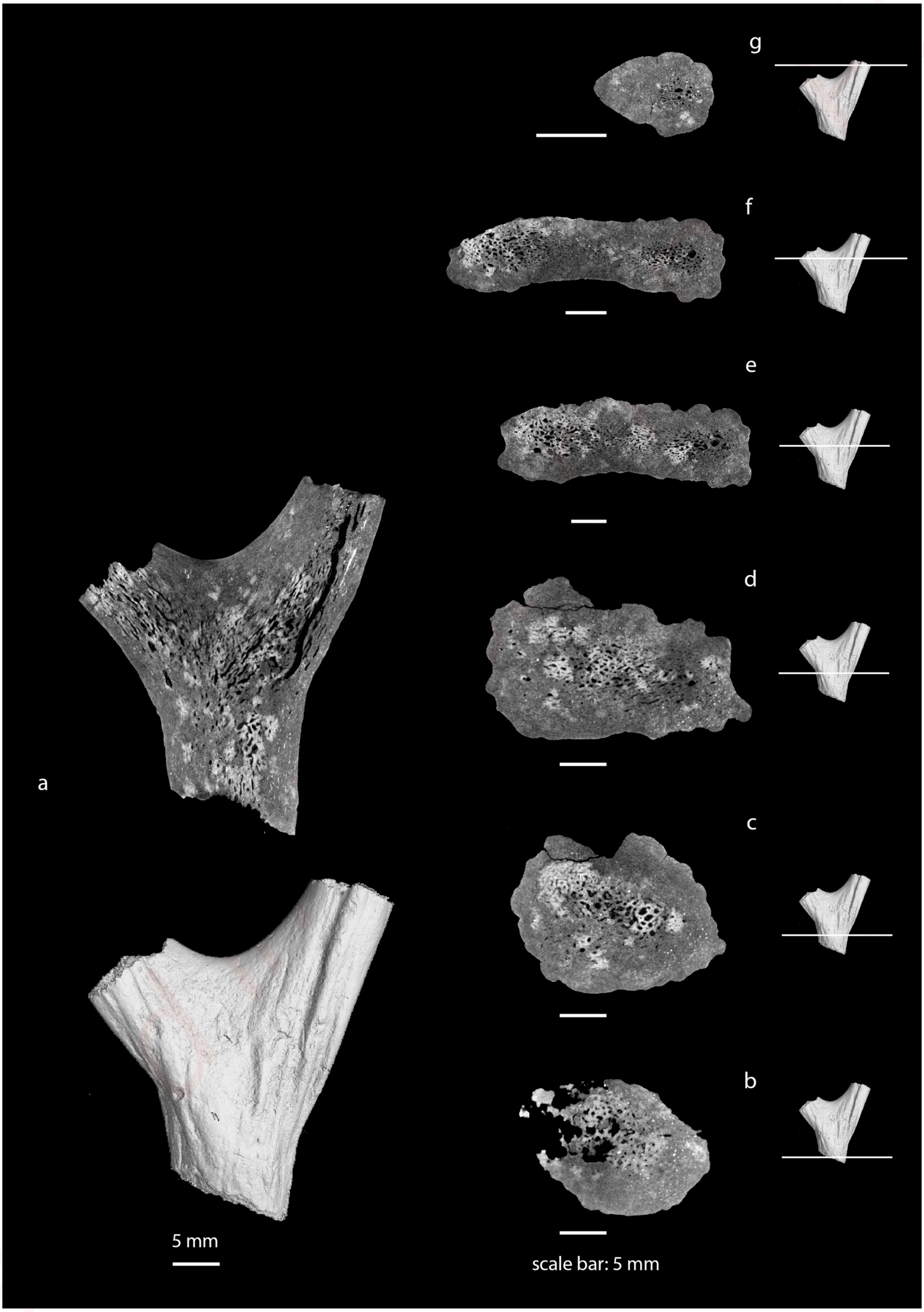

**Online Resource 26**: Radiographic sections of *Heteroprox eggeri*, SNSB-BSPG 1959 II 5268, Sandelzhausen (Germany), Middle Miocene (MN5).

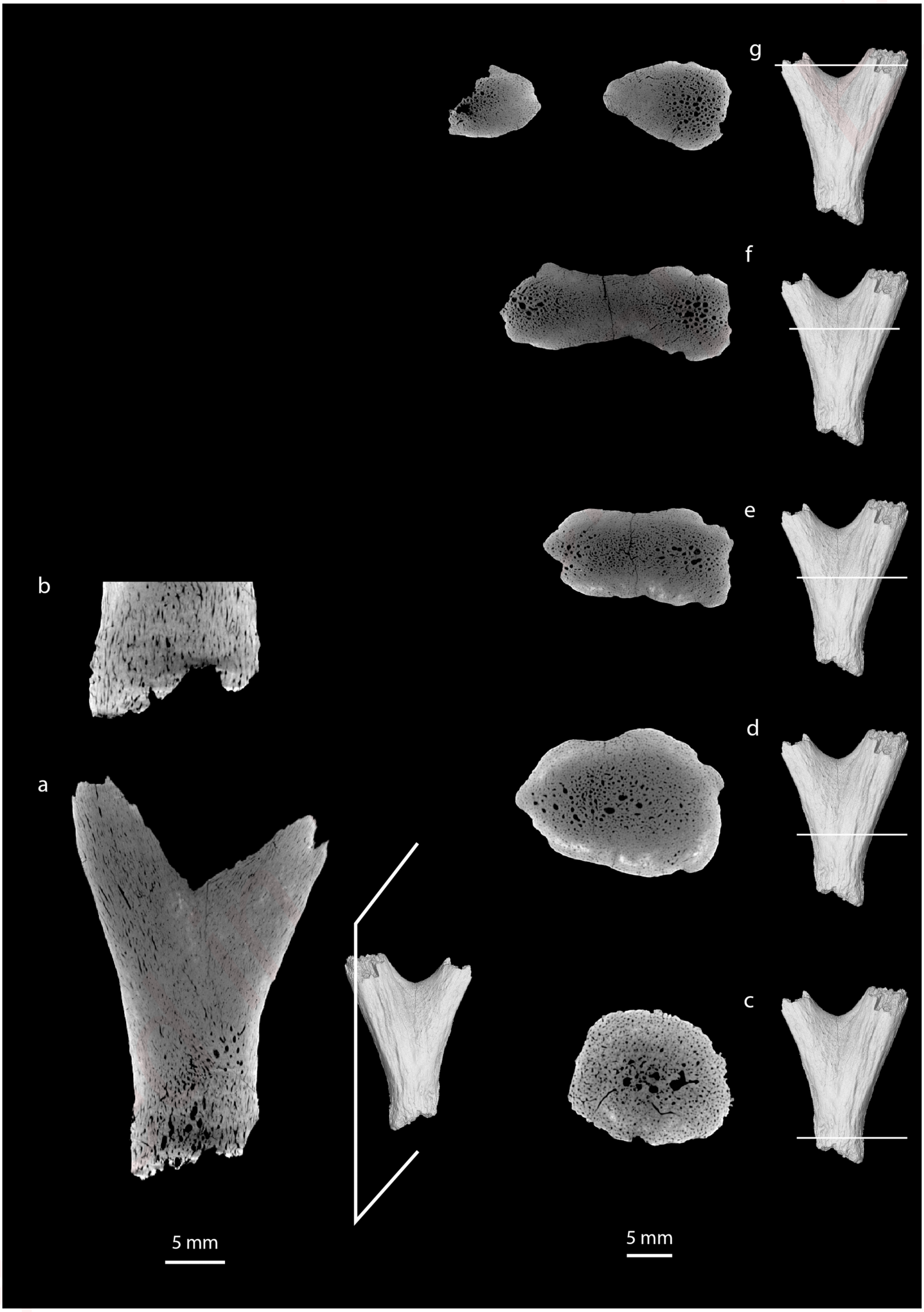

**Online Resource 27**:  Radiographic sections of *Heteroprox eggeri*, SNSB-BSPG 1959 II 5258, Sandelzhausen (Germany), Middle Miocene (MN5).

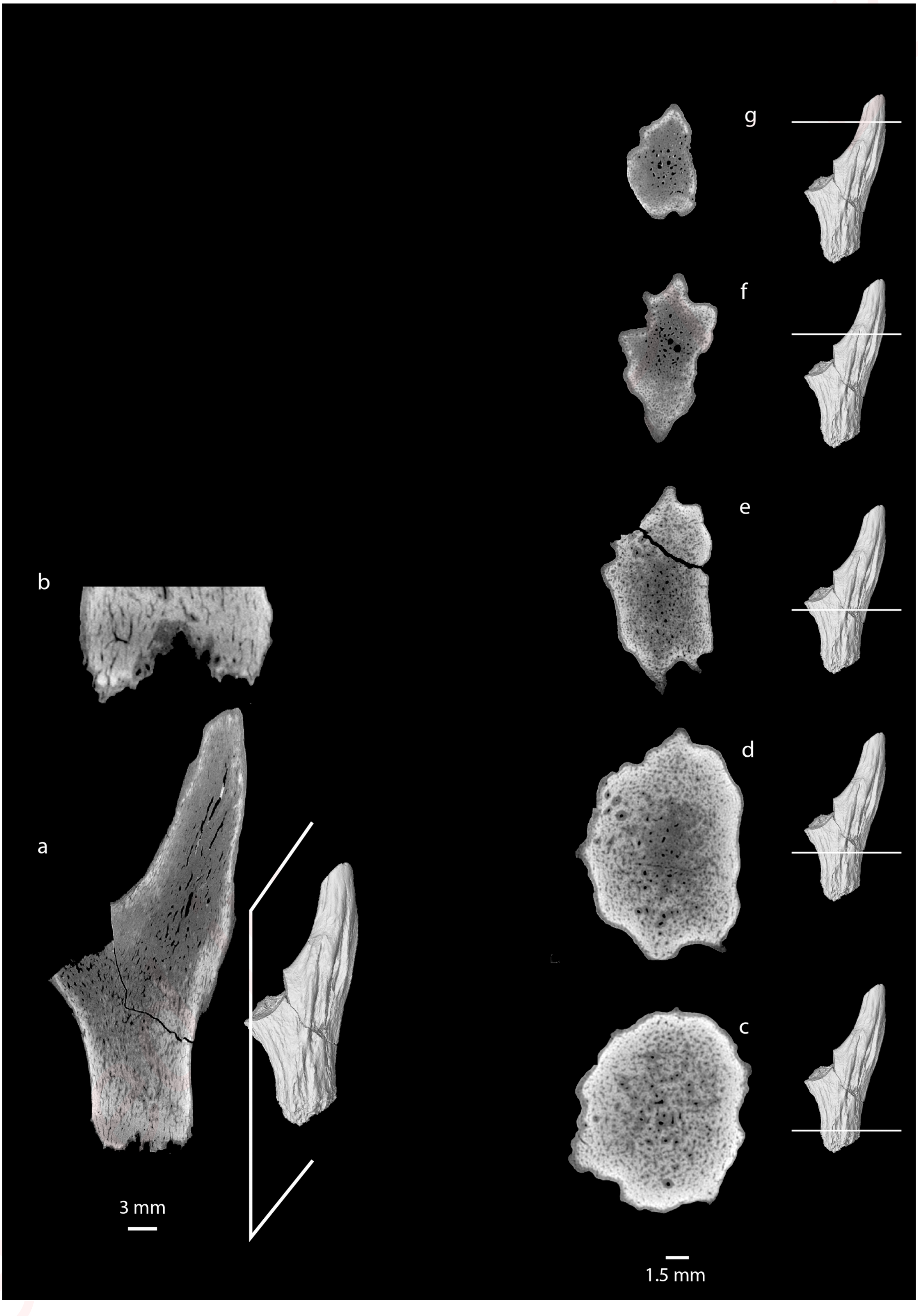

**Online Resource 28**:  Radiographic sections of *Heteroprox eggeri*, SNSB-BSPG 1959 II 5202, Sandelzhausen (Germany), Middle Miocene (MN5).

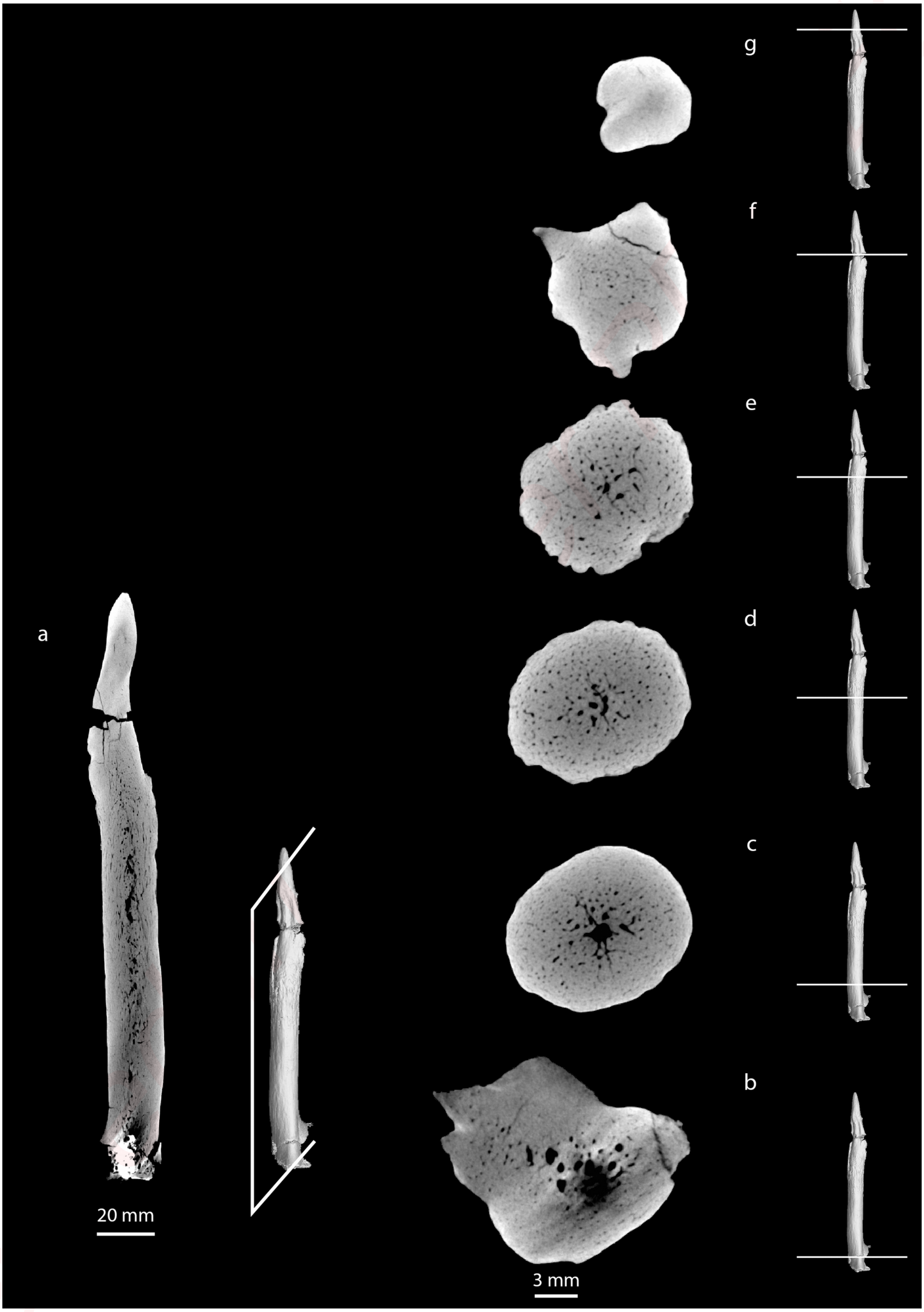

**Online Resource 29**: Radiographic sections of *Heteroprox eggeri*, holotype, SNSB-BSPG 1959 II 5249, Sandelzhausen (Germany), Middle Miocene (MN5).

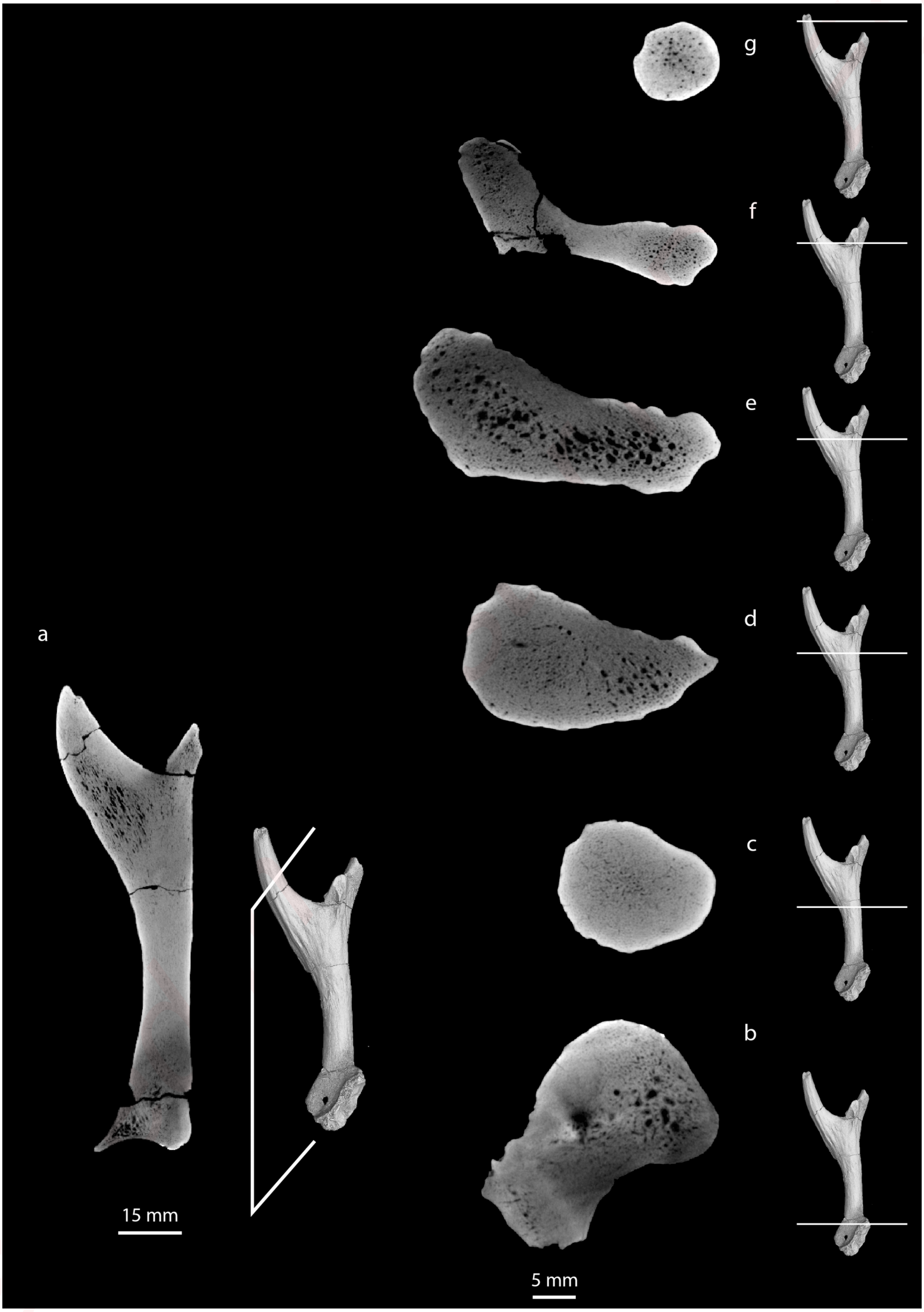

**Online Resource 30**:  Radiographic sections of *Lagomeryx parvulus*, SNSB-BSPG 1959 II 678, Sandelzhausen (Germany), Middle Miocene (MN5).

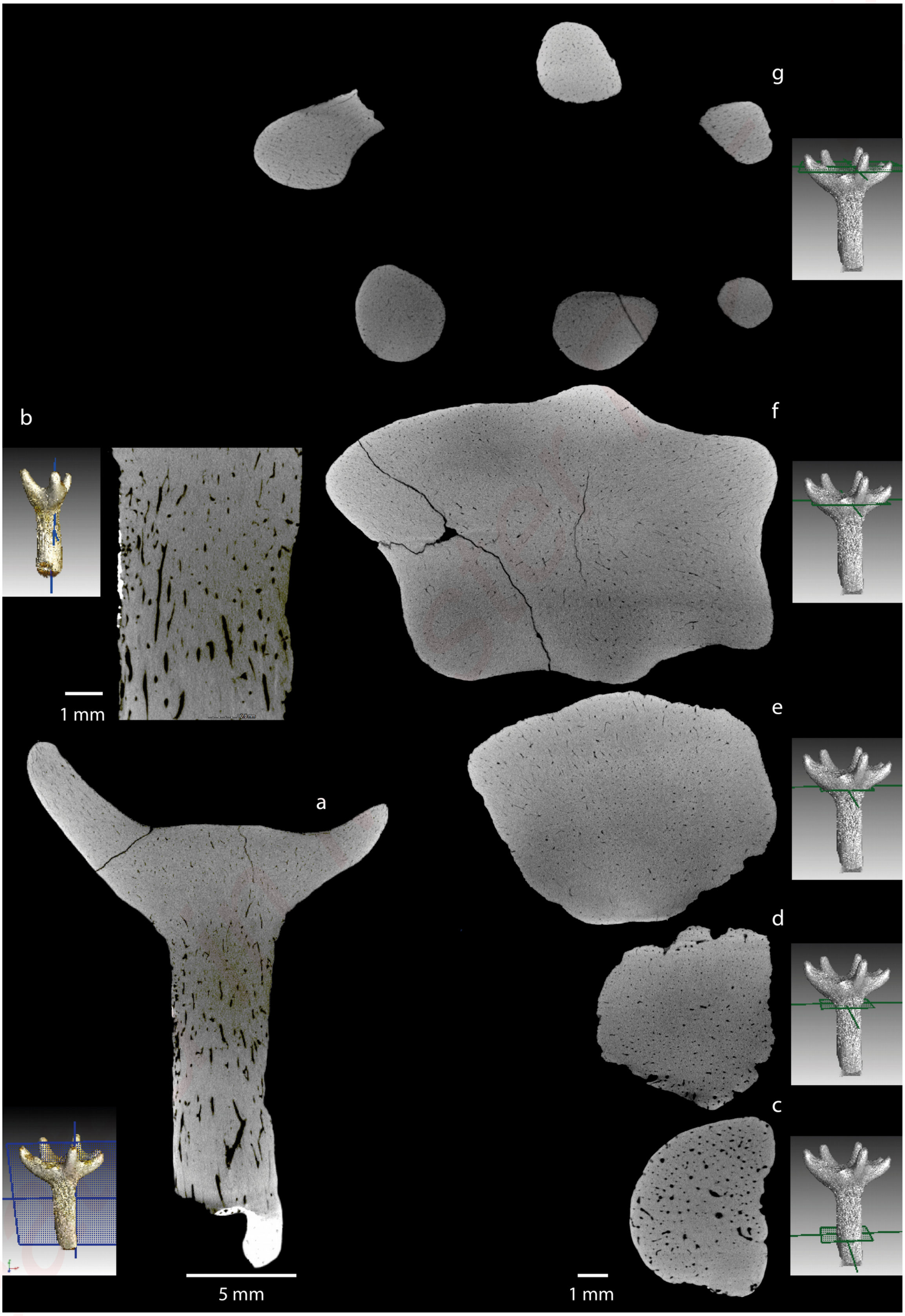

**Online Resource 31**:  Radiographic sections of *Dicrocerus elegans*, SNSB-BSPG 1993 I 35, Sansan (France), Middle Miocene (MN6).

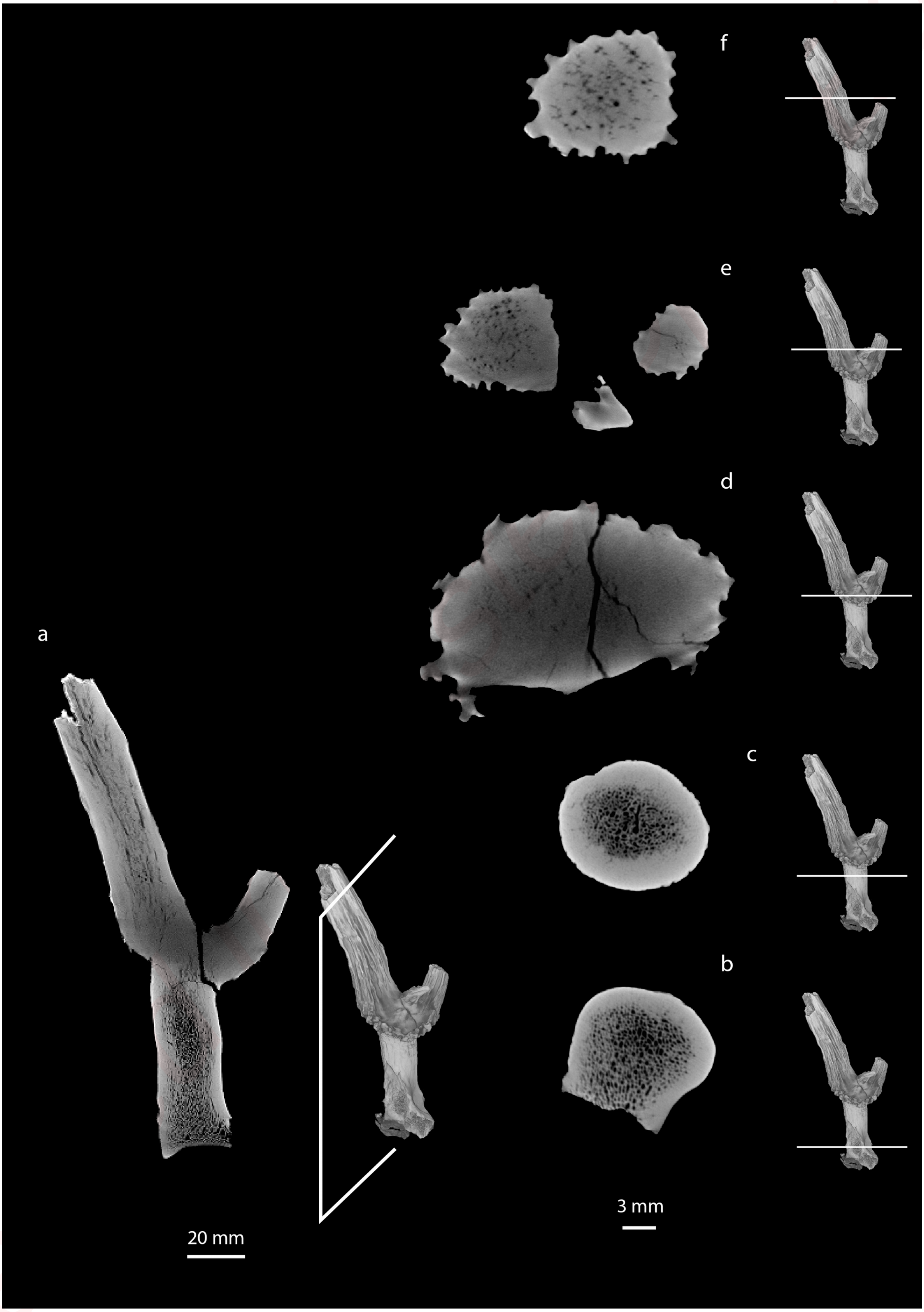


Creditor

**Online Resource 32**: Radiographic sections of *Paradicrocerus elegantulus*, SNSB-BSPG 1993 I 21, Wollersdorf (Germany), Middle Miocene (MN5).

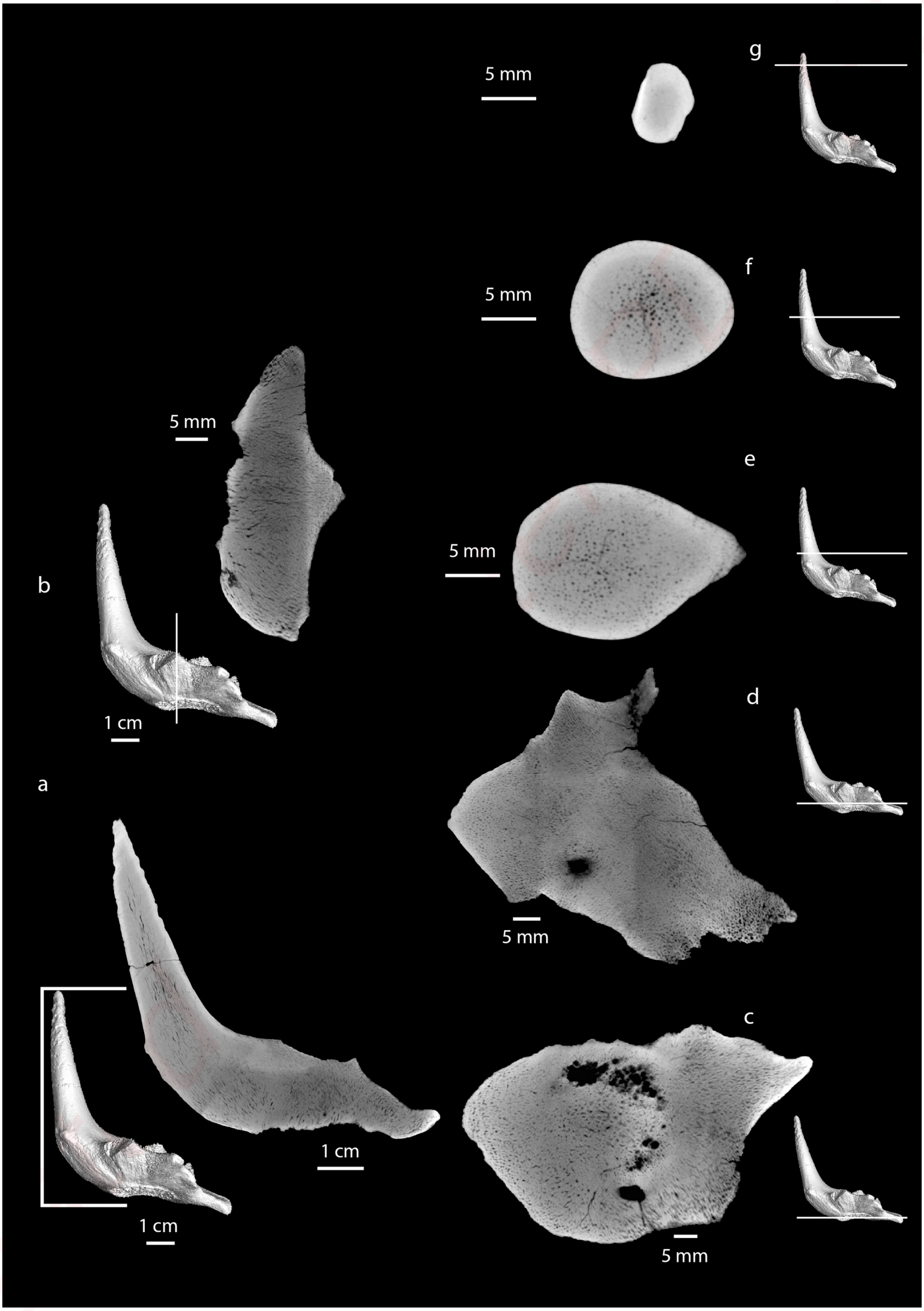

**Online Resource 33**: Radiographic sections of *Paradicrocerus elegantulus*, holotype, NMA 79-5004/761, Stätzling (Germany), Middle Miocene (MN5).

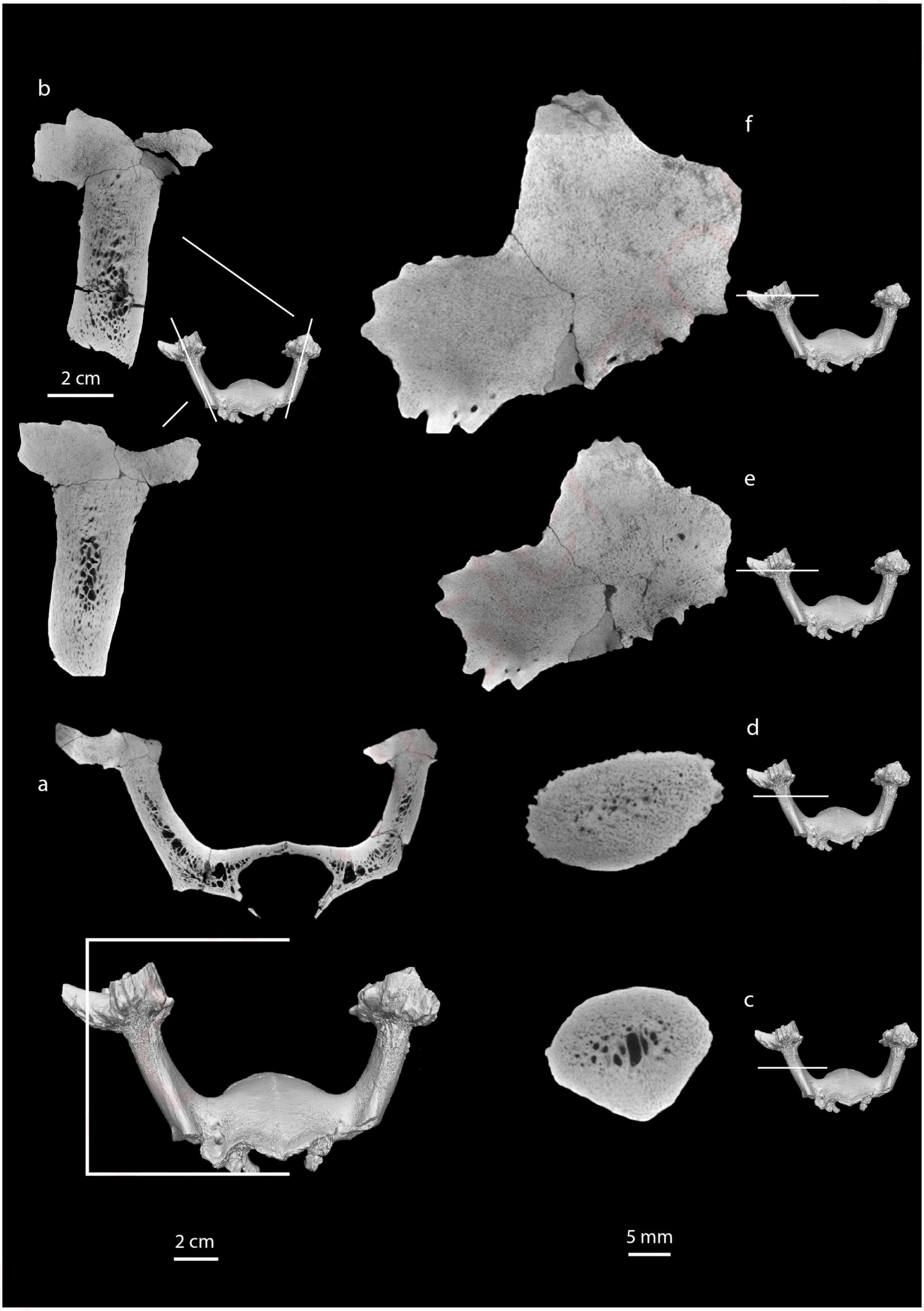

**Online Resource 34**: Radiographic sections of *Heteroprox larteti*, SMNS no number, Steinheim (Germany), Middle Miocene (MN7).

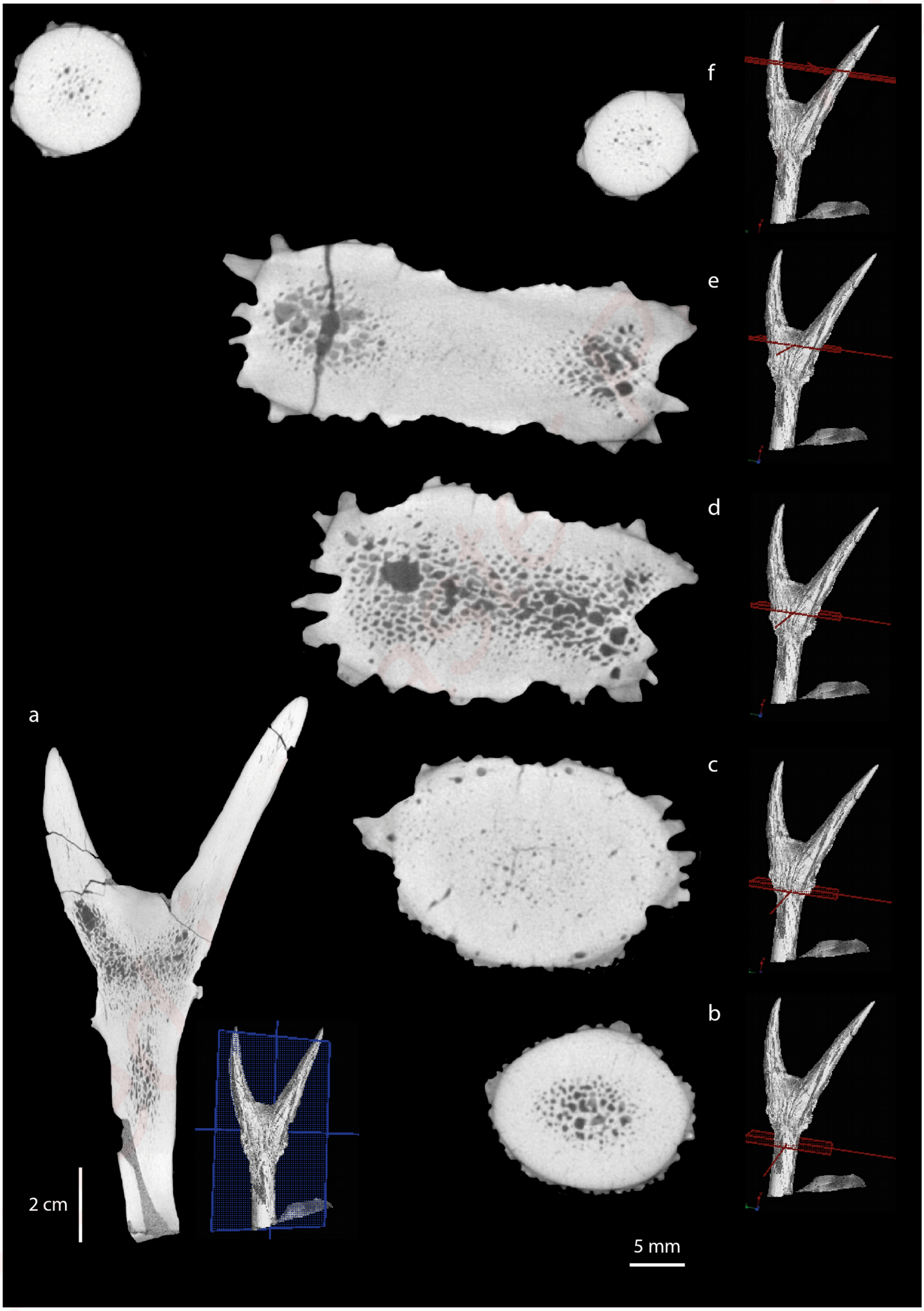

**Online Resource 35**: Radiographic sections of *Euprox furcatus*, SNSB-BSPG 1966 XIV 34, Breitenbrunn (Germany), Middle Miocene (MN8).

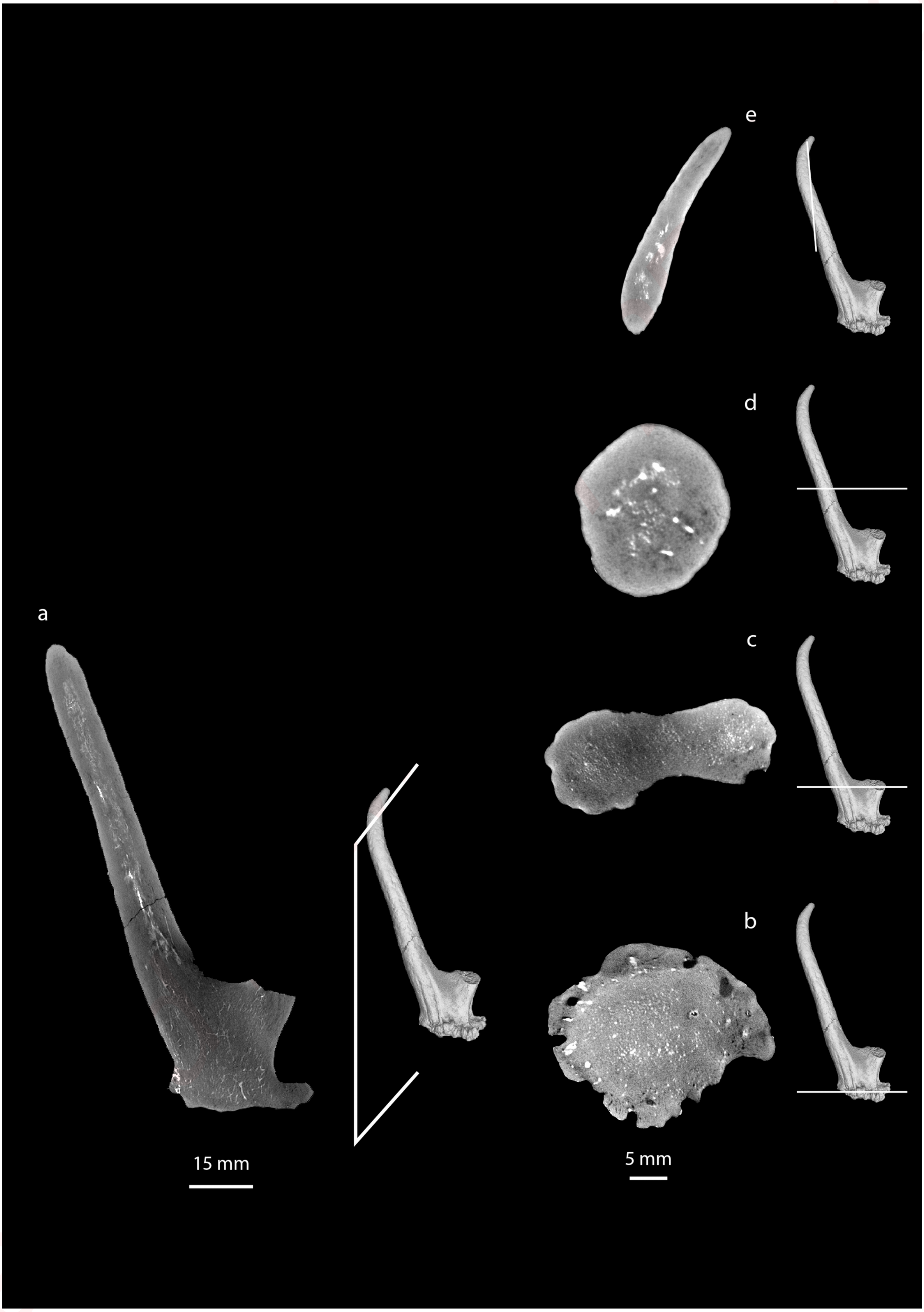


Creditor

**Online Resource 36**: Radiographic sections of *Euprox furcatus*, SNSB-BSPG 1950 I 30, Massenhausen (Germany), Middle Miocene (MN8).

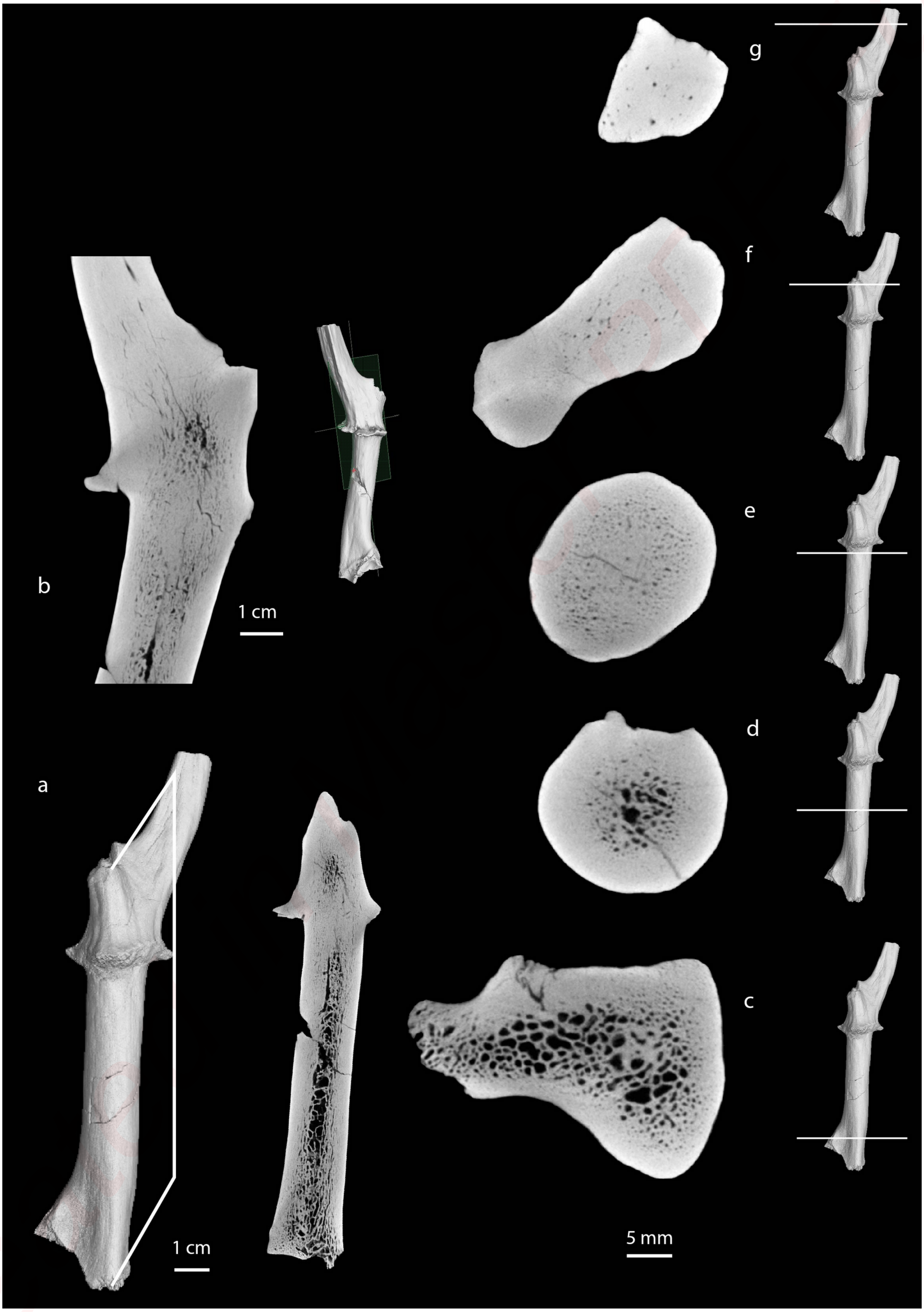

**Online Resource 37**: Radiographic sections of *Muntiacus muntjak*, SNSB-ZSM 1966 237b, Tierpark Hellabrunn München (Germany), Recent.

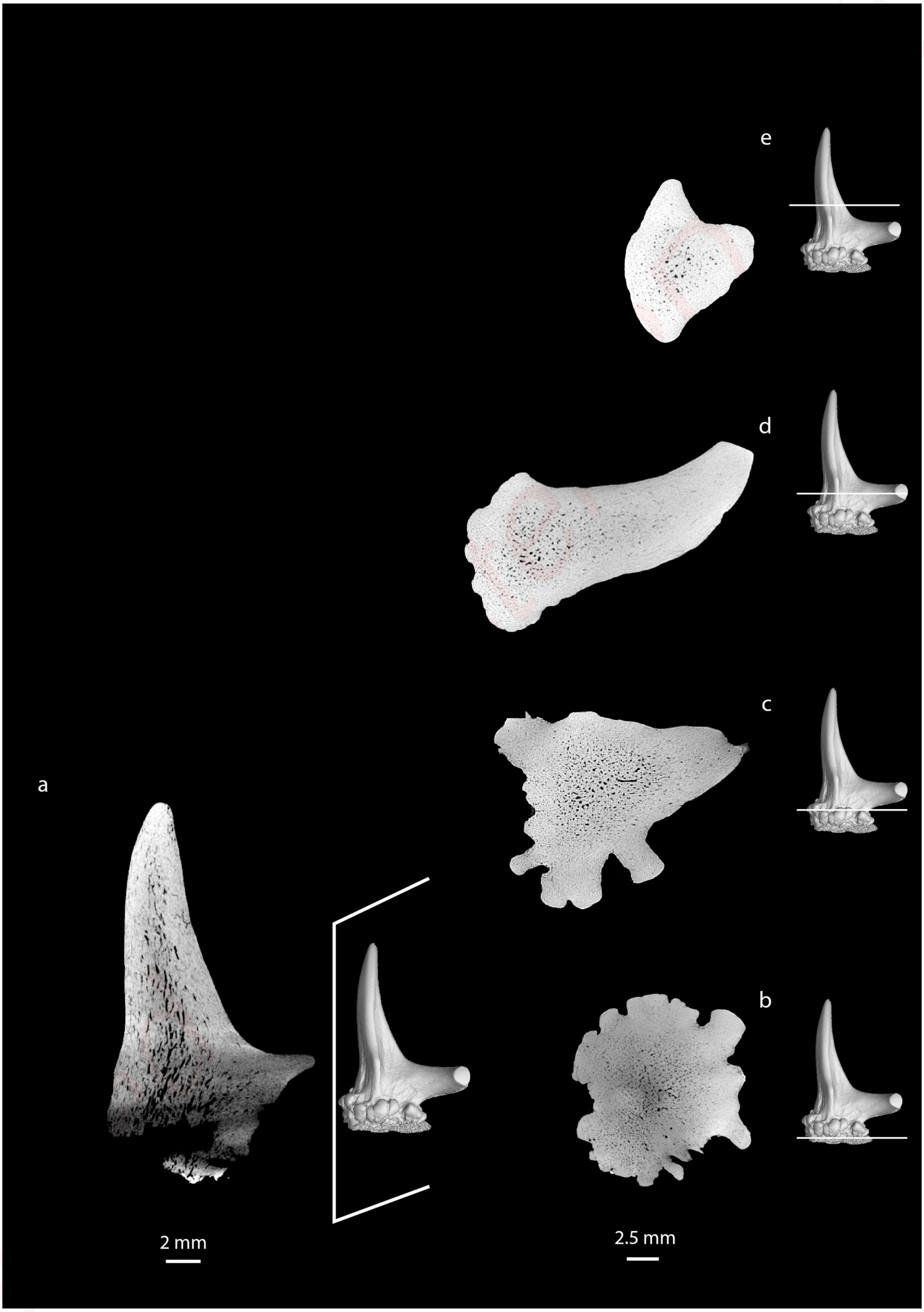

**Online Resource 38**: Radiographic sections of *Procervulus dichotomus*, SMNS 45140, Langenau (Germany), Early Miocene (MN4).

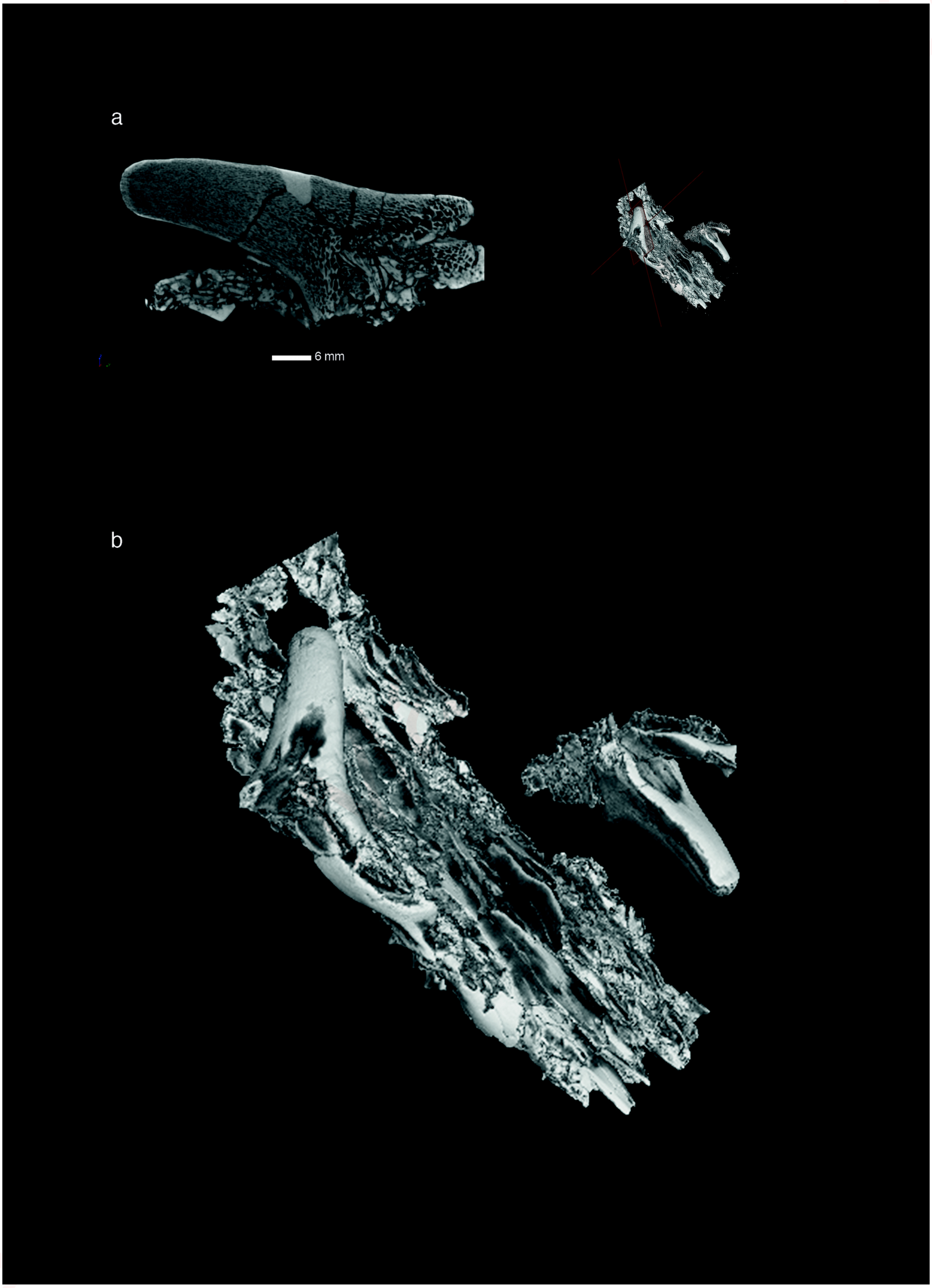
